# Lifting the curse from high dimensional data: Automated projection pursuit clustering for the variety of biological data modalities

**DOI:** 10.1101/2024.04.18.589981

**Authors:** Claire Simpson, Evgeniy Tabatsky, Zainab Rahil, Devon J. Eddins, Sasha Tkachev, Florian Georgescauld, Derek Papalegis, Martin Culka, Tyler Levy, Ivan Gregoretti, Andrei Chernyshev, Hartmut Koeppen, Guenther Walther, Eliver E. B. Ghosn, Darya Orlova

## Abstract

Unsupervised clustering is a powerful machine-learning technique widely used to analyze high-dimensional biological data. It plays a crucial role in uncovering patterns, structure, and inherent relationships within complex datasets without relying on predefined labels. In the context of biology, high-dimensional data may include transcriptomics, proteomics, and a variety of single-cell omics data. Most existing clustering algorithms operate directly in the high-dimensional space, and their performance may be negatively affected by the phenomenon known as the curse of dimensionality. Here, we show an alternative clustering approach that alleviates the curse by sequentially projecting high-dimensional data into a low-dimensional representation. We validated the effectiveness of our approach, named APP, across various biological data modalities, including flow and mass cytometry data, scRNA-seq, multiplex imaging data, and T-cell receptor repertoire data. APP efficiently recapitulated experimentally validated cell-type definitions and revealed new biologically meaningful patterns.

## Introduction

Modern biological data can be quite complex and high-dimensional, making it challenging to uncover meaningful insights from the data. Clustering is often employed to discover interesting patterns in the data by partitioning it into groups/clusters where data points within the same cluster are more similar to each other than to those in other clusters. High-dimensional clustering and projection pursuit [Friedman et al., 1974; Friedman et al., 1982; Huber, 1985] both aim to address the problem of discovering patterns in the data. However, they approach the problem from different angles.

High-dimensional clustering aims to group similar data points together based on a similarity measure or fit to a posited generative model directly in the high-dimensional space. It retains the original data’s full information, but could suffer from the “curse of dimensionality” [Hastie et al., 2009; Bellman 1957; Bellman 1961], which results in data becoming sparse and distances between observations becoming uninformative. Consequently, the traditional clustering methods can fail to accurately reveal biological patterns [Orlova et al., 2018] (and see Supplementary Tables 1 and 2 in [Meehan et al., 2019]).

Projection pursuit, on the other hand, aims to find lower-dimensional projections of the data where certain interesting patterns, structures, and features manifest themselves. This is motivated by the fact that in many situations the relevant information (such as cluster relationships) is contained in a lower dimensional subspace [Hastie et al., 2009], with the remaining dimensions being uninformative. Projection pursuit involves finding projections that maximize some criteria or interesting property. Once an interesting set of projections has been found, existing structures (clusters) can be extracted and analyzed separately. Projection pursuit can reveal hidden structures and relationships in the data that might be difficult to detect in the original high-dimensional space due to the curse of dimensionality. However, it requires defining a criterion or property to optimize for, and the choice of criterion will determine the types of patterns that the optimization will search for. Additionally, identifying the right projection can be computationally intensive since one needs to explore d^N projections, where d is the data set dimensionality, and N is the dimensionality of the low dimensional projection. While the latter challenge is computational rather than a fundamental scientific limitation, it can be a serious practical hurdle.

The concept of exhaustively exploring low-dimensional projections of high-dimensional data has existed for a few decades. Historically, efforts were made to systematically explore low-dimensional projections, referred to as a “grand tour” [Cook et al., 1995], or to optimize specific criteria to identify profitable projections. However, challenges in determining the optimal criterion and the computational complexities associated with processing numerous low-dimensional projections have hindered the widespread adoption of projection pursuit methods for data clustering tasks.

Here we combined the principles of projection pursuit [Friedman et al., 1974; Friedman et al., 1982; Huber, 1985] and clustering into an Automated Projection Pursuit (APP) clustering approach to automatically uncover interesting structures in high-dimensional data. In traditional projection pursuit, the analyst manually adjusts the projections to find interesting patterns. In the APP approach, as implemented here, we automatically find the low dimensional projections with the smallest data density between the resulting clusters and further recursively and exhaustively analyze each resulting cluster until no further splits in the data are detected. This allows for the reproducible discovery of clusters that might be obscured in the high-dimensional space, and, as we show here, is less sensitive to the curse of dimensionality. Finally, to address the challenging computational aspects of projection pursuit clustering we developed an algorithm to efficiently find the most profitable projection at each recursive step.

We demonstrate the versatility and accuracy of APP applied to various data modalities, including flow and mass cytometry data, scRNAseq, multiplex imaging data, and TCR repertoire data. Our results show that APP recapitulates experimentally validated cell type definitions in a variety of data modalities, and offers additional biological insights. These insights include evaluating hypotheses regarding the existence of a binding motif between in CDR3b of TCRs that recognize the same peptide, as well as uncovering a pattern involving charged amino acid residues that contribute to stabilizing the interface between CDR3a and CDR3b chains of TCR.

To gain a deeper understanding of scenarios where projection pursuit methods, such as APP, outperform widely used high-dimensional clustering methods, we initially applied them to biological data with a known ground truth. In this dataset, each cell population was quantified and functionally validated. To facilitate a quantitative evaluation of APP’s performance compared to other methods, we developed a label transfer pipeline based on supervised UMAP (https://umap-learn.readthedocs.io/en/latest/supervised.html). This approach provides a convenient framework for comparing and visualizing the agreement between clustering algorithm methods and the ground truth and accounts for the underlying data topology. This aspect is particularly vital, especially in the context of analyzing data from disease settings. As demonstrated in an illustrative example, our analytical approach enabled the discovery of a novel population of myeloid cells notably enriched in hospitalized COVID-19 patients.

## Materials and methods

### Data Overview

#### Flow cytometry data

Whole blood from consenting COVID-19 patients and healthy donors (studies approved by the Emory Institutional Review Board (IRB) under protocol numbers IRB00058507, IRB00057983, and IRB00058271) were collected by standard venipuncture, then samples were processed as previously described [Eddins et al., 2022]. Peripheral blood mononuclear cells (PBMCs) were isolated from whole blood following serum collection using the EasySep™ Direct Human PBMC Isolation Kit (StemCell Technologies) following the manufacturer’s instructions. We then performed either a custom monocyte-enrichment procedure (via negative selection) utilizing Mojosort™ anti-PE Nanobeads (BioLegend) and PBMCs stained with PE-conjugated CD3ε (clone: UCHT1), CD19 (clone: SJ25C1), CD56 (clone: NCAM), and CD57 (clone: HNK-1; all from BioLegend) for COVID-19 samples or the EasySep™ Human Monocyte Enrichment Kit without CD16 Depletion kit (StemCell Technologies) for health donor samples.

An aliquot of monocyte-enriched PBMCs (<10^7^ total) was resuspended in fluorescence-activated cell sorter (FACS) buffer in 5 mL FACS tubes and pre-incubated with Human TruStain FcX™ (BioLegend). The 28-color extracellular staining master mix (Supplementary Table 1) was prepared 2X in BD Horizon™ Brilliant Stain Buffer (BD Biosciences) and added 1:1 to cells. After staining, cells were fixed with 4% paraformaldehyde, then washed with FACS buffer, and resuspended in 200-1000 µL FACS buffer for acquisition using BD FACSDiva™ Software on the Emory Pediatric/Winship Flow Cytometry Core BD FACSymphony™ A5.

#### Mass cytometry (CyTOF) data

A comprehensive 38-parameter mass cytometry panel was applied to healthy human blood samples from ten consenting volunteers to compare the frequencies of 28 immune cell subsets [Toghi Eshghi et al., 2019]

#### scRNAseq data

PBMC dataset: Publicly available scRNA-seq counts data from 2,700 single peripheral blood mononuclear cells (PBMC) was accessed from 10X Genomics (https://cf.10xgenomics.com/samples/cell/pbmc3k/pbmc3k_filtered_gene_bc_matrices.tar.gz).

Wildtype and 5XFAD mouse model dataset: Single-nucleus RNA-seq counts data for 3 wildtype and 3 5XFAD 7-month-old mouse brains was downloaded from the Gene Expression Omnibus (GEO) database (accession number GSE140510) [Zhou et al., 2020]. Droplet-based 5’ end massively parallel single-cell RNA sequencing had been performed on the samples, and data processing was done using the Cell Ranger Single-Cell Software Suite from 10x Genomics by the originators of the data.

#### Multiplex imaging data

CD11c, SIRPɑ, CD163, CD206, CD68, CD45, HLA-DRA and Pan-Keratin antibodies were conjugated to oligonucleotides (oligos) and then validated in the SignalStar Multiplex IHC assay to assess the myeloid compartment of the tumor microenvironment. Paraffin-embedded human squamous cell carcinoma tissue was tested using 8-plex panel Pan-Keratin (C11) & CO-0003-488 SignalStarTM Oligo-Antibody Pair #63566, CD68 (D4B9C) & CO-0007-594 SignalStarTM Oligo-Antibody Pair #77318, CD206/MRC1 (E2L9N) & CO-0035-488 SignalStarTM Oligo-Antibody Pair #99626, CD163 (D6U1J) & CO-0022-750 SignalStarTM Oligo-Antibody Pair #71043, SIRPɑ/SHPS1 (D6I3M) & CO-0034-647 SignalStarTM Oligo-Antibody Pair #80150, CD45 (Intracellular Domain) (D9M8I) & CO-0013-647 SignalStarTM Oligo-Antibody Pair #32740, CD11c (D3V1E) & CO-0017-594 SignalStarTM Oligo-Antibody Pair #85384, HLA-DRA (E9R2Q) & CO-0023-750 SignalStarTM Oligo-Antibody Pair #58446 using SignalStarTM mIHC technology.

All 8 primary antibodies are applied at once in one primary incubation step. A network of complementary oligonucleotides with fluorescent channels 488, 594, 647, 750 nm amplify the signal of up to 4 oligo-conjugated antibodies in the first round of imaging, followed by removal and amplification of 4 additional antibodies in the second round of imaging. Images were acquired on the PhenoImager HT (Akoya Biosciences). The antibodies were quantitatively validated in the SignalStar assay to ensure maximum fluorescent signal with minimal background, and compared against the chromogenic gold standard.

#### TCR repertoire data

The TCR repertoire data utilized in this study was obtained from the McPAS-TCR database [Tickotsky et al., 2017]. McPAS-TCR is a manually curated resource containing human and mouse TCR sequences associated with various pathologies and their cognate antigens. We downloaded the September 10, 2022 version of McPAS-TCR (latest available), providing over 13,000 TCR CDR3-beta chain and epitope pairs.

### Data analysis

#### Flow cytometry data

Manual, use-guided analyses were performed using AutoGate [Meehan et al., 2019] (available at: cytogenie.org) and FlowJo™ v10.8 (BD Biosciences).

#### Mass cytometry (CyTOF) data

Comprehensive conventional manual gating strategy for 38-parameter human immunophenotyping is described at [Toghi Eshghi et al., 2019].

#### scRNAseq data

Both datasets were processed using Seurat v3. Counts were log-normalized and scaled, and UMAP reduction and PCA (principal component analysis) were performed. Seurat’s default graph-based clustering algorithm was used to identify cell-type clusters (https://satijalab.org/seurat/articles/pbmc3k_tutorial.html). Cell types were annotated by comparing known biomarkers with the markers calculated for each cluster.

Principal components (10 for the PBMC dataset and 20 for the brain dataset [Zhou et al., 2020]) were extracted from the Seurat objects in order to run the APP clustering procedure, which produced new cluster identifications for each cell. New cell annotations were identified by comparing known biomarkers with cluster markers calculated using the new identifications. Differently-matched and non-matched cells were identified and tabulated. Dimensional reduction plots and heat maps were produced using Seurat, and other visualizations were produced using ggplot.

#### Multiplex imaging data

To interpret SignalStar data collected in two imaging rounds (Round 1 and Round 2) with the PhenoImager HT (Vectra Polaris) several data pre-processing and processing steps were performed. More specifically, whole slide imaging data collected with PhenoImager HT underwent image stamping and whole section selection in the Phenochart (Akoya Biosciences), with the further spectral unmixing and autofluorescence removal done in the Inform (Akoya Biosciences) to distinguish true signals from background noise and ensure accurate quantification of each fluorophore signal.

TIFF components were then exported from the Inform (Akoya Biosciences) into the QuPath [Bankhead et al., 2017] software where the TIFF components stitching, image alignment co-registration, and image fusion were sequentially performed. These steps allow simultaneous visualization of multiple markers signals from which were recorded across different cycles or time points.

Nuclear and membrane segmentation were then done using Cellpose QuPath extension [Stringer et al., 2021] on the pre-processed images. Specifically, the ‘nuclei’ base model of the Cellpose algorithm was employed, with the DAPI nuclear signal from Round 1 serving as its input. Expected diameter of detected nuclei was set to zero to allow for automatic computation by Cellpose. To approximate cell boundaries, a nucleus expansion algorithm implemented in Cellpose was employed, with the cellExpansion parameter set to 5 micrometers. Cell expansion was constrained to 1.5 times the size of the nucleus, controlled by the cellConstrainScale parameter. Additionally, tile size was set to 2048 pixels and the setOverlap parameter that accounts for overlaps between the tiles was set to 100 pixels.

Following segmentation twenty features per marker were extracted for each single cell and used for the subsequent cellular analysis including cell phenotyping. Specifically, measurements of marker mean, median, maximum, minimum, and standard deviation were calculated for each of the identified cell compartments, namely the nucleus, cytoplasm, membrane, and the entire cell.

#### TCR repertoire data

The TCR CDR3b, CDR3a sequences and associated peptide epitope sequence data underwent conversion into embeddings using recent techniques in Large Language Model (LLM) technology. Evolutionary Scale Modeling (ESM) [Lin et al., 2022] has recently harnessed LLMs to create a collection of protein language models. Specifically, we utilized the esm2_t33_650M_UR50D model from ESM to initially generate embeddings for TCR CDR3b, CDR3a and peptide epitope sequences. These embeddings, characterized by a high dimensionality (1280 dimensions), were independently created for TCR CDR3b, CDR3a, and peptide epitope sequences. Subsequently, we concatenated the embeddings (1280D for TCR CDR3b and 1280D for peptide; 1280D TCR CDR3a, 1280D CDR3b, and 1280D peptide) to capture the combined information pertaining to TCR-antigen interactions.

The concatenation of these embeddings results in a feature vector (2560D for CDR3b and peptide; 3840D for CDR3a, CDR3b and peptide) that encapsulates the unique characteristics of both the TCR and antigenic sequences. This combined representation aims to capture the intricacies of TCR-antigen interactions. PCA was then applied to these combined embeddings, and the first 30 principal components were used in APP clustering. This approach enabled thorough exploration and analysis of the dataset, unveiling intricate patterns and relationships within the combined TCR-antigen sequence space.

The sequence similarity within each class (CDR3a, CDR3b and peptide epitope) was calculated by aligning each pair of sequences and computing the Blosum62 score for the alignment (utilizing the Bio.pairwise2 module in the Biopython package (https://biopython.org/docs/1.75/api/Bio.pairwise2.html)). The Blosum62 score offers a quantitative measure of the similarity or dissimilarity between amino acids at specific positions in protein sequences, relying on observed frequencies of substitutions in related proteins. It is commonly utilized in sequence alignment algorithms to assess the evolutionary relationships between proteins and identify regions of conservation or divergence. Within each cluster, an average sequence similarity was calculated by averaging the scores for each unique pair of sequences found in that cluster. Between each pair of clusters, an average sequence similarity score was calculated by averaging the scores for each unique pair of sequences between the two clusters. Unique pairs were used to avoid biasing within-cluster average scores for clusters containing many repeated sequences.

Cluster sequence logos for peptides were generated by selecting all sequences of uniform length. This length was defined as the rounded average sequence length among all sequences in a cluster within a given class of sequences. This approach ensured the ∼70-80 percent (varies among the clusters) coverage for epitope sequences. The distribution of amino acid residues at each position was then calculated for these selected sequences. The Python package LogoMaker was employed to create probability matrices for the sequence logos, which were subsequently utilized for the analysis of amino acid R group properties.

All CDR3a and CDR3b sequences used in this study underwent alignment using the multiple alignment program for amino acid or nucleotide sequences (MAFFT), accessible at https://mafft.cbrc.jp/alignment/server/large.html?aug31, with default settings. The aligned sequences were further employed for the analysis of amino acid R group properties.

#### TCR-pMHC crystal structure analysis

Crystal structures under PDB accession codes 3GSN, 3PQY, 1OGA, 3O4L and 5EUO were used to analyze in detail interfaces between antigen and TCRa, antigen and TCRb, TCRa and TCRb. For each structure the analysis was performed using PISA (http://www.ebi.ac.uk/pdbe/prot_int/pistart.html) service [Krissinel et el., 2007]. A manual verification was performed for each structure using PyMOL Molecular Graphics System, Version 1.2r3pre (Schrödinger, LLC). Figures were generated with PyMOL.

#### Automated projection pursuit clustering based on the best separation score

The objective of determining the decision boundary with the lowest data density between resulting clusters at 2D projection shares a conceptual similarity with Hamilton’s principle of least action [The Feynman Lectures on Physics Vol. II Ch. 19: The Principle of Least Action], a fundamental variational principle in particle and continuum systems. According to Hamilton’s formulation, the genuine dynamical trajectory of a system, navigating between an initial and final configuration within a specified time, is identified by contemplating all conceivable trajectories that the system might follow. For each of these trajectories, the action—a functional of the trajectory—is computed, and the trajectory that renders the action locally stationary (traditionally termed ‘least’) is selected. The genuine trajectories are those that minimize the action. The overarching concept of examining all conceivable trajectories (or decision boundaries in our case) to identify the most optimal solution served as the foundational principle in devising the logic behind the APP algorithm.

The overall data clustering workflow (Figure 1) is constructed to unambiguously assign a cluster identification number (id) to each data point in the data set by recursively performing the following three steps: 1) presenting the multidimensional data in all its two-dimensional (2D) orthogonal projections; 2) for each 2D projection finding the decision boundary (i.e. the boundary separating cluster assignments) according to the local minimum density of the data points; 3) choosing the 2D projection that has decision boundary with the highest Calinski-Harabasz [Caliński et al., 1974] score, splitting the data along this decision boundary. The Calinski-Harabasz index, that is often used to evaluate the goodness of split, is calculated as a ratio of the sum of inter-cluster dispersion and the sum of intra-cluster dispersion for all clusters (where the dispersion is the sum of squared distances). These steps 1 and 2 are repeated recursively and exhaustively, until there are no further splits as defined by the user-inputed minimum cluster size parameter.

**Figure 1.**
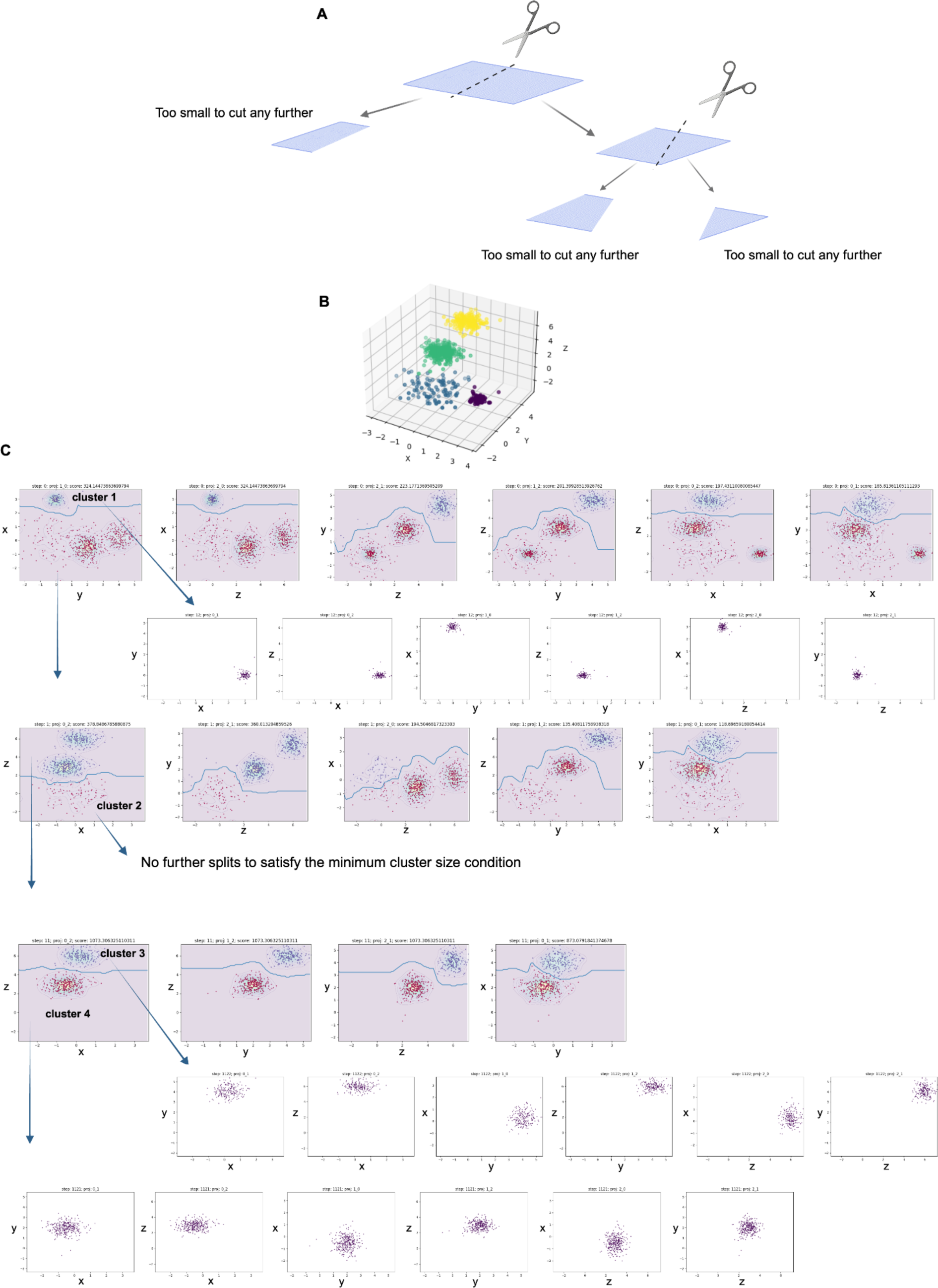
Automated projection pursuit clustering workflow. **A.** Metaphorical representation of projection pursuit clustering, where the paper sheet represents multidimensional data (implying that the data is complex and exists in a higher-dimensional space) projected into a pair of dimensions with the best separation (all other projections are not shown). Scissors symbolize the process of splitting the data into segments along a decision boundary (dotted line). This step reflects the idea of sequentially finding the most informative projections to separate the data effectively. **B.** To illustrate the concept behind the APP algorithm in a simplified manner we used a three-dimensional synthetic dataset. **C.** The APP algorithm systematically explores orthogonal two-dimensional projections, selects the one with the smallest density distribution along the decision boundary (represented by a blue line on the two-dimensional projections), and recursively splits the data until the defined stop criteria are met. This approach helps uncover meaningful patterns and structures in the data.

The generalized structure for such an algorithm may look as follows.

In each recursive step:

If the number of cells at the input to the recursion step is less than 2* min_cluster_size (user defined parameter), then this piece of data is considered as the final cluster, cluster id is assigned, and no further splits are performed. The algorithm exits the recursion step.

Alternatively, for each 2D dimensional (x,y) projection mapped onto a unit square (side length of 1) we build the Gaussian-smoothed histogram H(x,y) of the data points. For the optimal number of bins of 2D histogram, we use Mann’s formula [Mann et al., 1942] taking into account the number of the data points *n* of the current 2D projection. The challenge of determining the optimal number of histogram bins, contingent on the number of data points, remains an ongoing issue without unanimous consensus in the literature. The optimal choice must strike a balance between having too few bins (resulting in poor resolution) and too many bins (leading to increased noise). Various approaches and formulas exist, and in our perspective, we adopted what we believe to be the most reasonable one. The Gaussian smoothing width σ is taken as a free parameter of the algorithm. Then the number of bins *N* for each of the two coordinates is the square root of the total number of two-dimensional bins multiplied by the width of the Gaussian smoothing:

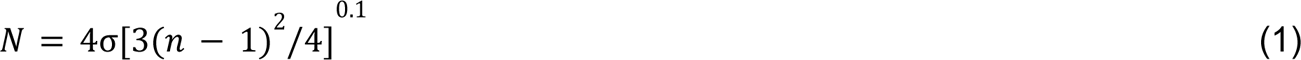

Thus, the use of Gaussian smoothing not only reduces the statistical noise of the data, but also allows to increase the number of histogram bins. Calculating the optimal number of histogram bins depending on the number of data points considered at each recursion step made it possible to significantly speed up the calculations, since it reduced the number of algorithm operations due to the reduction in the size of clustered projections during program operation.

To initiate the search for a decision boundary function *y*(*x*) (Supplementary Figure 1) we build *H*_1_function that is a sum of the initial distribution density *H* and a parabolic function *H_g_* - the “*y*_0_-gravity potential”. Function *H_g_* (*y*) is added for partial straightening the decision boundary function *y*(*x*) along the *x*-axis. Addition of the *H* (*y*) function to the data histogram results in a constraint for the following decision boundary condition: *y*(*x*_0_, *q*) = *y*_0_ (*q*) and in a constraint that *y*(*x*, *q*) will be as close as possible to the *y*_0_ (*q*) for the every **x* ∈* [**x*_min_*, **x*_max_*].

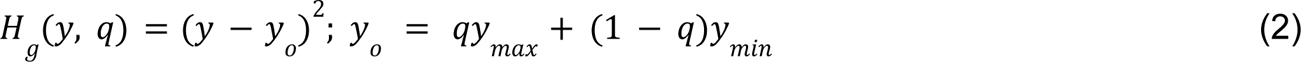

Where *q* ∈ [0, 1] - a numeric parameter.

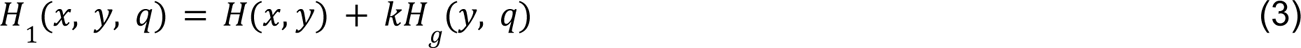

Where *k* is a positive constant, the optimal value of which is calculated by the following expression:

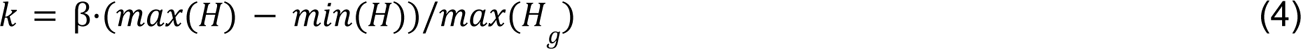

The coefficient *k* plays a crucial role in achieving a balance between the parabola and the data, particularly in determining the trajectory of the decision boundary. And the greater the coefficient *k* the greater the straightening effect. Нere the multiplier β is a free parameter that allows to vary the degree of influence of the “*y*_0_ -gravity potential” on the clustering process. Our empirical assessment shows that a value of β=0.1 (i.e. 10% “gravity”) gives fairly good clustering results in many cases.

To “draw” a decision curve on a 2D plane one needs to know its’ initial [*y*(*x_min_*)] as *y* and final [*y*(*x_min_*)] as *y* boundary conditions. The explicit but more computationally intense solution would be to search for a decision boundary for each possible *y*_0_ and *y*_1_ in a given 2D projection. To optimize this process, instead of that approach we made the parabolic function *H_g_* be depend on a parameter *q* that is used to introduce the next incremental step along the axes (i.e., δ*q* = 0.1).

To find the decision boundary with the smallest data density between the resulting clusters, for every parameter value *q* ∈ [0, 1] (Supplementary Figure 1) we search for extremals *f_q_* (*x*) (an analogue of trajectory from analytical mechanics) and the curvilinear integral S(*q*) (an analogue of action from analytical mechanics) of the probability density along the decision boundary:

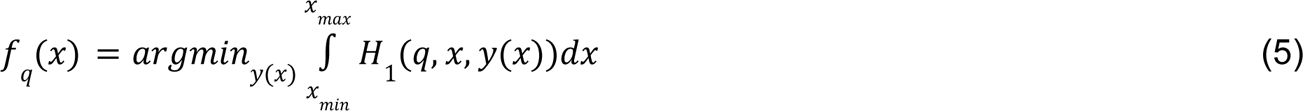

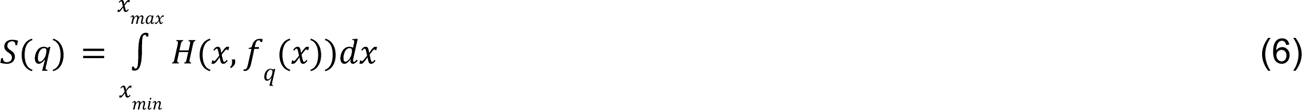

Further, we need to find the values *q*_0_, that would satisfy the following condition: *S*(*q*_0_− δ*q*) > *S*(*q*_0_) < *S*(*q*_0_ + δ*q*). For such values, *fq*_0_(**x**) will be a true extremal or decision boundary. Once a decision boundary is found data is then splitted along that decision boundary. If both parts of the data obtained after splitting contain more than min_cluster_size cells, then this decision boundary is added to the list of decision boundaries for a given projection. If more than one decision boundary is found on a given projection, these decision boundaries are then ranked according to their Calinski-Harabasz score, and the decision boundary with the maximum Calinski-Harabasz score is chosen to represent the given projection.

Then, among all possible 2D projections at a given recursion step, the algorithm chooses the one that contains the decision boundary with the maximum Calinski-Harabasz score. The data is then splitted along this decision boundary and the new recursive step is initiated on each of two data pieces independently, if both parts of the data obtained after splitting contain more than min_cluster_size cells. Alternatively, if this latter condition doesn’t hold for any of the projections in the recursive iteration, then the iteration’s inputs form the final cluster. In this case, the algorithm assigns the final cluster id to this piece of data and recursion stops.

#### Automated label transfer across samples

At a high level, the label transfer pipeline enables the use of a labeled (or partially labeled) set of points to learn a metric on the data. This learned metric is then employed as a measure of distance between new, unlabeled points. The immediate practical applications of such a pipeline, demonstrated here, include the automation of an expert-defined manual gating strategy, and the assessment of clustering algorithm performance against the ground truth cluster labels. We have developed a concise four-step pipeline that serves both these needs, facilitating the visualization and quantification of the misclassification rate between the clustering algorithm and the ground truth labels (see Supplementary Figure 2). Additionally, it streamlines the automation of an expert-defined manual gating strategy.

The pipeline starts with creating a supervised UMAP embedding (https://umap-learn.readthedocs.io/en/latest/supervised.html) using labeled or partially labeled training sample(s) that has both marker expression data and ground truth cluster labels. UMAP is applied to this training data with the goal to learn a distance metric that best separates the classes while preserving their relationships in the marker space. Using equal weights for marker expression and ground truth cluster labels ensures that both data-driven and prior knowledge are considered in the embedding.

This step itself provides an opportunity to assess the quality of the ground truth cluster labels by observing an agreement (or disagreement) between the data topology and clustering decisions. By observing the agreement or disagreement between the data’s topological structure in the UMAP space and the provided ground truth cluster labels, one can gain insights into whether the ground truth labels accurately reflect the underlying structure of the data (Supplementary Figure 3). High agreement between the UMAP topology and the ground truth labels suggests that the ground truth labels are representative of the data’s natural clustering patterns. Conversely, disagreements may indicate issues with the ground truth labels.

As we show here, this step is valuable not only for clustering evaluation but also for quality control and refinement of the ground truth labels themselves (Supplementary Figure 3). If discrepancies between the UMAP structure and ground truth labels are identified, it might prompt a reevaluation or improvement of the ground truth annotations. Overall, this dual-purpose step enhances the robustness of the pipeline by allowing one to simultaneously assess the clustering algorithm’s performance and the quality of the ground truth labels used for evaluation.

In the following step (Supplementary Figure 2), the set of labeled points is employed to learn a metric on the data. This learned metric subsequently serves as a distance measure between new unlabeled points, facilitating the projection of an unlabeled test set into the UMAP embedding space constructed using the training set. This ensures that the test set occupies the same reduced-dimensional space as the training set. The test set is then subjected to clustering using the Support Vector Clustering (SVC) [Ben-Hur et al., 2001] algorithm directly applied in the supervised UMAP embedding space. Given that clustering in this context is confined to two dimensions (UMAP_x and UMAP_y), we opted for an algorithm that refrains from assigning any of the events to noise and, at the same time, offers computational superiority over APP. Subsequently, the QFMatch algorithm [Orlova et al., 2018] is employed to align the cluster labels between the test set (with cluster ids defined by SVC or assigned by the clustering algorithm under assessment—see Supplementary Figure 4) and the training set (with ground truth cluster ids). The alignment of labels is crucial as it accommodates the following scenarios: 1) transferring cluster labels from the test set to the training set; 2) directly assessing the agreement of clustering decisions made by multiple clustering algorithms (see Supplementary Figure 4).

In the final step, we compute the number of misclassified events per cluster ID. This quantifies how effectively the clustering algorithm has assigned data points to clusters in comparison to the ground truth.

Beyond clustering algorithm evaluation, this pipeline holds broader applications in supervised learning tasks. As demonstrated here, it can be employed to transfer labels from one sample to another or from a partially labeled dataset to the remaining data in a given set. This feature proves particularly valuable in scenarios where labeled data is limited.

## Results

### Performance on data with functionally validated ground truth labels

To objectively assess the APP performance against the widely used clustering algorithms in application to realistic, biologically relevant data, we used ground truth data where each cell population was quantified and functionally validated. To generate such biologically relevant data with the known ground truth we combined spleen cells from the GFP+Wild-type and RAG-KO mice at five different proportions (Figure 2A). RAG-KO mice are deficient in generating lymphoid lineages, while GFP+ mice represent a healthy immune system with constitutive GFP expression in all immune cell lineages. This allows for easy and accurate detection of immune cell populations by flow cytometry, simplifying the identification and quantitation of cell lineages. The latter ensures that the clusters obtained by clustering algorithms can be compared against a known biological truth [Zimmerman, 2011].

**Figure 2.**
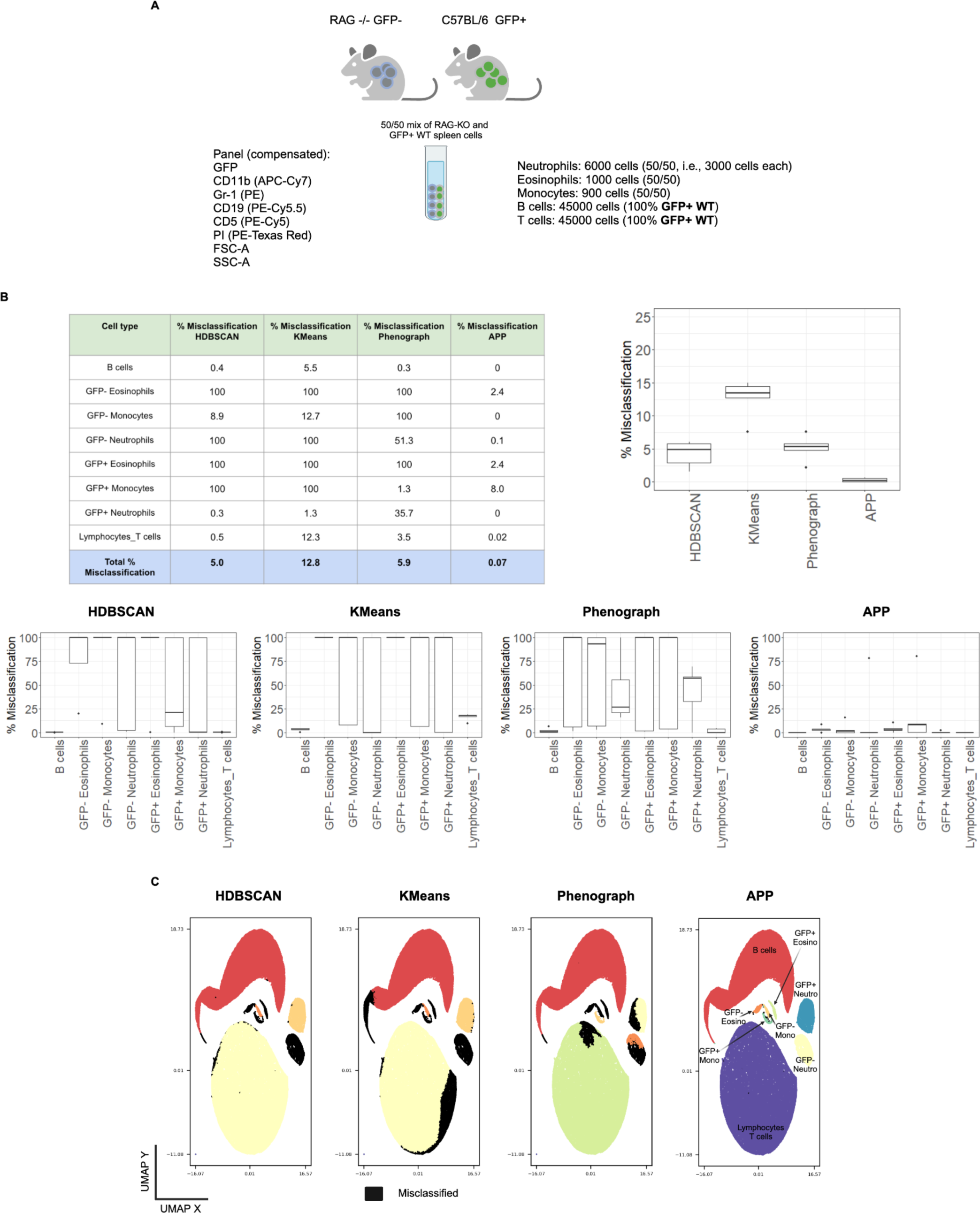
APP outperforms widely used clustering methods in application to the representative biological dataset that has functionally validated ground truth labels. **A.** A representative ground truth sample, selected from several similar types of samples, was generated by mixing cells in equal proportions (50% each) from the GFP+ Wild-type spleen and RAG-KO spleen. It’s important to note that RAG-KO mice lack B and T cells. **B.** The APP algorithm’s performance in cell population classification is being assessed, and its effectiveness is being compared to that of state-of-the-art clustering algorithms. The evaluation encompasses the blending of wild-type spleen cells and RAG-KO spleen cells at five different proportions (50/50, 25/75, 12.5/87.5, 75/25, 87.5/12.5). Evaluation metrics encompass total population misclassification and per cell type misclassification. The results are averaged over five data points corresponding to the distinct cell mixing proportions. The table displays results specifically for the 50/50 cell mixture. **C.** The misclassification for the 50/50 mix is visually represented in black color using the automated label transfer pipeline.

To assess the performance of the APP algorithm in comparison to state-of-the-art clustering algorithms, we selected two generally widely used high-dimensional clustering algorithms, irrespective of data origin: HDBSCAN (https://pypi.org/project/hdbscan/) and KMeans (https://scikit-learn.org/stable/modules/generated/sklearn.cluster.KMeans.html). Additionally, we included Phenograph (https://pypi.org/project/PhenoGraph/), a method widely used in the flow cytometry field and applied to other datasets reported in this study. Our selection of algorithms reflects a diverse range, encompassing density-based (HDBSCAN), centroid-based (KMeans), and graph-based (Phenograph) clustering methods. This ensures a comprehensive comparison, considering different clustering paradigms and their suitability for various data structures.

As demonstrated here, both HDBSCAN, KMeans (K=8), and Phenograph encounter challenges in robustly detecting rare cell populations when coexisting with more abundant cell populations in the same sample (Figure 2B,C). When clustering algorithms operate across multiple dimensions simultaneously, they may face difficulties in effectively detecting and distinguishing sparse populations from more prevalent ones. The increased sparsity within the vast high-dimensional space poses a challenge in identifying clusters that exist in lower-dimensional subspaces. Our findings illustrate that even with seven dimensions (excluding live/dead PI from clustering), clustering algorithms may encounter challenges when dealing with multiple dimensions simultaneously.

Our findings suggest that the APP clustering method excels in scenarios where there are clear distinctions between the cluster under consideration and the other cells in at least one of the dimensions. The APP method’s proficiency in identifying clusters with evident separations in one or more dimensions makes it well-suited for situations where distinct cell populations exist. In such cases, it can leverage the dimension(s) where the separation is apparent to successfully identify and differentiate clusters, even in the presence of much larger populations and noise in the data. This aligns with scenarios resembling cell phenotyping using flow/mass cytometry, imaging antibody panels, and scRNAseq data, where there are often identifiable patterns or markers distinguishing cell types. Another application, as demonstrated here, is clustering molecules, such as TCRs and their cognate peptides, based on their sequence similarity and other features.

On the flip side, high-dimensional clustering approaches may potentially outperform the APP method when there is no clear split between clusters in any of the dimensions, and the information about a given cluster is “distributed” across multiple dimensions. High-dimensional clustering algorithms excel in aggregating information from multiple dimensions simultaneously, which can be advantageous in situations where cluster boundaries are less well-defined. An example of this is in identifying cells’ activation states, highlighting a specific application where high-dimensional clustering may be preferred.

To mitigate some of the limitations of APP in scenarios with less clear cluster separations, we explored incorporating dimensionality reduction techniques, such as PCA, as a pre-processing step. As detailed in subsequent Results sections, this approach proved effective in revealing underlying structures in high-dimensional data, making it more accessible for clustering algorithms to identify meaningful clusters.

### Application to flow and mass cytometry data

The analysis of high-dimensional flow and mass cytometry data, characterized by dozens of parameters, presents two primary challenges: the requirement for unbiased, unsupervised discovery of cell populations within a given panel of markers, and the automation of an expert-defined manual gating strategy. The latter is crucial for clinical applications of flow and mass cytometry data, while the former advances research applications. In this context, we tackle both tasks by introducing an APP for unsupervised discovery of cell populations and an automated label transfer pipeline for the expert-defined gating automation.

Using an expert-defined manual gating strategy as the “gold standard” (Supplementary Figure 5), we evaluated the performance of APP in characterizing PBMCs from healthy donors and COVID-19 patients. APP demonstrated an overall performance accuracy exceeding 95 percent, significantly outperforming one of the widely used clustering algorithms in the flow cytometry field, Phenograph (Figure 3A). The primary source of misclassification for both algorithms stems from sparse cell populations (Figure 3B, Supplementary Figures 6, 7). A detailed comparison (Figure 3B, Supplementary Figure 7) reveals that APP and Phenograph may misclassify different portions of the data, and this discrepancy can be attributed to the distinct clustering logic employed by the two methods.

**Figure 3.**
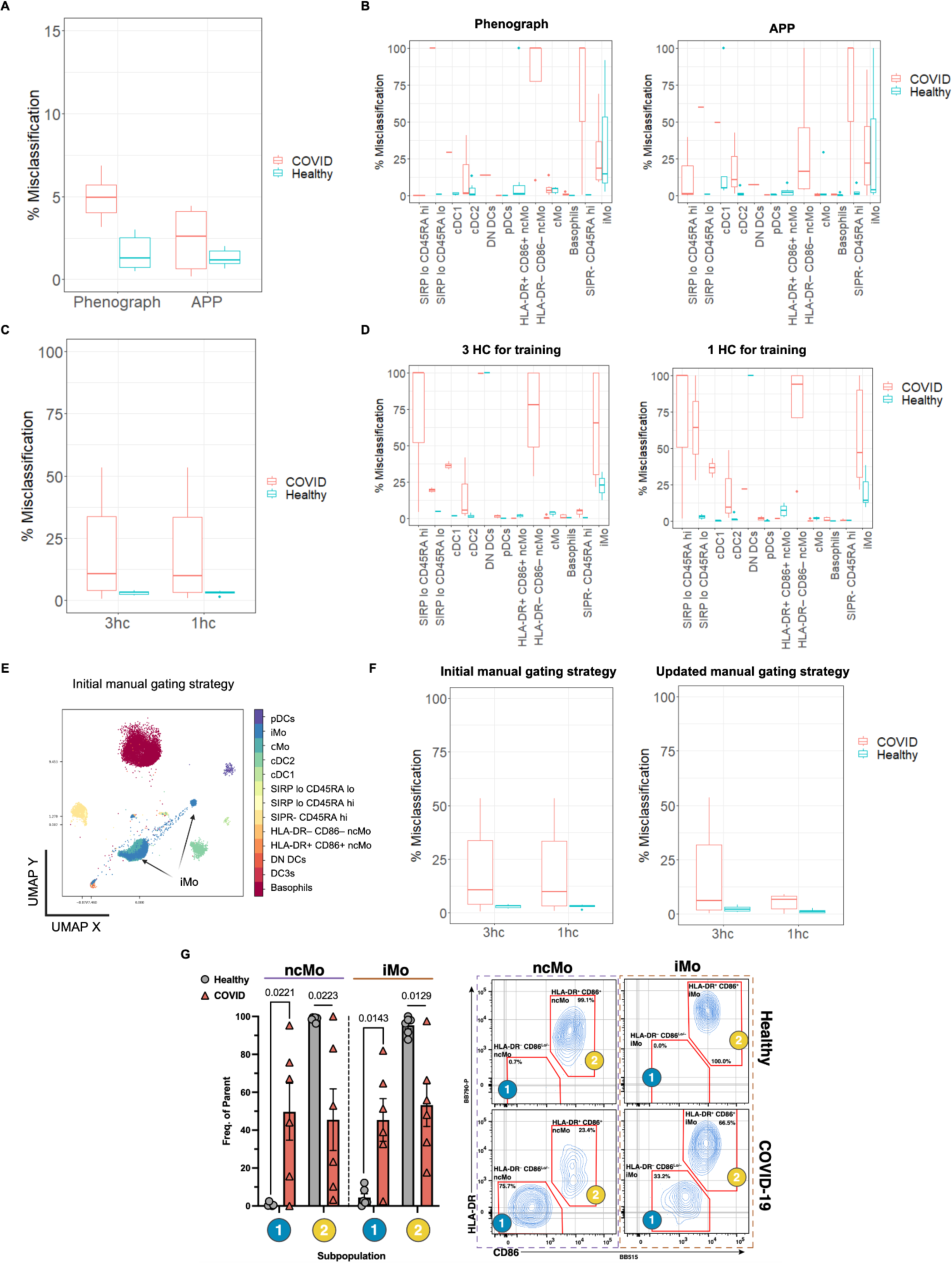
The label transfer pipeline reveals distinct cellular responses in COVID-19 patients and achieves over 99 percent accuracy in automating the manual gating strategy for healthy control samples, using minimal training data. **A.** On average, less than three percent of the data were misclassified when comparing APP clustering decisions to the manual ground truth labels. **B.** The primary source of misclassification between manual and automated clustering decisions is sparse cell types. Cell populations are ordered from most abundant to least abundant (left to right). **C.** The automation of the gating strategy application, implemented via the automated label transfer pipeline, achieves very high accuracy for healthy control (HC or hc) samples. **D.** Misclassification evaluation on a per-cell-population basis allows the identification of populations that are most problematic for the automated gating strategy transfer, providing targeted insights for improvement. As in panel A, cell populations are ordered from most abundant to least abundant. **E.** The label transfer pipeline facilitates the observation of agreement or disagreement between data topology and ground truth cluster labels. It provides insights into the accuracy of ground truth labels and serves as a quality control step. Detected discrepancies (e.g., iMo cell population is more heterogeneous than defined by the original manual gating strategy) may prompt reevaluation or refinement of annotations (**F**), resulting in the discovery of biologically meaningful cell populations (**G**, HLA-DR-CD86lo ncMo and iMo populations that are unique to COVID patients). Minimum cluster size and bin size input parameters of 100 and 50, respectively, were used for both APP and the label transfer pipeline to generate the results presented in panels A-G.

Furthermore, we assessed the label transfer pipeline’s capability to automate the expert-defined gating strategy. The pipeline achieved remarkably high performance for the healthy control cohort, exceeding 99 percent. This level of accuracy was attained even when using just one randomly chosen manually gated healthy control sample as a training set (Figure 3C&D). In the analysis of COVID-19 samples, there were more discrepancies between the manual gating strategy established on healthy control samples and the label transfer pipeline’s outcomes. However, as shown here, this apparent “discrepancy” may indicate that the original expert-defined gating strategy, established on healthy control samples, might require adjustment when applied to disease samples. For instance, as demonstrated here, COVID-19 disease samples exhibit more complex and diverse cell populations than the healthy control samples used to establish the initial manual gating strategy.

Our labels transfer pipeline includes an immediate sanity check to assess the quality of the “ground truth” labels. This check examines whether the ground truth labels align with the underlying data topology. If there is a disagreement, such as a cell population defined as homogeneous in manual gating but being spread across multiple clusters on the supervised UMAP plot, it suggests a potential issue with the original gating strategy (Figure 3E). Indeed, revisiting the original expert-defined gating strategy, as illustrated in Figure 3F&G and Supplementary Figure 5B, led to the discovery of a COVID-19-specific cell population labeled as HLA-DR–, CD86Lo/– intermediate monocytes (iMo).

Thus, APP identified a new population of myeloid cells specifically enriched in hospitalized COVID-19 patients. Notably, this myeloid cell subset lacks cell-surface expression of key proteins relevant for antigen presentation (HLA-DR) and co-stimulation of T-cells (CD86), likely affecting viral antigen presentation and T-cell activation. These findings highlight new mechanisms of SARS-CoV-2-induced immuno-modulation that underlie the COVID-19 immunopathology in hospitalized patients.

The label transfer pipeline was also tested in its application to mass cytometry data, achieving an overall 87 percent accuracy (Supplementary Figure 8) when using the manual gating labels, as described in [Toghi Eshghi et al., 2019]. Discrepancies between the underlying data topology and manually assigned cell populations (Supplementary Figure 8) serve as a source of reduced accuracy for the label transfer pipeline.

### Application to scRNAseq data

In contrast to flow and mass cytometry, which measure dozens of dimensions for each individual cell, the dimensionality of gene expression data is often on the order of thousands of genes per cell. Given the high-dimensional nature of gene expression data, dimensionality reduction techniques, such as PCA (Principal Component Analysis), are often employed as a pre-processing step before clustering to extract meaningful patterns and reduce the computational complexity associated with analyzing a large number of genes. We decided to test APP clustering performance on PCA preprocessed scRNAseq data since dealing directly with the combinatorial combinations of all possible pairwise projections from thousands of genes can be computationally prohibitive.

For this purpose we choose a publicly available scRNAseq dataset generated from human PBMCs and characterized by 10X Genomics. We compared APP clustering performance to the performance of a clustering process using KNN and Louvain algorithms as implemented in the R package Seurat (referred to in this manuscript as “Louvain clustering” and for our purposes, treated as the ground truth since it is a workflow widely used by the field today). The overall misclassification rate between Louvain clustering and APP clustering decisions in the PBMCs dataset is about thirty percent, but less than fifteen percent overall when memory CD4 T cells are excluded (Figure 4 A&B). APP clustering faces challenges in resolving the distinction between Naive and Memory CD4 T cells in the dataset (Figure 4B), likely due to the fact that these cells essentially represent distinct functional states and responses from the same population of cells. And there is no clear split between these two populations in any pairwise dimensions explored by APP (see Supplementary Data) since the differentiation between Naive and Memory CD4 T cells may rely on simultaneous changes in several genes and that gene set can vary based on factors such as the context of the immune response, the microenvironment etc.

**Figure 4.**
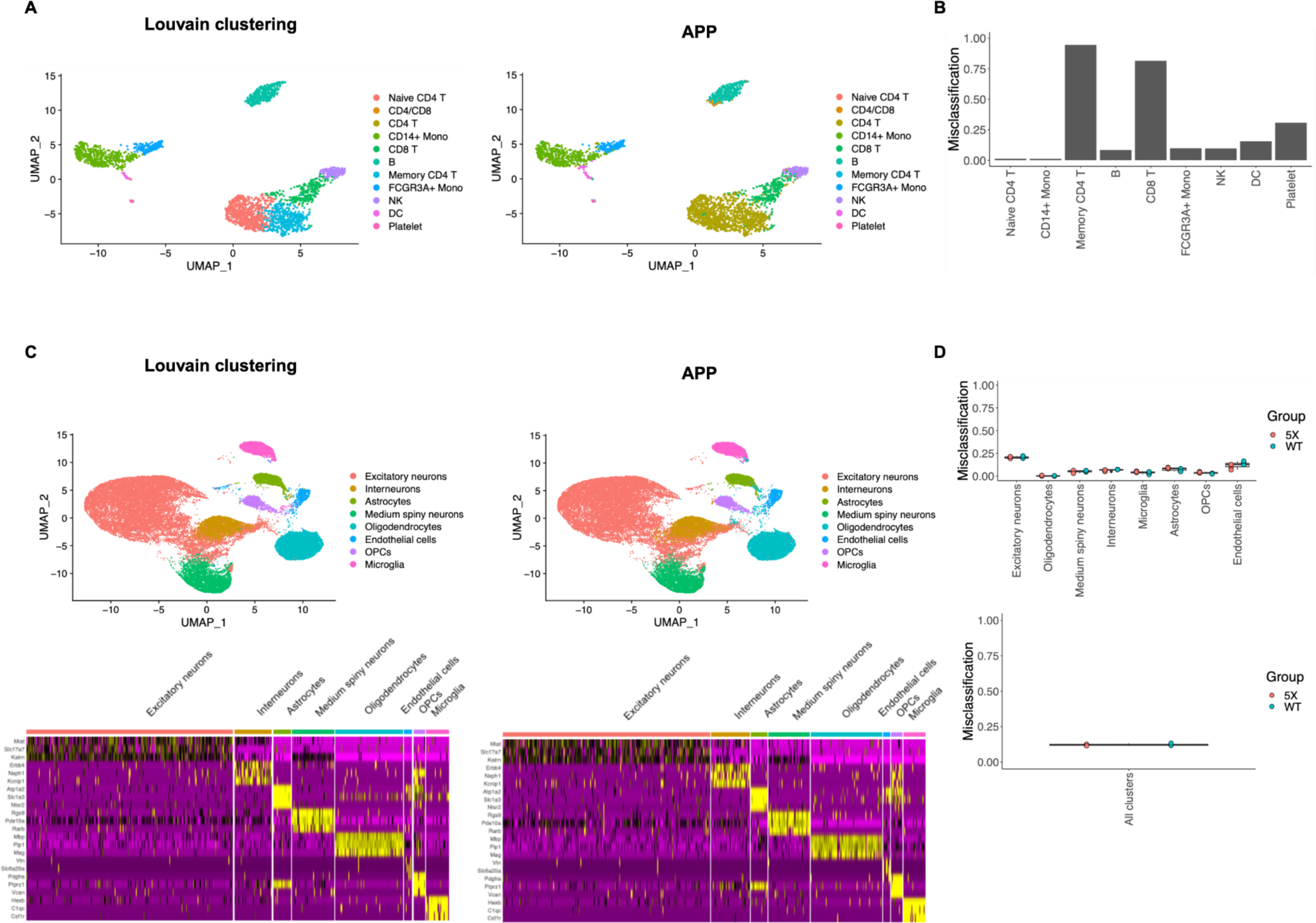
Both Louvain clustering and APP clustering, applied to PCA-reduced scRNAseq data, exhibit good alignment with each other. The overall misclassification rate between Louvain and APP clustering decisions in the PBMCs dataset is about 29 percent. The misclassification rates are determined with reference to the “ground truth” clusters, indicating how many cells in each ground truth cluster produced by Louvain were marked as misclassified. APP clustering encounters challenges in distinguishing between CD4 Naive and Memory cells in the dataset (**A,B**). **C.** There is a high degree of concordance (approximately 87 percent) between Louvain and APP clustering when applied to pooled combinations of wildtype (WT) and Alzheimer Disease model (5X) mice samples. Heatmaps illustrate the concordance of gene expression patterns in each cluster type as identified by the two clustering algorithms. Panel **D** presents per cell type misclassification, providing insights into specific cell types where misclassification occurs.

In our analysis, we further examined the group of cells that Louvain clustering identified as B cells and APP as T cells, labeled as “CD4/CD8” in Figure 4A. We projected this cell population in B cell, T cell and other marker space to gain insights into the APP algorithm decision logic (see Supplementary Figure 9). It becomes apparent that while the “CD4/CD8” cell population exhibits high expression of MS4A1, a B cell-specific marker, it also demonstrates relatively high expression of S100A4, which is a memory CD4 T cell marker, not a B cell-specific marker. This dual expression pattern likely contributed to the source of confusion for APP clustering.

Louvain clustering and APP clustering were next applied to a snRNA-seq mouse brain dataset from the wildtype and Alzheimer disease model (Figure 4C). The overall mismatch rate between the two methods was about thirteen percent, indicating a good level of agreement (Figure 4D). In this dataset, we did not see any “ground truth” clusters that could not be discovered through APP clustering, as we did in the PBMC dataset. All Louvain clusters had equivalent APP clusters with matching marker genes (Figure 4C). The significant areas of discrepancy tended to be in cells which were assigned to their original Louvain clusters with some level of ambiguity.

### Application to multiplex imaging data

Recent advancements in multiplex imaging technologies, exemplified by SignalStar, have greatly enhanced our capacity to profile individual cells within the tissue context. These technologies enable the simultaneous visualization of multiple biomolecules at the single-cell or even the subcellular level. This capability provides valuable insights into cellular heterogeneity, spatial organization, and the composition of tissues.

We applied APP clustering to characterize cellular composition within the tissue context of the human squamous lung carcinoma sample using the 8-plex SignalStar myeloid cell panel. The 8-plex panel (CD11c, SIRPɑ, CD163, CD206, CD68, CD45, HLA-DRA and Pan-Keratin) generated a dataset of 160 features per cell for approximately 230,000 cells. The 160-feature set comprises 8 antibodies x 5 statistics x 4 cell compartments. Here, 5 statistics represent the measurements of marker mean, median, maximum, minimum, and standard deviation, while 4 cell compartments denote the nucleus, cytoplasm, membrane, and the entire cell. We used PCA to reduce the dimensionality of the dataset to 30 dimensions and subsequently performed APP clustering (see Figure 5A&B).

**Figure 5.**
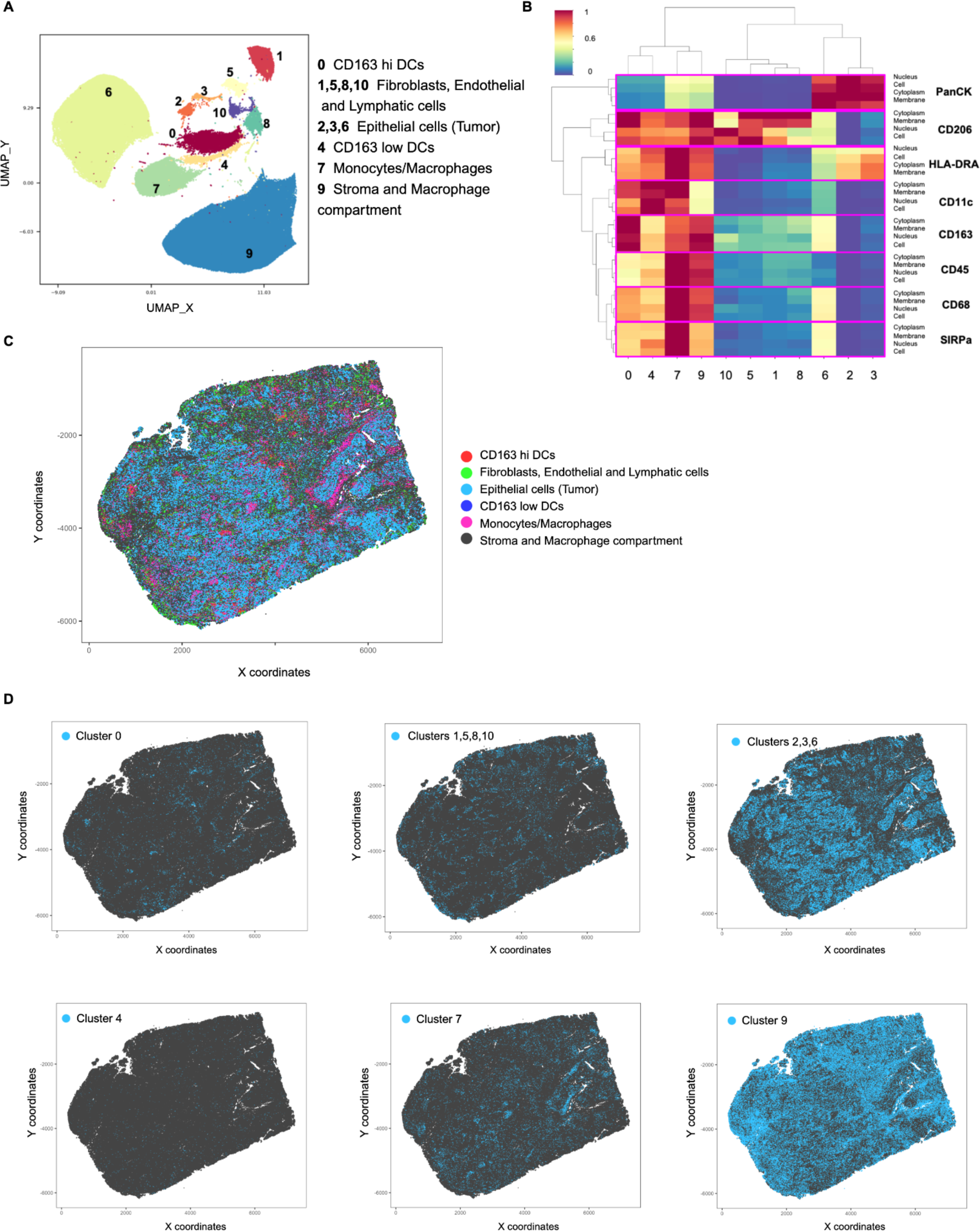
APP clustering successfully leverages multiplexed imaging data to gain insights into biologically meaningful cell populations within the complex tissue context of a human squamous lung carcinoma sample. Eleven cell clusters detected by APP (**A**) were annotated based on the pattern of median marker expression (**B**) and their spatial location (**C,D**).

This clustering approach readily identifies various cell types, including tumor epithelium, stromal, and immune cells. The marker composition of the panel illustrates the phenotypic while reflecting functional heterogeneity within the myeloid compartment: Populations of macrophages and monocytes (Clusters 9 and 7), typically identified by prominent expression of CD68 and CD206, can further be subdivided based on high vs low expression of CD11c and SIRP-alpha [Xu et al., 2017]; similarly, CD11c+ dendritic cells (DCs) can belong to CD163-high or CD163-low (Clusters 0 and 4, respectively), functionally distinct populations [Dutertre et al., 2019; Bourdely et al., 2020; Comi et al., 2020].

The representation of individual clusters (Figure 5A) not only demonstrates phenotypic heterogeneity but also provides valuable clues regarding the relative proportion of phenotypes. Macrophages are known to be prominent cell populations within the microenvironment of diverse human tumor types including NSCLC [Sedighzadeh et al., 2021; Laviron et al., 2022] and based on the size of the combined clusters, macrophages are a highly prevalent cell type in the TME of this squamous NSCLC. PanCK expression identifies epithelial cells, in this particular case cancer cells, unambiguously and the majority of PanCK+ cells seem to be low or negative for all other markers (cluster 6). However, two smaller PanCK+ clusters, 2 and 3, are identified showing elevated levels of HLA-DR; induction of HLA-DR expression in malignant epithelial cells is a known phenomenon, particularly in an inflammatory milieu [Miura et al., 2021; Senosain et al., 2021]. Interestingly, cells of these two clusters seem to preferentially localize to the epithelial-stromal interface with physical proximity between epithelial and immune cells and a potentially high local concentration of inflammatory cytokines.

Cell type label assignments were determined through a combination of marker expression patterns and the spatial location of identified clusters on the tissue slide (Figure 5 C&D). Given that this dataset lacks ground truth labels, known labels for each cell population, the biological relevance of APP clustering decisions was evaluated by domain experts. Pathology and immunology experts independently assessed and confirmed the adequacy of APP clustering in characterizing and distinguishing meaningful cell populations within the slide tissue, particularly in the context of the human squamous lung carcinoma sample.

### Application to TCR-peptide sequence representation data

Recent advancements in Large Language Models (LLMs) have shown significant promise for applications in the analysis of protein sequence data [Madani et al., 2023; Chandra et al., 2023; Ruffolo et al., 2024]. LLMs have been investigated for functional annotation of protein sequences [Quintana et al., 2023], demonstrating capabilities in predicting protein functions, interactions, as well as antibody and TCR specificity [Leem et al., 2022; Wu et al., 2023]. These models excel at capturing contextual relationships within sequences. In the context of TCR sequences, this proficiency involves understanding the specific arrangement of amino acids and their roles in recognizing and binding to antigens. Analyzing the language-like patterns in TCR and antigen sequences using these models may unveil insights into the determinants of specificity.

Although LLMs showcase remarkable capabilities in capturing intricate patterns and relationships within sequences, their black-box nature can pose challenges to interpretability. To address this, clustering techniques can be employed to interpret LLM decisions. In this example using human and mouse TCR sequences and their cognate antigens (Figure 6), we illustrate that the combination of LLMs and APP clustering methods offers a synergistic approach for uncovering patterns, relationships, and semantic structures within large and high-dimensional datasets of TCR-antigen sequences.

**Figure 6.**
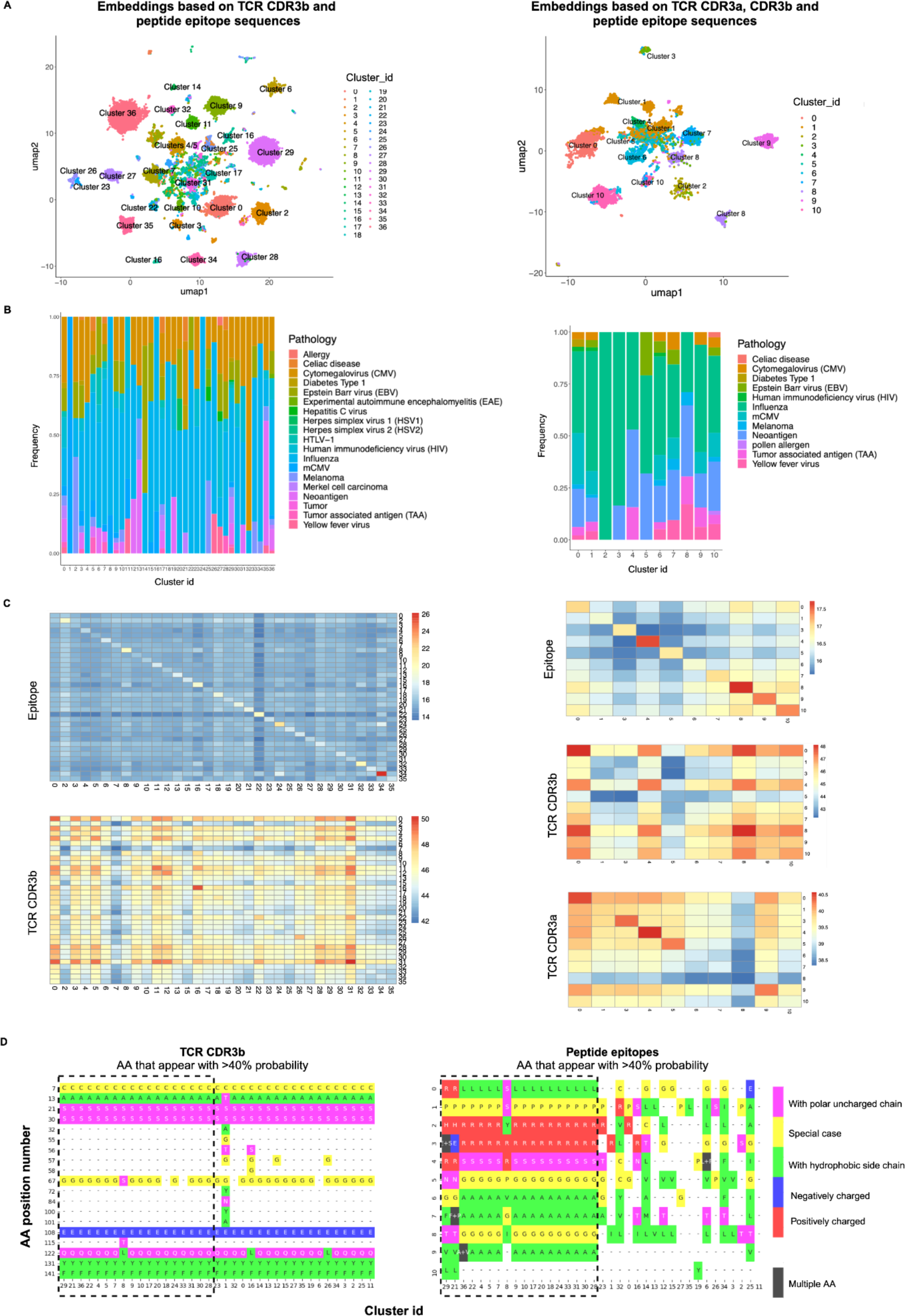
There is no identifiable common binding motif or a universal amino acid R group attribute characterizing the pMHC-TCR interaction. The clusters annotated by APP (**A**) are, in most cases, heterogeneous in terms of associated pathology, but some clusters are disproportionately enriched for particular pathologies, such as Epstein-Barr virus in cluster 14 or cytomegalovirus in cluster 32 in the clustering based on CDR3b and epitope sequences (**B**). Half of the pathologies in this dataset are associated with multiple unique epitope sequences, with an average of 8 unique epitope sequences per pathology. When sequence similarity scores are calculated for each pair of unique epitope sequences in the dataset and those scores are averaged across each unique pair within a cluster and and across each unique pair between all pairwise clusters, average similarity scores are systematically higher within clusters—on the diagonal line in these heatmaps—than between clusters. The same pattern is observed for CDR3b and CDR3a sequences to a lesser degree (**C**). Note that Cluster 2 from the CDR3b, CDR3a, and epitope clustering was removed from the epitope and CDR3a heatmaps because it contained < 20 unique epitope sequences (all other clusters had between 41-125 unique epitope sequences), causing it to have average within-cluster scores high enough to visually overwhelm the heatmaps, so as to allow the patterns between other clusters to be more perceptible, and clusters 1, 14, and 36 were removed from the CDR4b and epitope clustering because their average within-cluster scores were also visually distracting outliers. Per cluster amino acid enrichment analysis (**D**) has unveiled that, while there is no singular characteristic universally defining TCR-pMHC interactions, specific clusters may demonstrate shared electrostatic patterns involving charged amino acids at the interface. A noteworthy example includes a set of clusters (highlighted with a dashed line) enriched with the LPRRSGAAGA peptide, characterized by positively charged amino acids.

To illustrate this point, we created embeddings for over 13,000 unique data points containing TCR CDR3b and cognate epitope sequence information for over 350 unique peptides (left side of Figure 6A). Additionally, we independently generated embeddings for a subset of approximately 4000 unique TCRs with sequence information for both TCR CDR3a and CDR3b (right side of Figure 6A). These embeddings were generated independently for each sequence class (CDR3a, CDR3b, and epitope) and then concatenated for CDR3b-epitope and CDR3a/b-epitope. PCA was subsequently performed, and the first 30 PCs were utilized for APP clustering, employing a minimum cluster size of 100. The composition of identified clusters was characterized based on epitope content and their association with disease information retrieved from the McPAS-TCR database (Figure 6B).

By combining embeddings constructed independently for TCRs and peptides, we generated a fused representation that captures similarities on both the TCR and peptide sides (Supplementary Figure 10). This approach highlights co-similarities between TCRs and peptides across multiple pairs, enabling exploration of the joint sequence space of TCR-peptide interactions. This comprehensive view could offer valuable immunological insights into the complexities of immune recognition, cross-reactivity, specificity, and diversity.

Nevertheless, the inherent black-box nature of LLMs presents a challenge in immediately interpreting the rationale behind the grouping of data points within embeddings. Introducing a clustering approach to these embeddings and subsequently overlaying domain knowledge and interpretable feature information onto the clustering decisions provides an opportunity to partially unravel the decision-making process of LLMs. This method allows for a more nuanced understanding and interpretation of the patterns and associations encoded within the model, bridging the gap between the complexity of LLMs and the need for interpretability in various applications.

One evident hypothesis to investigate was assessing whether sequence similarity played a significant role as one of the driving forces behind the clustering of TCR-peptide data points within the class-separated and the combined embeddings. This investigation aimed to uncover the impact of sequence similarity on the grouping patterns within the combined embeddings, providing valuable insights into the underlying factors influencing the model’s behavior. To evaluate this hypothesis, we calculated sequence similarity independently for peptides, TCR CDR3b, and TCR CDR3a among all pairwise combinations of clusters (refer to Figure 6C and Supplementary Figure 10).

The within-cluster average sequence similarity consistently displayed higher scores compared to inter-cluster average sequence similarity in all three cases: for peptides, CDR3a, and CDR3b. This is evident from the observed higher Blosum62 score diagonal pattern on the heatmaps (Figure 6C, Supplementary Figure 10, Supplementary Figure 11 A,B). As depicted in Supplementary Figures 10 and 11 sequence similarity, particularly in cases of peptide epitopes, plays a significant role in data clustering when class-separate embeddings are constructed. A clear pattern of peptide clustering based on disease categories (pathogens vs cancers, etc.) is observed. However, once TCR and peptide embeddings are concatenated, the data points undergo significant rearrangement in the embedded space. The disease of origin becomes a less significant feature for peptides. Overall, while sequence similarity between peptide epitopes and between TCRs still contributes to clustering, it becomes less prominent, as evidenced by a significant decrease in the median Blosum62 score on the diagonal, particularly for peptide epitopes.

It’s noteworthy that certain peptides grouped together, such as those within cluster 4 on the right side of Figure 6B, originate from antigens associated with diseases that are not immediately related. For instance, cluster 4 predominantly consists of peptides derived from cancer neoantigens and Influenza. What’s particularly interesting is that TCRs recognizing these peptides also exhibit a closer sequence similarity in their CDR3a and CDR3b regions within cluster 4 compared to TCRs outside of this cluster (Figure 6C). This suggests a potential shared recognition pattern among TCRs responding to sequence-similar peptides from disparate diseases within the same cluster.

The clustering based on sequence similarity indicates a commonality in the TCR responses, implying a common recognition pattern that extends across distinct disease contexts. From an immunology perspective, this observation could potentially be explained via TCRs’ promiscuity towards structurally similar epitopes, even if they originate from different antigens. This promiscuity could be facilitated by commonalities in binding motifs among TCRs. Nonetheless, our analysis, illustrated in Supplementary Figure 11, reveals the absence of an immediately discernible common binding motif. This holds true even among CDR3b sequences that specifically recognize the same peptide epitope (LPRRSGAAGA). It is worth noting that the promiscuity works both ways, namely that a TCR can recognise multiple epitope peptides, but also an epitope peptide can be recognized by multiple TCRs. Our clustering results show that the recognition patterns can be very different, since different peptide-TCR pairs corresponding to the same peptide are sorted into different clusters in many cases (Supplementary Figure 11).

By further exploring the joint sequence space, we observed a novel pattern of “similarity” between the data points emerging. This led to significantly different within-cluster arrangements for both peptides and TCR CDR3b. The label transfer pipeline was employed to quantitatively assess the degree of rearrangement that both peptide and TCR CDR3b classes undergo between the single class embeddings and concatenated embeddings (Supplementary Figure 12). Only 23 percent of peptide epitope data (and 12 percent for TCR CDR3b) is arranged in similar patterns between the single class and concatenated embeddings space. This suggests that, during the exploration of the joint sequence space of TCR-peptide interactions, a new rationale for data point similarity that is not necessarily relevant to the origin of the antigen peptide emerged.

In our pursuit of a deeper understanding of the arrangement of data points in the joint sequence space, we explored the possibility of utilizing the characteristic properties of amino acid R groups as a novel similarity criterion. However, we found no immediate patterns of complementarity or cluster-specific motifs defined by the properties of amino acid R groups (refer to Figure 6D, Supplementary Figures 10&11). The diversity among TCRs and the peptides they recognize suggests that the mechanisms of interaction can vary significantly between individual TCR-peptide pairs, with no obvious rules being detected so far. For instance, in Supplementary Figure 13, we highlight a case where a positively charged residue from the peptide side interacts with the TCR chains through the formation of hydrogen bonds, underscoring the intricate and diverse nature of these molecular interactions.

An intriguing pattern did emerge, though, revealing a prevalence of negatively charged amino acid residues on CDR3b and positively charged residues on CDR3a chains across almost all identified clusters (see Supplementary Figure 13). To delve into the potential significance of these charged residues, we meticulously examined five TCR-pMHC crystal structures (PDB accession numbers 3GSN, 3PQY, 1OGA, 3O4L, 5EUO). Upon closer investigation of these structures, it became evident that these charged amino acid residues play a role in stabilizing the interface between CDR3a and CDR3b, orienting partners to each other in three-dimensional space. Most frequently, this stabilization occurs through hydrogen bonds, although there is also an instance of ionic interaction (Supplementary Figure 13E).

## Conclusions

Identifying robust patterns in high-dimensional data is a challenging and crucial task in various scientific fields. High-dimensional data often poses challenges such as the curse of dimensionality, where traditional methods may struggle due to increased sparsity.

Projection pursuit has been proposed as a potential approach to address or mitigate some of the challenges posed by the curse of dimensionality. As a concept this technique has been around for several decades, and its development and application span across different fields such as statistics, machine learning, and data visualization. The idea of seeking interesting projections of high-dimensional data can be traced back to the 1970s and 1980s [Kruskal, 1969; Friedman et al., 1974; Friedman et al., 1982; Huber, 1985].

While the concept of projection pursuit has been around for decades, high-dimensional clustering has gained more popularity and attention, especially in the context of the computational challenges posed by modern data analysis. High-dimensional clustering methods are explicitly designed to directly address the task of grouping data points into clusters. Projection pursuit approaches were originally designed as powerful tools for exploring interesting projections and not explicitly assigning a unique cluster id to each event in the data set. Additionally, the high-dimensional nature of modern data instantly leads to a challenge of efficiently traversing myriads of low dimensional projections generated by the projection pursuit approach.

Here, we have integrated the principles of projection pursuit and clustering to create an automated projection pursuit clustering approach, designed to reveal noteworthy structures in high-dimensional data and assign cluster labels. The “projection” aspect of our approach involves orthogonal projection of high-dimensional data into a two-dimensional space. The “pursuit” component is guided by the concept of, at each step, identifying a projection with the smallest data density distribution along the decision boundary. The automated clustering aspect is executed through the recursive and exhaustive application of the “projection” and “pursuit” steps. To facilitate the computational challenge associated with exploring a high number of low-dimensional projections, we reduced the number of operations required by the algorithm at each recursive step by adopting the calculation of the optimal number of histogram bins. We also implemented parallelization of the APP algorithm enhancing its computational performance. Future improvements may focus on additionally optimizing specific algorithm components and/or exploring alternative computational strategies such as leveraging high-performance computing when a relatively large amount of high-dimensional data is needed to be processed.

In general, projection pursuit clustering can offer advantages in certain scenarios compared to traditional high-dimensional clustering methods. Projection pursuit can be effective when certain dimensions or variables are more important for clustering than others. It actively seeks projections that highlight important features, helping to focus on relevant dimensions and potentially improving cluster separation.

In scenarios characterized by sparse or imbalanced data, where there is a high degree of sparsity amid more abundant populations (which can extend to situations involving outliers and/or noise), projection pursuit proves beneficial in identifying pertinent dimensions and enhancing cluster separation. Traditional methods may face challenges in handling sparsity due to a dearth of informative features. Projection pursuit seeks to discover projections that are not only conducive to clustering but also interpretable. If the objective is to extract meaningful insights from clustering outcomes and comprehend the contributions of individual dimensions, projection pursuit may be the preferred choice.

The approach of discovering patterns through projection pursuit and clustering proves to be versatile across various data modalities. In this context, our focus was specifically on high-dimensional biological data given its frequent representation of both abundant and sparse populations within the same sample. We demonstrated the APP method’s capability to replicate experimentally validated cell type definitions, emphasizing the biological relevance of the clusters identified by our approach. We conducted a performance comparison of APP with other widely adopted clustering methods. While, on the whole, APP’s results align well with other methods, there are instances where one method may outperform the other, and we illustrated these nuances using real-world datasets.

In many biological real-world datasets, the availability of a clear “ground truth” can be challenging. As illustrated in the examples presented, reliance on domain experts’ knowledge-driven clustering or clustering done with widely-adopted approaches serves as a substitute for ground truth. While expert-driven clustering provides a valuable reference point, a more accurate (albeit labor-intensive) method for assessing clustering performance involves conducting functional tests on groups of cells assigned to the same cluster. By observing the functional “purity” and homogeneity of a given cluster compared to other cell clusters in the sample, researchers can achieve a more precise evaluation of the clustering results.

In one of such examples we used a data set with functionally validated ground truth and demonstrated that APP was able to effectively recapitulate experimentally validated cell-type definitions better than other widely used clustering approaches. We also presented APP’s utility in discovering new biologically meaningful patterns. By combining the strengths of LLMs for sequence analysis with the interpretability provided by clustering, we gained deeper insights into the complex relationships within TCR-pMHC sequence datasets. This integrative approach contributes to the elucidation of patterns that may have important implications for better understanding of TCR and pMHC function and design. Our results also underline the ongoing challenges with the design of universal pMHC-TCR specificity machine-learning models when showing that the pMHC-TCR recognition patterns are not always self-evident.

### Data availability

Flow cytometry data for six healthy controls and six COVID-19 patient samples is available upon request.

Ten healthy control cytof data is available at https://flowrepository.org/id/FR-FCM-Z24F.

scRNAseq PBMC dataset is available through the 10X Genomics website: https://cf.10xgenomics.com/samples/cell/pbmc3k/pbmc3k_filtered_gene_bc_matrices.ta r.gz.

Wildtype and 5XFAD mouse model scRNEseq data is available for download from the Gene Expression Omnibus (GEO) with the accession number GSE140510.

Human squamous cell carcinoma stained with SignalStar mIHC technology is available at https://data.mendeley.com/datasets/5vfz9vhm2s/1

TCR repertoire data is available as part of the McPAS-TCR database [http://friedmanlab.weizmann.ac.il/McPAS-TCR/]

### Code availability

Source code is available at https://github.com/cellsignal/projectionpursuit

## References

1. Friedman, J. H., & Tukey, J. W. (1974). A Projection Pursuit Algorithm for Exploratory Data Analysis. In IEEE Transactions on Computers: Vol. C–23 (Issue 9, pp. 881–890). 10.1109/t-c.1974.224051

2. Friedman, J. H. & Stuetzle, W. (1982). Projection pursuit methods for data analysis, in Modern Data Analysis, R.L., Launer & A.F., Siegel, eds, Academic Press (pp. 123–147).

3. Hastie, T., Tibshirani, R., & Friedman, J. (2009). The Elements of Statistical Learning: Data Mining, Inference, and Prediction (2nd ed.). Stanford University Press.

4. Bellman, R. E. & Rand Corporation (1957). Dynamic programming. Princeton University Press (pp. ix.) ISBN 978-0-691-07951-6.

5. Bellman, R. E. (1961). Adaptive control processes: a guided tour. Princeton University Press. ISBN 9780691079011.

6. Orlova, D. Y., Herzenberg, L. A., & Walther, G. (2017). Science not art: statistically sound methods for identifying subsets in multi-dimensional flow and mass cytometry data sets. In Nature Reviews Immunology (Vol. 18, Issue 1, pp. 77–77). 10.1038/nri.2017.150

7. Meehan, S., Kolyagin, G. A., Parks, D., Youngyunpipatkul, J., Herzenberg, L. A., Walther, G., Ghosn, E. E. B., & Orlova, D. Y. (2019). Automated subset identification and characterization pipeline for multidimensional flow and mass cytometry data clustering and visualization. In Communications Biology (Vol. 2, Issue 1). 10.1038/s42003-019-0467-6

8. Feynman, R., Leighton, R. B., & Sands, M. (1963). The Feynman Lectures on Physics Vol. II Ch. 19: The Principle of Least Action. Addison–Wesley.

9. Cook, D., Buja, A., Cabrera, J., & Hurley, C. (1995). Grand Tour and Projection Pursuit. In Journal of Computational and Graphical Statistics (Vol. 4, Issue 3, p. 155). JSTOR. 10.2307/1390844

10. Eddins, D. J., Yang, J., Kosters, A., Giacalone, V. D., Pechuan-Jorge, X., Chandler, J. D., Eum, J., Babcock, B. R., Dobosh, B. S., Hernández, M. R., Abdulkhader, F., Collins, G. L., Orlova, D. Y., Ramonell, R. P., Sanz, I., Moussion, C., Eun-Hyung Lee, F., Tirouvanziam, R. M., & Ghosn, E. E. B. (2023). Transcriptional reprogramming of infiltrating neutrophils drives lung pathology in severe COVID-19 despite low viral load. In Blood Advances (Vol. 7, Issue 5, pp. 778–799). American Society of Hematology. 10.1182/bloodadvances.2022008834

11. >Toghi Eshghi, S., Au-Yeung, A., Takahashi, C., Bolen, C. R., Nyachienga, M. N., Lear, S. P., Green, C., Mathews, W. R., & O’Gorman, W.E. (2019). Quantitative Comparison of Conventional and t-SNE-guided Gating Analyses. In Frontiers in Immunology (Vol. 10). 10.3389/fimmu.2019.01194

12. Zhou, Y., Song, W. M., Andhey, P. S., Swain, A., Levy, T., Miller, K. R., Poliani, P. L., Cominelli, M., Grover, S., Gilfillan, S., Cella, M., Ulland, T. K., Zaitsev, K., Miyashita, A., Ikeuchi, T., Sainouchi, M., Kakita, A., Bennett, D. A., Schneider, J. A., … Colonna, M. (2020). Human and mouse single-nucleus transcriptomics reveal TREM2-dependent and TREM2-independent cellular responses in Alzheimer’s disease. In Nature Medicine (Vol. 26, Issue 1, pp. 131–142). 10.1038/s41591-019-0695-9

13. Tickotsky, N., Sagiv, T., Prilusky, J., Shifrut, E., & Friedman, N. (2017). McPAS-TCR: a manually curated catalogue of pathology-associated T cell receptor sequences. In J. Wren (Ed.), Bioinformatics (Vol. 33, Issue 18, pp. 2924–2929). Oxford University Press (OUP). 10.1093/bioinformatics/btx286

14. Bankhead, P., Loughrey, M. B., Fernández, J. A., Dombrowski, Y., McArt, D. G., Dunne, P. D., McQuaid, S., Gray, R. T., Murray, L. J., Coleman, H. G., James, J. A., Salto-Tellez, M., & Hamilton, P. W. (2017). QuPath: Open source software for digital pathology image analysis. In Scientific Reports (Vol. 7, Issue 1). 10.1038/s41598-017-17204-5

15. Lin, Z., Akin, H., Rao, R., Hie, B., Zhu, Z., Lu, W., Smetanin, N., Verkuil, R., Kabeli, O., Shmueli, Y., dos Santos Costa, A., Fazel-Zarandi, M., Sercu, T., Candido, S., & Rives, A. (2022). Evolutionary-scale prediction of atomic level protein structure with a language model. Cold Spring Harbor Laboratory. 10.1101/2022.07.20.500902

16. Krissinel, E., & Henrick, K. (2007). Inference of Macromolecular Assemblies from Crystalline State. In Journal of Molecular Biology (Vol. 372, Issue 3, pp. 774–797). 10.1016/j.jmb.2007.05.022

17. Calinski, T., & Harabasz, J. (1974). A dendrite method for cluster analysis. In Communications in Statistics - Theory and Methods (Vol. 3, Issue 1, pp. 1–27). 10.1080/03610927408827101

18. Stringer, C., Wang, T., Michaelos, M., & Pachitariu, M. (2020). Cellpose: a generalist algorithm for cellular segmentation. In Nature Methods (Vol. 18, Issue 1, pp. 100–106). 10.1038/s41592-020-01018-x

19. Mann, H. B. and Wald, A. (1942) On the Choice of the Number of Class Intervals in the Application of Chi-Square Test. In Annals of Mathematical Statistics (Vol. 13, Issue 3, pp. 306–317).

20. Ben-Hur, A. et al. (2001). Support Vector Clustering. In Journal of Machine Learning Research (Vol. 2, pp. 125–137).

21. Orlova, D. Y., Meehan, S., Parks, D., Moore, W. A., Meehan, C., Zhao, Q., Ghosn, E. E. B., Herzenberg, L. A., & Walther, G. (2018). QFMatch: multidimensional flow and mass cytometry samples alignment. In Scientific Reports (Vol. 8, Issue 1). 10.1038/s41598-018-21444-4

22. Zimmerman, N. (2011). A computational approach to identification and comparison of cell subsets in flow cytometry data. Ph.D. Thesis, Stanford University. Available: https://stacks.stanford.edu/file/druid:hg137hq6178/Zimmerman-Dissertation-v2-augmented.pdf.

23. Xu, M. M., Pu, Y., Han, D., Shi, Y., Cao, X., Liang, H., Chen, X., Li, X.-D., Deng, L., Chen, Z. J., Weichselbaum, R. R., & Fu, Y.-X. (2017). Dendritic Cells but Not Macrophages Sense Tumor Mitochondrial DNA for Cross-priming through Signal Regulatory Protein α Signaling. In Immunity (Vol. 47, Issue 2, pp. 363–373.e5). 10.1016/j.immuni.2017.07.016

24. Dutertre, C.-A., Becht, E., Irac, S. E., Khalilnezhad, A., Narang, V., Khalilnezhad, S., Ng, P. Y., van den Hoogen, L. L., Leong, J. Y., Lee, B., Chevrier, M., Zhang, X. M., Yong, P. J. A., Koh, G., Lum, J., Howland, S. W., Mok, E., Chen, J., Larbi, A., … Ginhoux, F. (2019). Single-Cell Analysis of Human Mononuclear Phagocytes Reveals Subset-Defining Markers and Identifies Circulating Inflammatory Dendritic Cells. In Immunity (Vol. 51, Issue 3, pp. 573–589.e8). 10.1016/j.immuni.2019.08.008

25. Bourdely, P., Anselmi, G., Vaivode, K., Ramos, R. N., Missolo-Koussou, Y., Hidalgo, S., Tosselo, J., Nuñez, N., Richer, W., Vincent-Salomon, A., Saxena, A., Wood, K., Lladser, A., Piaggio, E., Helft, J., & Guermonprez, P. (2020). Transcriptional and Functional Analysis of CD1c+ Human Dendritic Cells Identifies a CD163+ Subset Priming CD8+CD103+ T Cells. In Immunity (Vol. 53, Issue 2, pp. 335–352.e8). 10.1016/j.immuni.2020.06.002

26. Comi, M., Avancini, D., Santoni de Sio, F., Villa, M., Uyeda, M. J., Floris, M., Tomasoni, D., Bulfone, A., Roncarolo, M. G., & Gregori, S. (2019). Coexpression of CD163 and CD141 identifies human circulating IL-10-producing dendritic cells (DC-10). In Cellular & Molecular Immunology (Vol. 17, Issue 1, pp. 95–107). 10.1038/s41423-019-0218-0

27. Sedighzadeh, S. S., Khoshbin, A. P., Razi, S., Keshavarz-Fathi, M., & Rezaei, N. (2021). A narrative review of tumor-associated macrophages in lung cancer: regulation of macrophage polarization and therapeutic implications. In Translational Lung Cancer Research (Vol. 10, Issue 4, pp. 1889–1916). 10.21037/tlcr-20-1241

28. Laviron, M., Petit, M., Weber-Delacroix, E., Combes, A. J., Arkal, A. R., Barthélémy, S., Courau, T., Hume, D. A., Combadière, C., Krummel, M. F., & Boissonnas, A. (2022). Tumor-associated macrophage heterogeneity is driven by tissue territories in breast cancer. In Cell Reports (Vol. 39, Issue 8, p. 110865). 10.1016/j.celrep.2022.110865

29. Miura, Y., Anami, T., Yatsuda, J., Motoshima, T., Oka, S., Suyama, K., Inoshita, N., Kinowaki, K., Urakami, S., Kamba, T., & Komohara, Y. (2021). HLA-DR and CD74 Expression and the Immune Microenvironment in Renal Cell Carcinoma. In Anticancer Research (Vol. 41, Issue 6, pp. 2841–2848). 10.21873/anticanres.15065

30. Senosain, M.-F., Zou, Y., Novitskaya, T., Vasiukov, G., Balar, A. B., Rowe, D. J., Doxie, D. B., Lehman, J. M., Eisenberg, R., Maldonado, F., Zijlstra, A., Novitskiy, S. V., Irish, J. M., & Massion, P. P. (2021). HLA-DR cancer cells expression correlates with T cell infiltration and is enriched in lung adenocarcinoma with indolent behavior. In Scientific Reports (Vol. 11, Issue 1). 10.1038/s41598-021-93807-3

31. Madani, A., Krause, B., Greene, E. R., Subramanian, S., Mohr, B. P., Holton, J. M., Olmos, J. L., Jr., Xiong, C., Sun, Z. Z., Socher, R., Fraser, J. S., & Naik, N. (2023). Large language models generate functional protein sequences across diverse families. In Nature Biotechnology (Vol. 41, Issue 8, pp. 1099–1106). 10.1038/s41587-022-01618-2

32. Chandra, A., Tünnermann, L., Löfstedt, T., & Gratz, R. (2023). Transformer-based deep learning for predicting protein properties in the life sciences. In eLife (Vol. 12). 10.7554/elife.82819

33. Ruffolo, J. A., & Madani, A. (2024). Designing proteins with language models. In Nature Biotechnology (Vol. 42, Issue 2, pp. 200–202). 10.1038/s41587-024-02123-4

34. Quintana, F., Treangen, T., & Kavraki, L. (2023). Leveraging Large Language Models for Predicting Microbial Virulence from Protein Structure and Sequence. In Proceedings of the 14th ACM International Conference on Bioinformatics, Computational Biology, and Health Informatics. BCB ’23: 14th ACM International Conference on Bioinformatics, Computational Biology, and Health Informatics. ACM. 10.1145/3584371.3612953

35. Leem, J., Mitchell, L. S., Farmery, J. H. R., Barton, J., & Galson, J. D. (2022). Deciphering the language of antibodies using self-supervised learning. In Patterns (Vol. 3, Issue 7, p. 100513). 10.1016/j.patter.2022.100513

36. Wu, K., Yost, K. E., Daniel, B., Belk, J. A., Xia, Y., Egawa, T., Satpathy, A., Chang, H. Y., & Zou, J. (2021). TCR-BERT: learning the grammar of T-cell receptors for flexible antigen-xbinding analyses. Cold Spring Harbor Laboratory. 10.1101/2021.11.18.469186

37. Kruskal, J. B. (1969). “Toward a practical method which helps uncover the structure of a set of observations by finding the line transformation which optimizes a new “index of condensation.”” Milton, RC, & Nelder, JA (eds), Statistical computation; New York, Academic Press (pp. 427–440).

38. Huber, P. J. (1985). Projection Pursuit. In The Annals of Statistics (Vol. 13, Issue 2). Institute of Mathematical Statistics. 10.1214/aos/1176349519

