## Supplementary Figures for "Lifting the curse from high dimensional data: Automated projection pursuit clustering for the variety of biological data modalities"

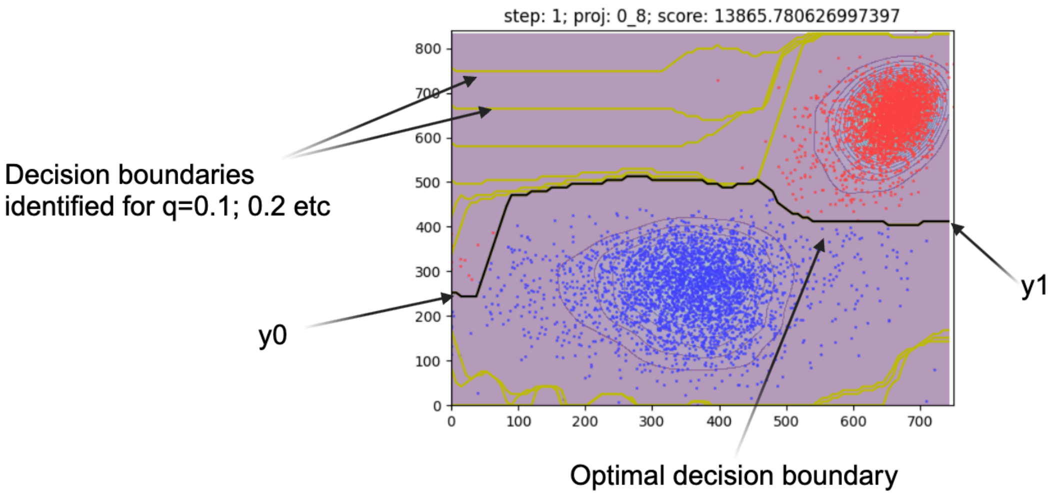


**Supplementary Figure 1. Example of an optimal decision boundary search for one of the 2D data projections.**

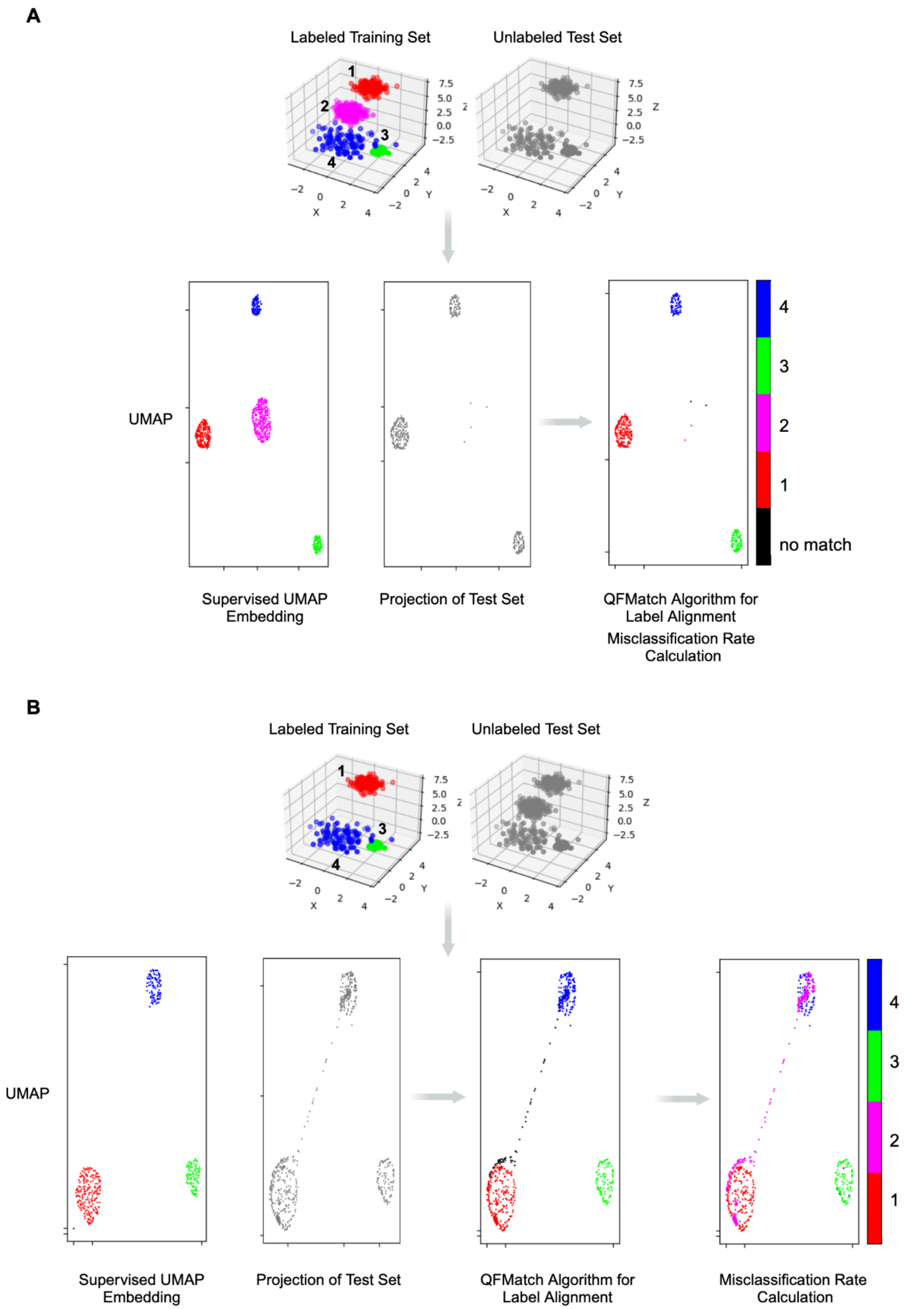
A
Labeled Training Set
Unlabeled Test Set
1
2
3
4
7.5
5.0
2.5
0.0
−2.5
Z
X
Y
UMAP
4
3
2
1
no match
Supervised UMAP Embedding
Projection of Test Set
QFMatch Algorithm for Label Alignment
Misclassification Rate Calculation
B
Labeled Training Set
Unlabeled Test Set
1
3
4
UMAP
4
3
2
1
Supervised UMAP Embedding
Projection of Test Set
QFMatch Algorithm for Label Alignment
Misclassification Rate Calculation

**Supplementary Figure 2. Workflow for the automated label transfer across samples.** This pipeline utilizes labeled or partially labeled training samples with the marker expression data and ground truth cluster labels. It applies UMAP to learn a distance metric that optimally separates classes while preserving relationships in the marker space. And then projects the unlabeled test set into the UMAP embedding space built using the training set. It further relies on the QFMatch algorithm to align cluster labels between the test set and the training set for the downstream calculation for the number of misclassified events per cluster id. Label transfer pipeline allows quantitatively comparing and aligning cluster labels across training and test samples in cases where clusters may be absent in either the test (**A**) or training data (**B**). Though the latter is a harder case to handle.

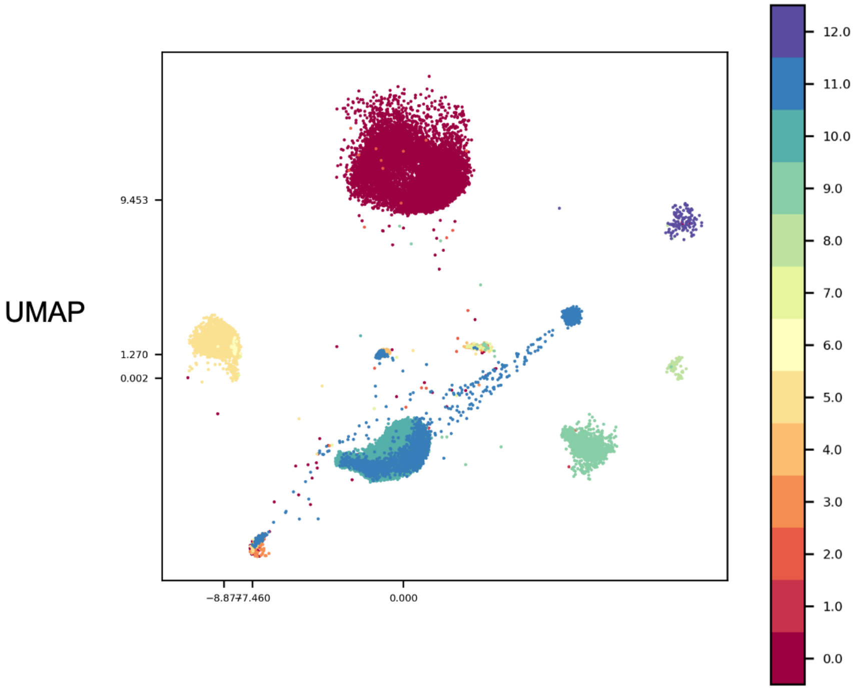


**Supplementary Figure 3. An example of a discrepancy between the data topology and clustering decisions.** As data topology suggests cluster 11 is more heterogeneous than was originally defined by the clustering approach (manual gating, in this case).

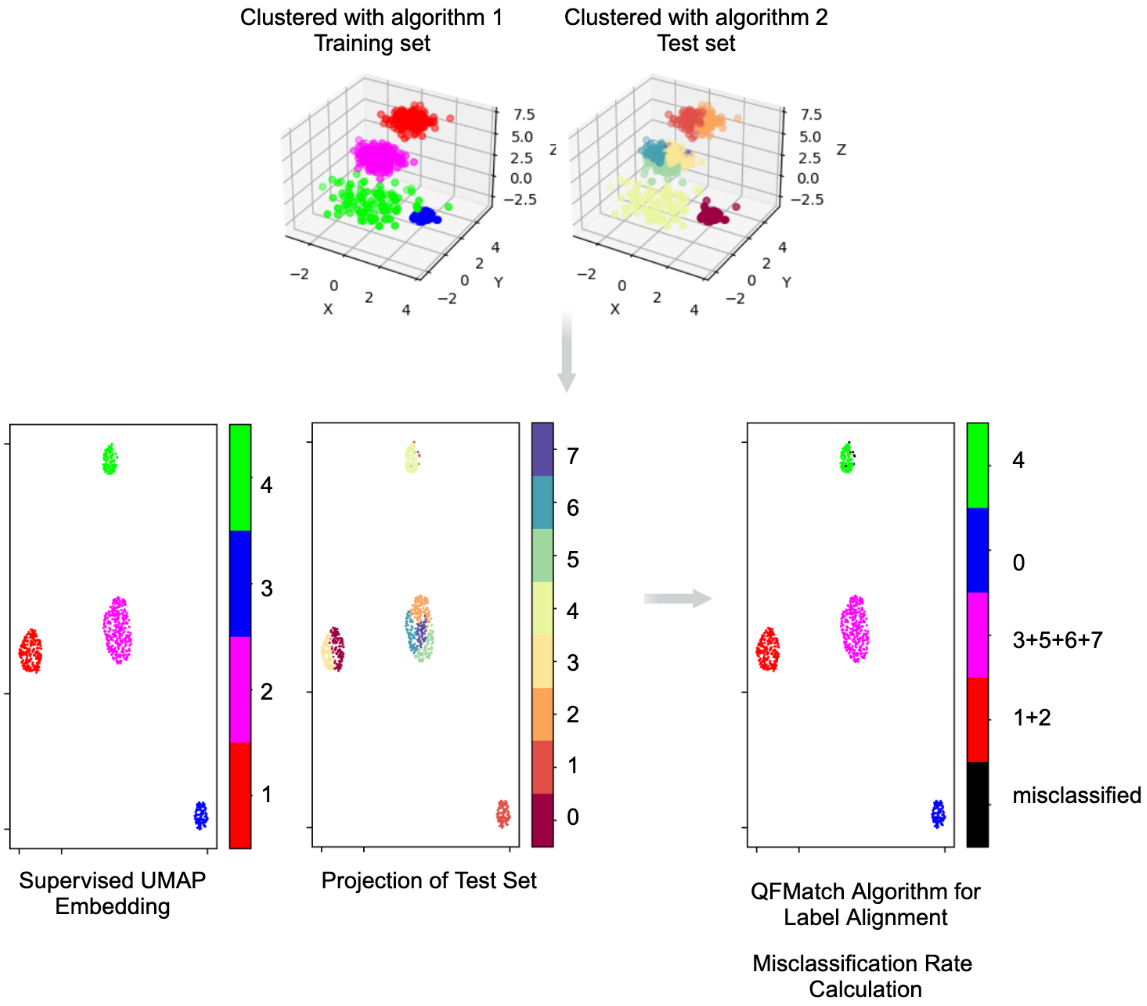


**Supplementary Figure 4. Cartoon representation of the label transfer pipeline application to quantitative comparison of two clustering algorithms decisions made on the same data set.**

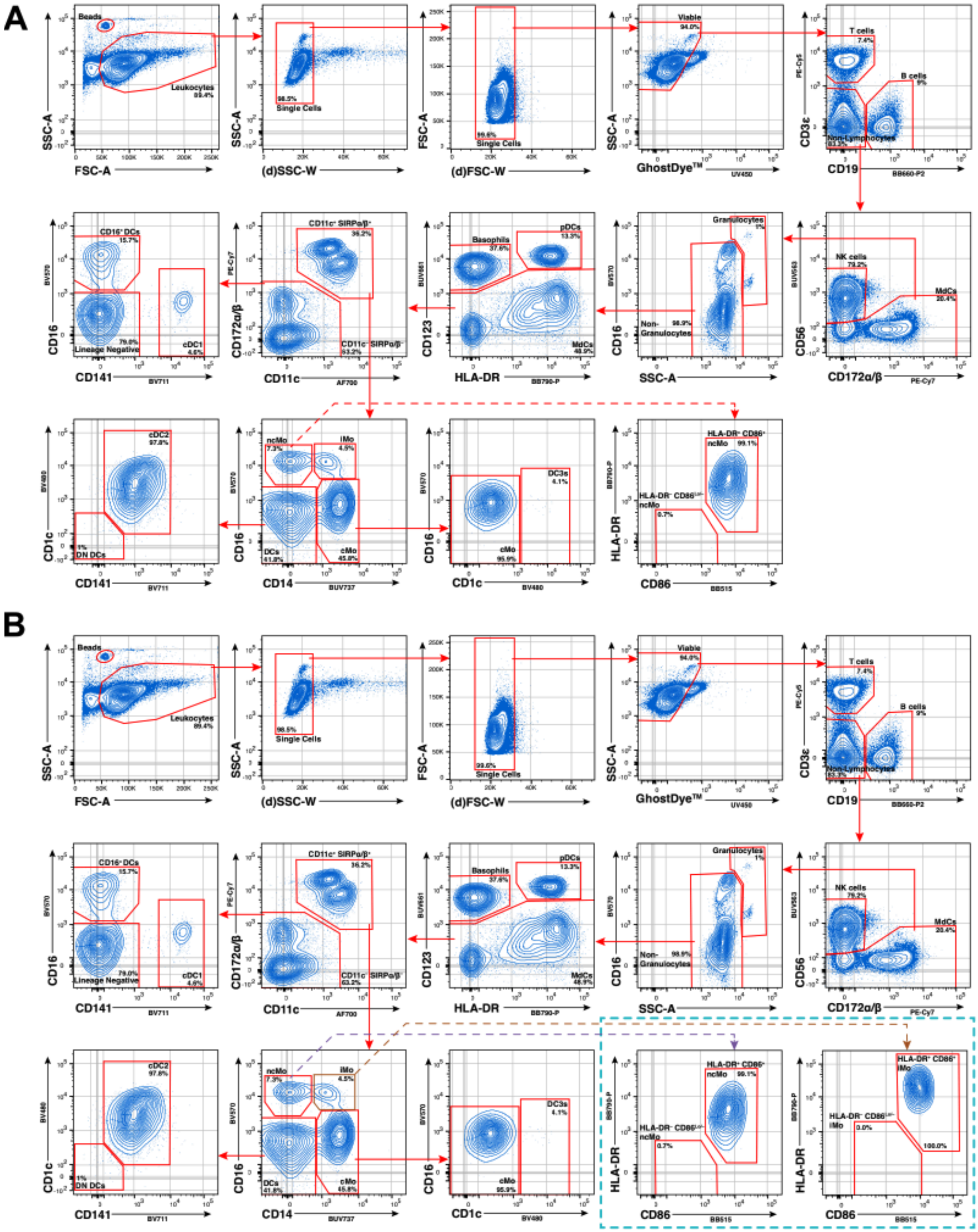


**Supplementary Figure 5. Expert-defined manual gating strategies used for high-dimensional flow cytometry data. A.** Representative gating strategy of a randomly selected health donor sample to interrogate myeloid-derived cells (MdCs) enriched from PBMCs (see methods). **B.** Original gating strategy from (A) was refined following the label transfer pipeline “sanity check”, which revealed additional heterogeneity with the intermediate monocyte (iMo) population of COVID-19 patients that was not ubiquitously present in health donors, similar to non-classical monocytes (ncMo; see blue box & Figure 3G).

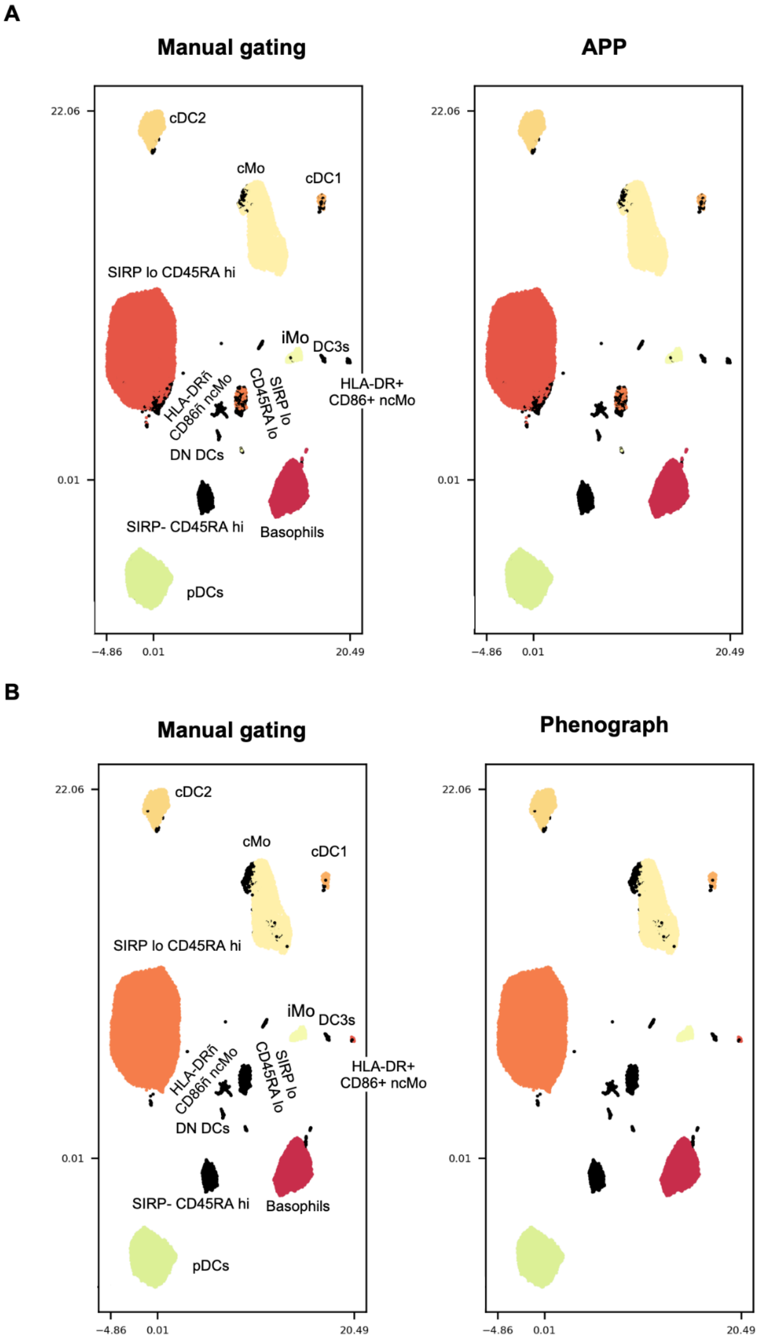


**Supplementary Figure 6. An illustration of misclassification by APP (A) and Phenograph (B) compared to manually gated cell population annotations.** Misclassified events, calculated using the automated label transfer pipeline, are highlighted in black.

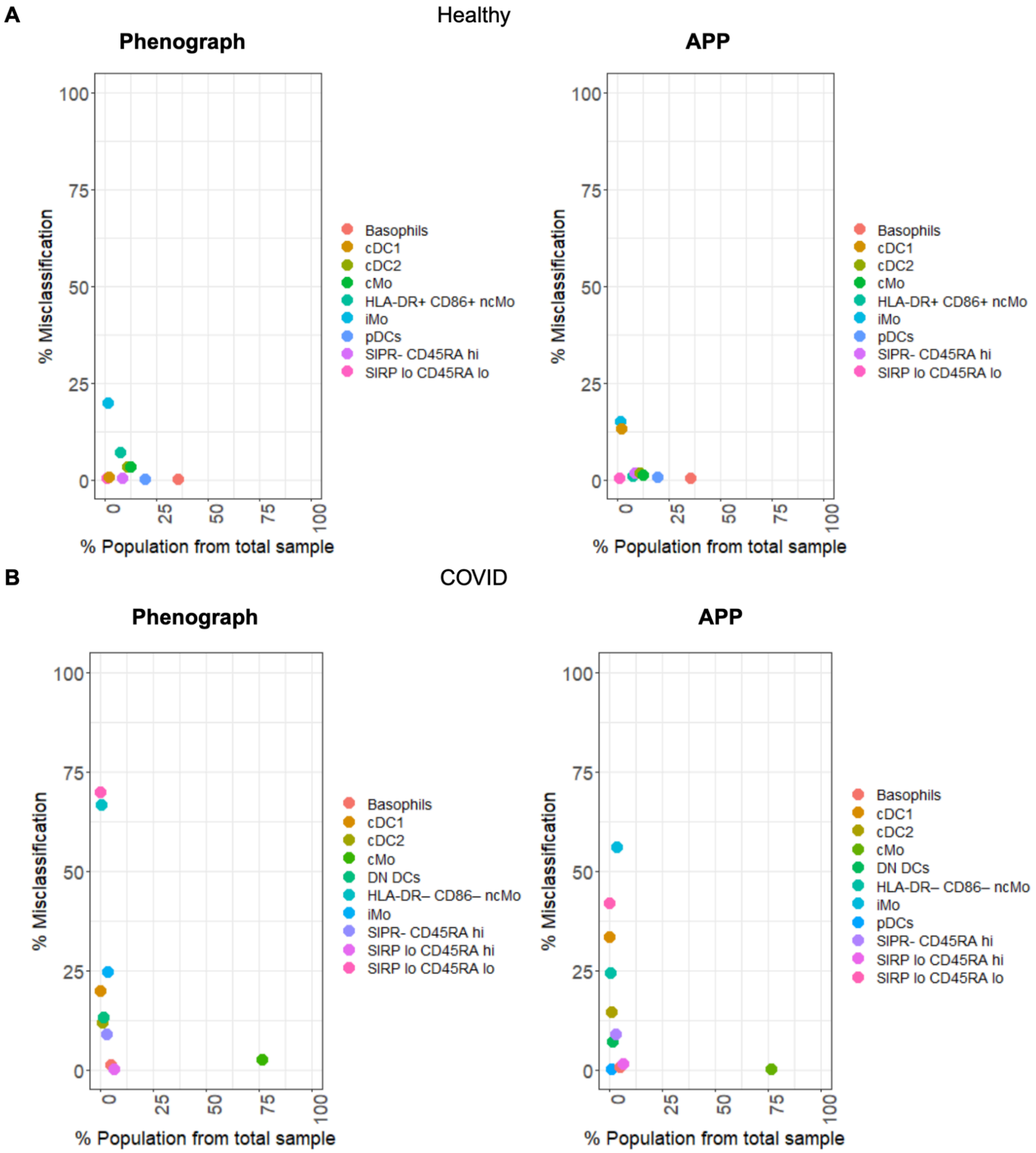


**Supplementary Figure 7. Smaller cell populations are more prone to misclassification by clustering algorithms in both healthy donor (A) and COVID-19 patient samples (B).**

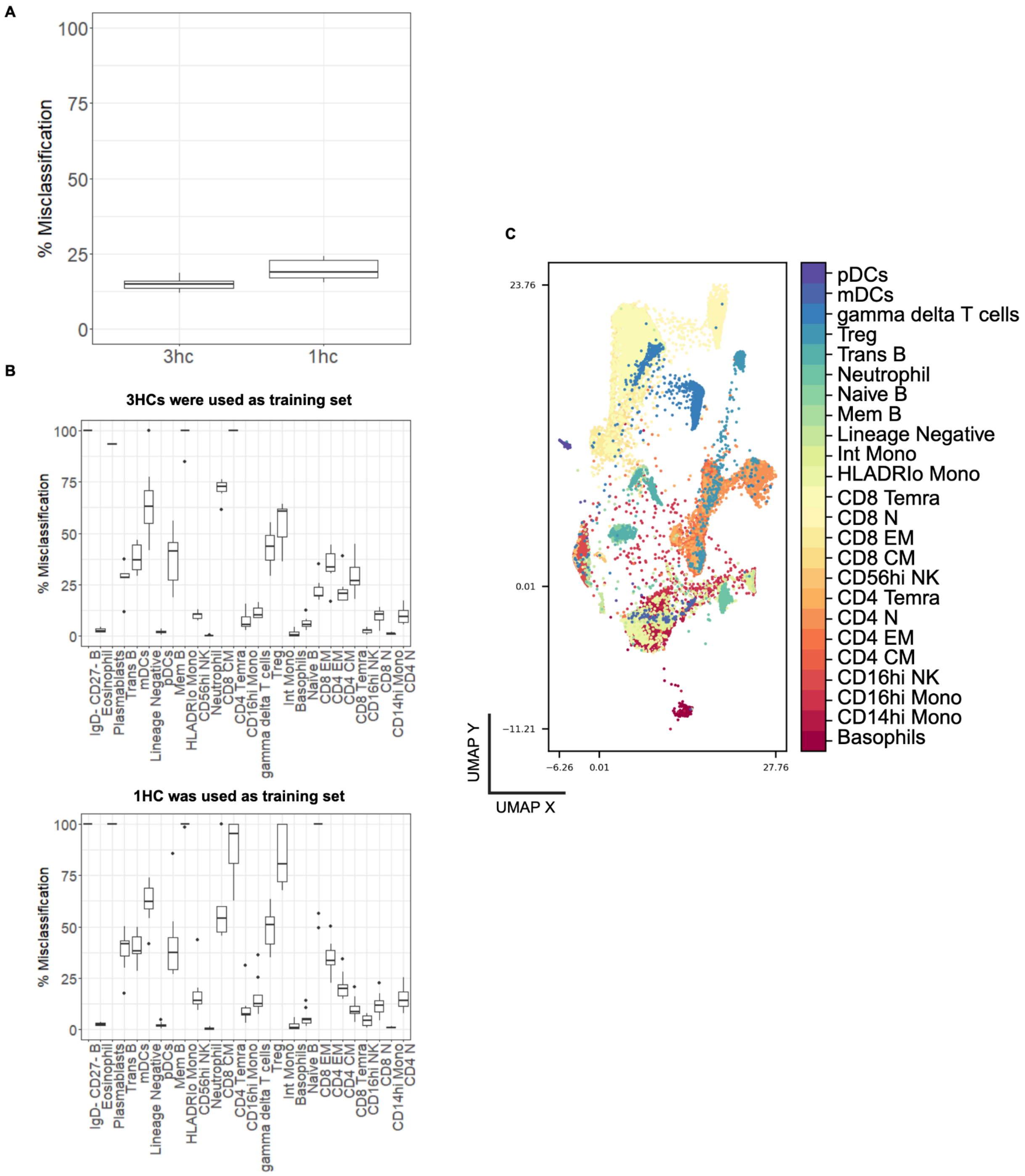


**Supplementary Figure 8. Discrepancies between the underlying data topology and the ground truth labels could negatively affect the performance of the label transfer pipeline. A.** Randomly chosen one and three (out of ten) healthy control (hc) PBMCs samples, characterized with the ~30 marker CYTOF panel, were used as training sets and the rest of samples were used as the test set. **B.** The primary source of misclassification arises from the more heterogeneous nature of cell populations than initially identified with the established expert-defined manual gating strategy. For instance, the gamma delta T cells population on panel **C** is actually distributed between the two clusters, as data topology suggests.

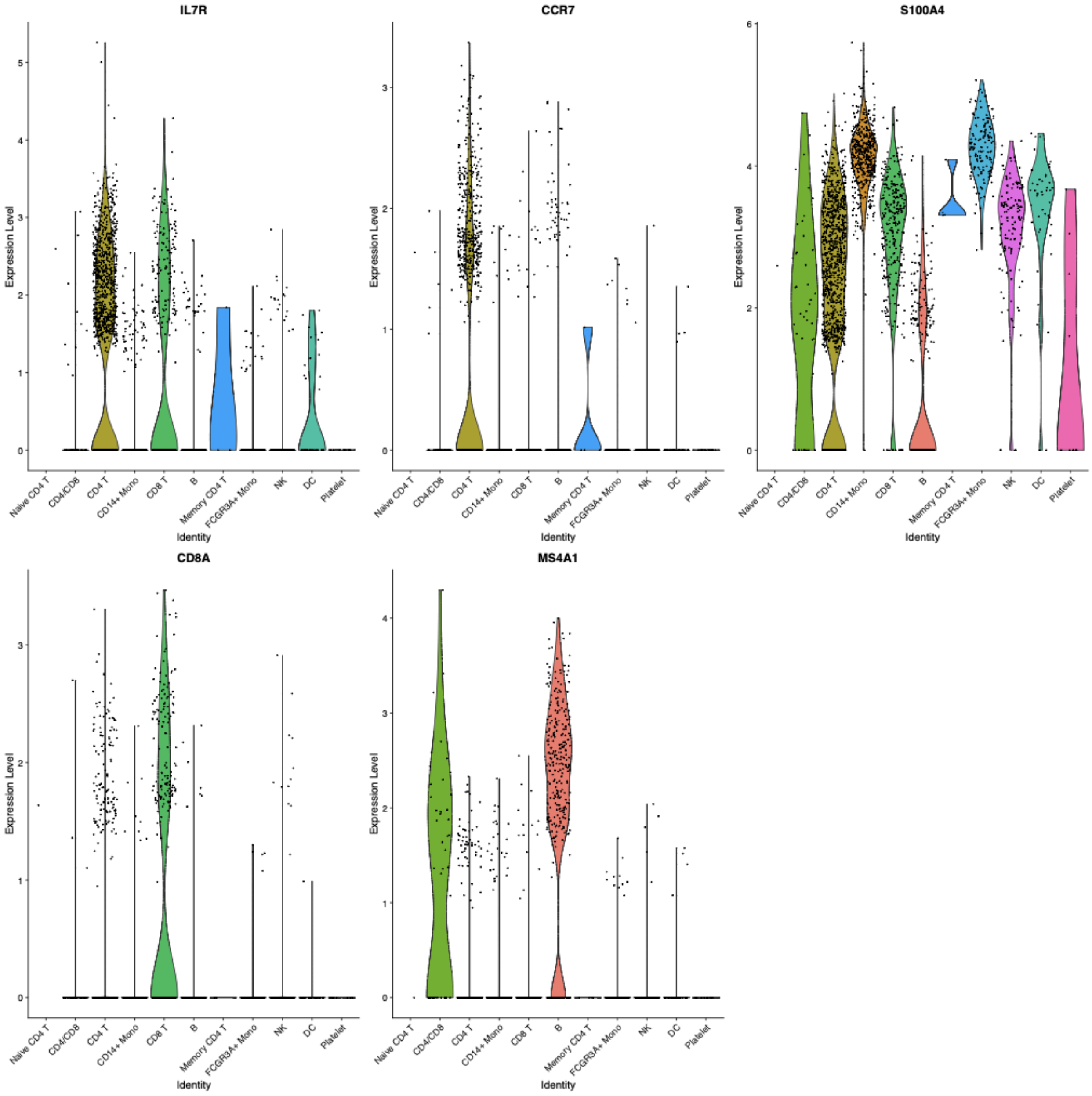


**Supplementary Figure 9. The expression of B and T marker genes in cell subsets identified within the PBMCs dataset.** MS4A1 is a B cell specific marker, the rest are CD4 and CD8 T cell specific markers. The group of cells that was misclassified by APP is here labeled "CD4/CD8".

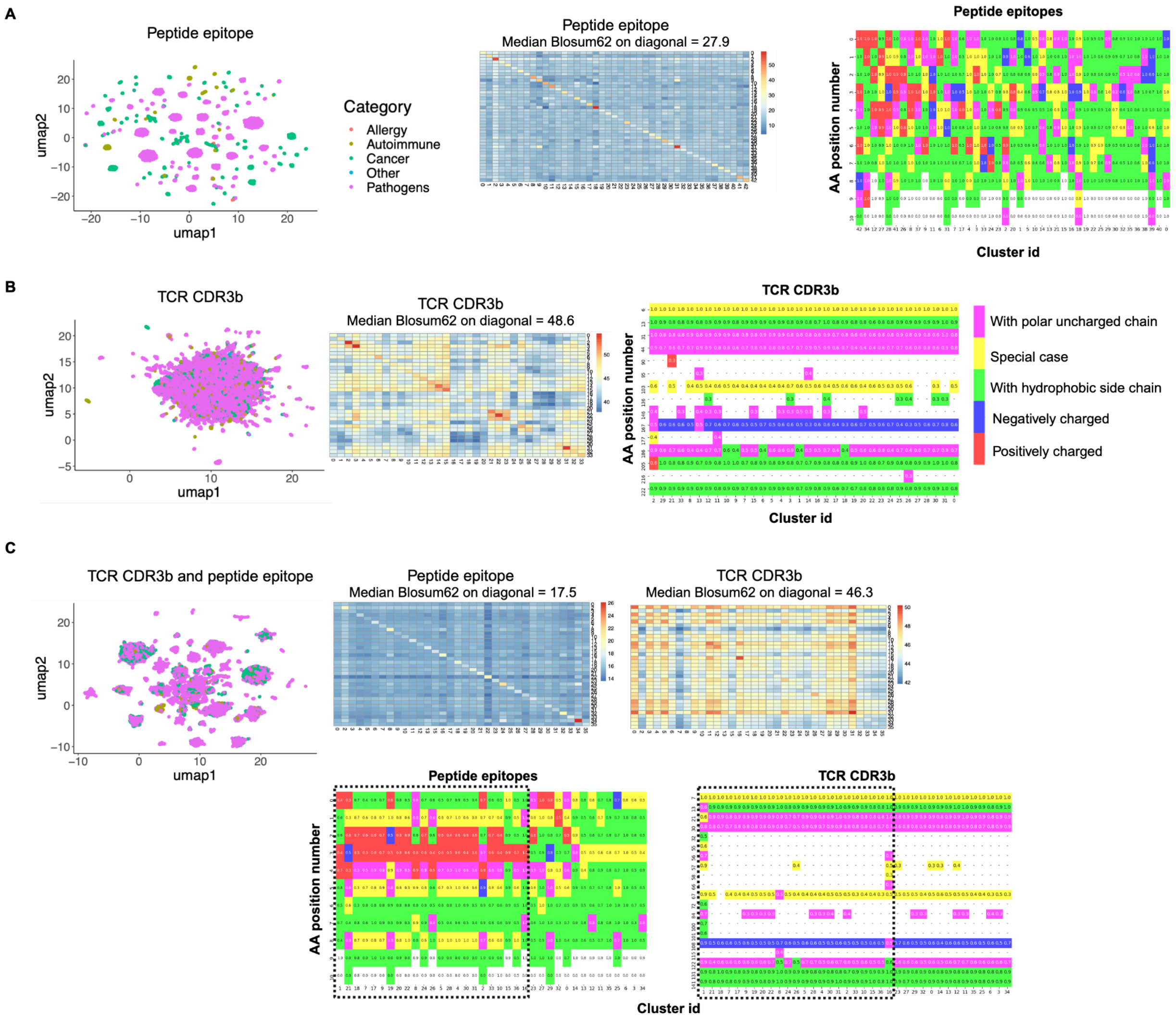


**Supplementary Figure 10. Sequence similarity and amino acid R group properties analysis of ESM embeddings generated for peptide epitope sequences (A), TCR CDR3b sequences (B), and the concatenation of both TCR and peptide embeddings (C).** Amino acids at a specific position within a designated cluster were categorized according to their R group properties (as indicated in the inserted legend). The most prevalent property within each group is presented, along with its corresponding probability.

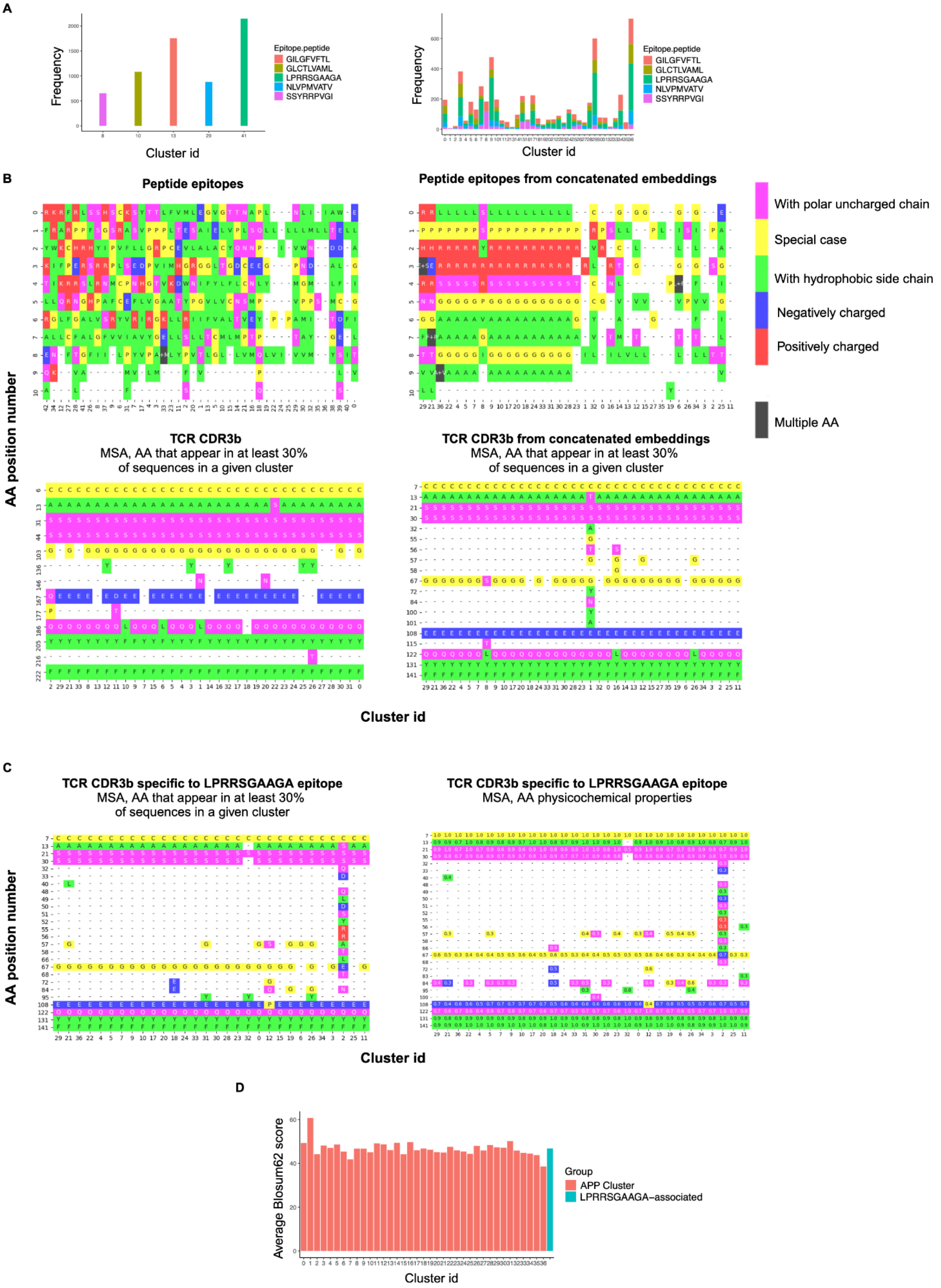
A
Frequency
Cluster id
Epitope.peptide
GILGFVFTL
GLCTLVAML
LPRRSGAAGA
NLVPMVATV
SSYRRPVGI
B
Peptide epitopes
Peptide epitopes from concatenated embeddings
With polar uncharged chain
Special case
With hydrophobic side chain
Negatively charged
Positively charged
Multiple AA
AA position number
TCR CDR3b
MSA, AA that appear in at least 30%
of sequences in a given cluster
TCR CDR3b from concatenated embeddings
MSA, AA that appear in at least 30%
of sequences in a given cluster
Cluster id
C
TCR CDR3b specific to LPRRSGAAGA epitope
MSA, AA that appear in at least 30%
of sequences in a given cluster
TCR CDR3b specific to LPRRSGAAGA epitope
MSA, AA physicochemical properties
AA position number
Cluster id
D
Average Blosum62 score
Cluster id
Group
APP Cluster
LPRRSGAAGA–associated

**Supplementary Figure 11. No common binding motif is identified on TCR CDR3b, even when considering similar peptide epitopes. A.** The distribution of the five most common peptides (in the database utilized for this study) appears highly cluster-specific in the context of single-class embeddings. However, when using concatenated embeddings (CDR3b and peptide), these five peptides are dispersed across multiple clusters. **B.** No discernible binding motifs appear in the TCR CDR3b, whether analyzed individually (left side) or as concatenated embeddings (right side). Even when focusing exclusively on CDR3b sequences specific to the LPRRSGAAGA peptide (**C**), these sequences exhibit no notable increase in sequence similarity (**D**), as approximated by the Blosum62 score, when compared to the CDR3b sequences clustered together in the concatenated embeddings data presented in Figure 6.

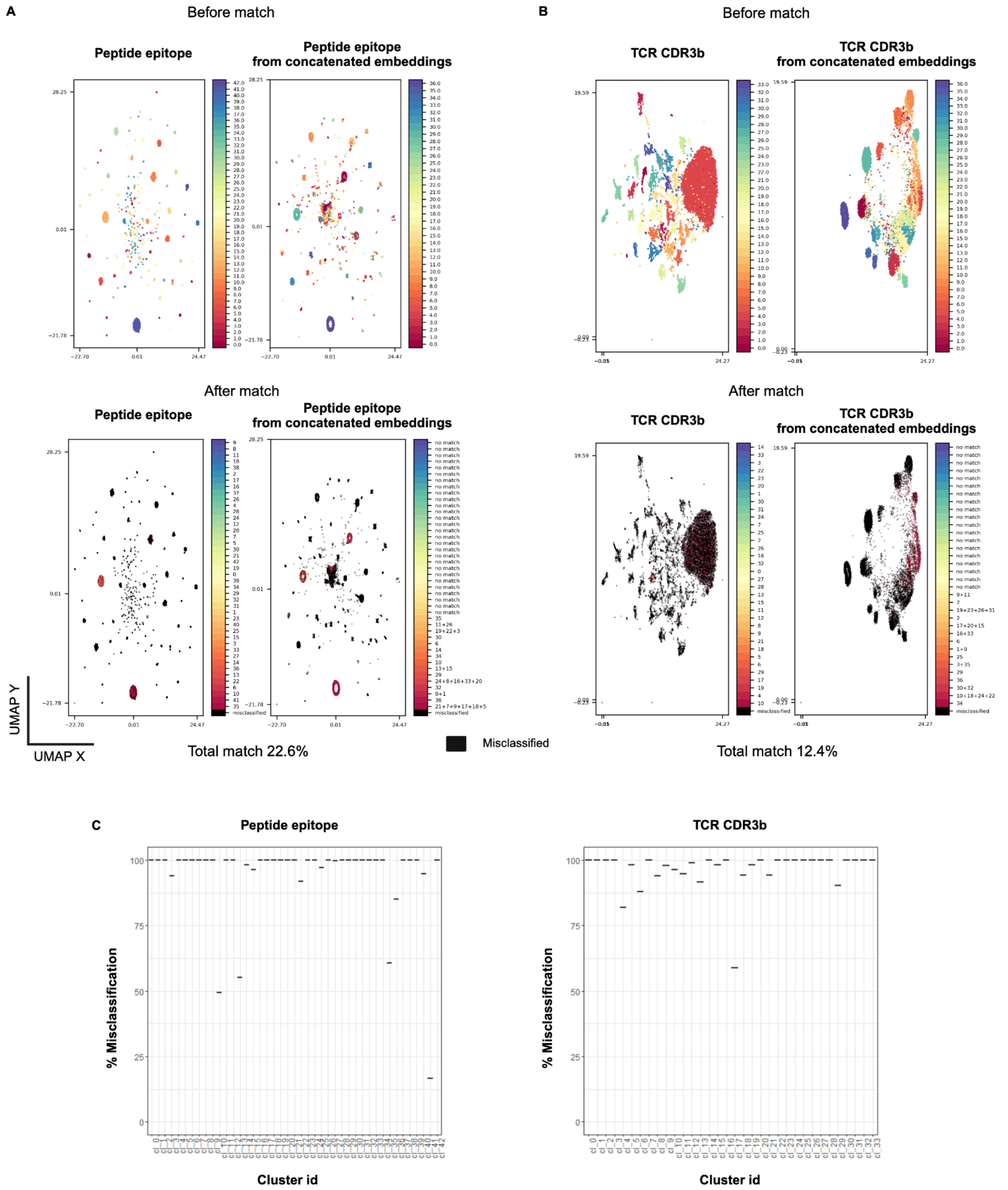


**Supplementary Figure 12. Concatenation of TCR CDR3b and peptide embeddings results in distinct similarity criteria captured by the ESM model.** The automated label transfer pipeline was employed to align clusters obtained for the same class of sequences (peptides (**A**) and TCR CDR3b (**B**)) generated from the single class embeddings and concatenated embeddings. The label transfer pipeline was executed on 30 PCs, using the APP cluster labels generated for the single class embeddings as the training set. On the top row of panels A and B, cluster ids are shown before the cluster alignment, and thus the same color may represent two unrelated clusters on the left and right UMAP plots for each class. On the bottom row of panels A and B, cluster ids are shown after the match/cluster alignment, and thus the same color represents aligned clusters within the same sequence class. Non-matched clusters are shown in black. **C.** Per-cluster id misclassification as estimated by the label transfer pipeline.

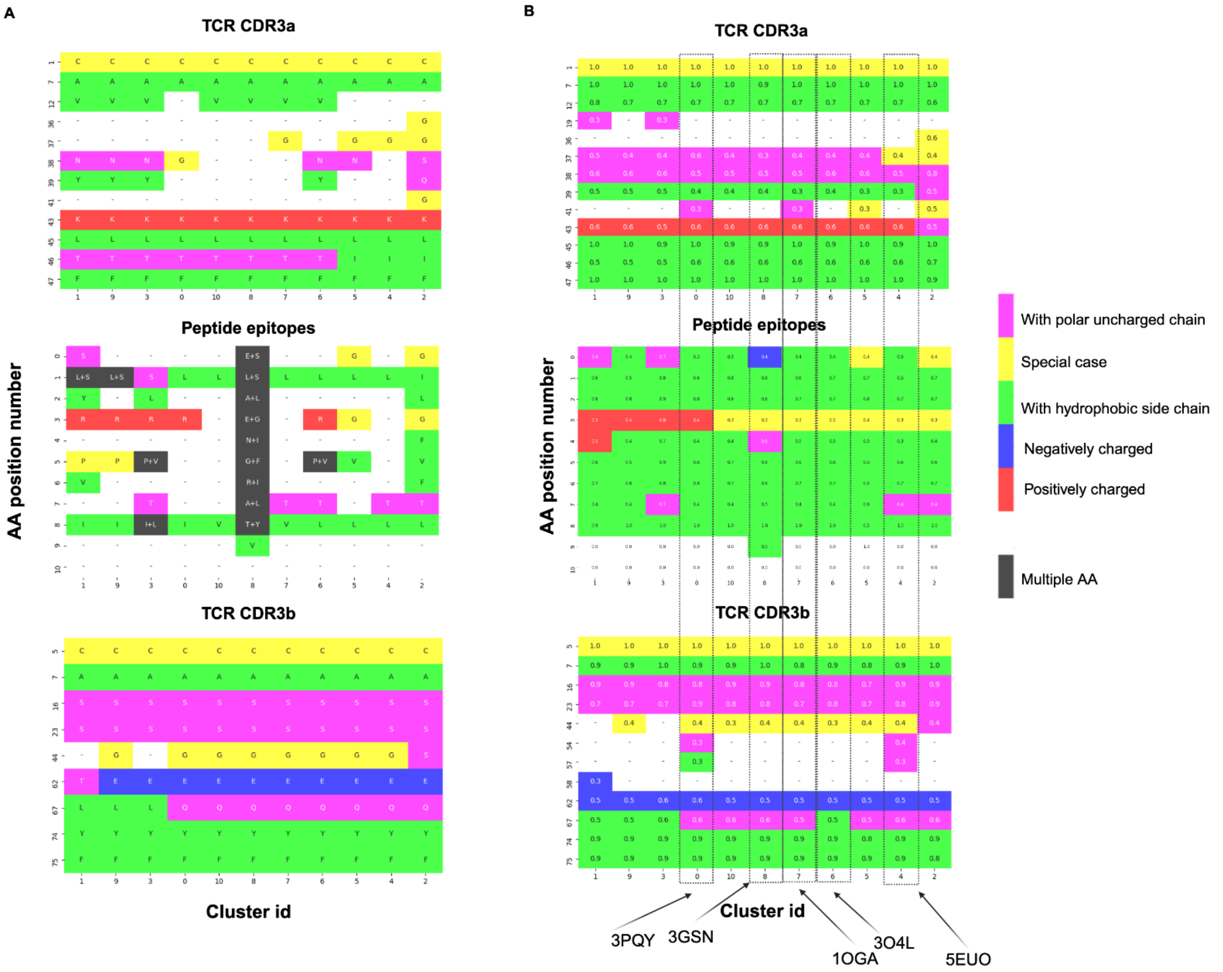


C

| PDB identifier | 1OGA | 3GSN | 3O4L | 5EUO | 3PQY |
|---|---|---|---|---|---|
| number of H-bonds with TCR-b | 5 | 4 | 1 | 4 | 3 |
| number of H-bonds with TCR-a | 2 | 0 | 2 | 3 | 3 |

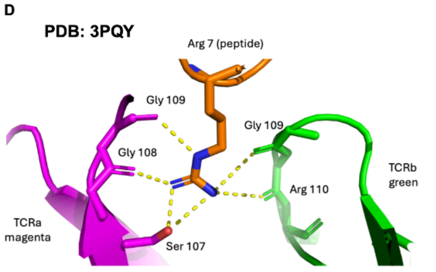


H bonds with TCRa involved:
lateral chain of Ser107; main chain Gly 108 and Gly 109

H bonds with TCRb involved:
main chain Gly 109 and Arg 110

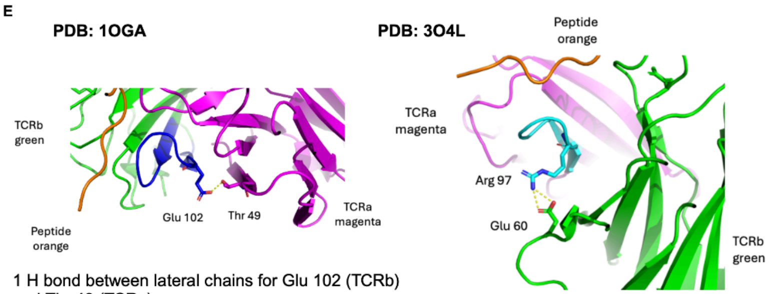


1 H bond between lateral chains for Glu 102 (TCRb) and Thr 49 (TCRa)

2 H bonds involved:
main chain Glu 60 and main chain Arg 97

**Supplementary Figure 13. The TCR CDR3b sequences exhibit an enrichment of negatively charged amino acids, while TCR CDR3a sequences are characterized**

**by an enrichment of positively charged amino acids. A.** Amino acids that appear with more than 40 percent probability (in TCR sequence) and more than 30 percent probability (in peptide sequence) in a given position and in a given cluster are color-coded based on their R group property. These physicochemical maps are constructed for the 4000 CDR3a-CDR3b-peptide data set presented on the right side of Figure 6. **B.** Amino acids at a specific position within a designated cluster were sorted into the corresponding R group property, as specified in the legend. The most dominant property within each group is displayed, along with its associated probability. The results presented focus on R group property classes that exhibit a prevalence of more than 30 percent at a given position within a designated cluster. Analysis of these properties for the peptides showed important hydrophobic enrichment in all clusters (middle panel right, green color). For five independent clusters, we found in the PDB five TCR-pMHC crystal structures and analyzed them. **C.** All showed that the antigen directly interacted with both, alpha- and beta- TCR units. Unexpectedly for such hydrophobic peptides was the presence of H-bonds, noticed for all of them (from 2 to 7 H-bonds) and involving both TCRab. In the center of peptides, we noticed either the presence of one positively charged amino acid (often Arg), or the presence of one Gly. **D.** Structure analysis of PDB_3PQY showed that Arg7 has its 3-nitrogens from the lateral chain involved in the formation of H-bonds with TCRa (4 H-bonds) and TCRb (2 H-bonds). We suggest that through the formation of all the H-bonds the peptide orients itself between the TCRs sub-units and complement the stable 3D surface of the TCRs. Such H-bonds also participate in the stability of the [peptide-TCRs] complex and the presence of Gly (B, middle panel right, yellow color) allows partial bending of the peptide, when necessary. **E.** We noticed for all CDR3a and CDR3b the respective presence of conserved Lys and Glu (A,B). Analysis of the previous 5 structures showed that these amino acids don't interact directly with the peptide, neither between themselves. They form H-bonds or ionic bonds between the two TCRs and through this mechanism participate in the pairing of the two subunits (see 1 H-bond between the conserved Glu102 (CDR3b) and Thr49 from TCRa; the ionic bonds bellow between Arg97 (CDR3a) and Glu60 (TCRb).

| | Laser | Detector | Antigen | Fluorophore | Clone | Isotype | Vendor | Cat. No. | Titration |
|---|---|---|---|---|---|---|---|---|---|
| | | B 488/10 | Side Scatter (SSC) | — | — | — | — | — | — |
| 1 | | B 515/20 | CD86 | BB515 | FUN-1 | Mouse IgG1, κ | BD Biosciences | 564544 | 1:10 |
| 2 | | B 610/20 | CD45-RA | BB630-P2 | HI-100 | Mouse IgG2a, κ | BD Biosciences | 624294 | 1:160 |
| 3 | 488 nm | B 670/30 | CD19 | BB660-P2 | HIB19 | Mouse IgG1, κ | BD Biosciences | 624295 | 1:50 |
| 4 | | B 710/50 | CD45 | BB700 | HI-30 | Mouse IgG1, κ | BD Biosciences | 746090 | 1:640 |
| 5 | | B 750/30 | CD4 | BB755-P | RPA-T4 | Mouse IgG1, κ | BD Biosciences | 624391 | 1:100 |
| 6 | | B 780/60 | HLA-DR (MHC II) | BB790-P | G46-6 | Mouse IgG2a, κ | BD Biosciences | 624296 | 1:50 |
| 7 | | V 431/28 | CD163 | BV421 | GHI/6I | Mouse IgG1, κ | BioLegend | 333612 | 1:50 |
| 8 | | V 470/15 | CD1c | BV480 | F10/21A3 | Mouse IgG1, κ | BD Biosciences | 746677 | 1:20 |
| 9 | | V 586/15 | CD16 | BV570 | 3G8 | Mouse IgG1, κ | BioLegend | 302035 | 1:100 |
| 10 | 405 nm | V 610/20 | CD169 | BV605 | 7-239 | Mouse IgG1, κ | BioLegend | 346010 | 1:100 |
| 11 | | V 670/30 | HLA-ABC (MHC-I) | BV650 | G46-2.6 | Mouse IgG1, κ | BD Biosciences | 740581 | 1:320 |
| 12 | | V 710/50 | CD141 | BV711 | 1A4 | Mouse IgG1, κ | BioLegend | 563155 | 1:100 |
| 13 | | V 740/35 | CD11b | BV750 | ICRF44 | Mouse IgG1, κ | BD Biosciences | 747357 | 1:20 |
| 14 | | V 780/60 | CD197 (CCR7) | BV785 | G043H7 | Mouse IgG2a, κ | BioLegend | 353230 | 1:50 |
| 15 | | YG 586/15 | XCR1 | PE | S15046E | Rat IgG2a, κ | BioLegend | 372604 | 1:50 |
| 16 | | YG 610/20 | CD206 | PE-Dazzle594 | 15-2 | Mouse IgG1, κ | BioLegend | 321130 | 1:200 |
| 17 | 561 nm | YG 670/30 | CD3ε | PE-Cy5 | UCHT1 | Mouse IgG1, κ | BioLegend | 555334 | 1:100 |
| 18 | | YG 710/50 | CD10 | PE-Cy5.5 | HI10a | Mouse IgG1, κ | Homemade | — | 1:40 |
| 19 | | YG 780/60 | CD172a/b (SIRPα/β1) | PE-Cy7 | SE5A5 | Mouse IgG1, κ | BioLegend | 323808 | 1:200 |
| 20 | | UV 379/28 | CD64 (FcγRI) | BUV395 | 10.1 | Mouse IgG1, κ | BioLegend | 740300 | 1:80 |
| 21 | | UV 450/50 | GhostDye™ (Viability) | UV450 | — | — | Tonbo Biosciences | 13-0868-T500 | 1:100 |
| 22 | 355 nm | UV 586/15 | CD56 | BUV563 | NCAM16.2 | Mouse IgG2b, κ | BD Biosciences | 612928 | 1:160 |
| 23 | | UV 670/30 | CD123 (IL-3Rα) | BUV661 | 9F5 | Mouse IgG1, κ | BD Biosciences | 741628 | 1:40 |
| 24 | | UV 740/35 | CD14 | BUV737 | M5E2 | Mouse IgG2a, κ | BD Biosciences | 612763 | 1:50 |
| 25 | | UV 820/60 | CD8α | BUV805 | SK1 | Mouse IgG1, κ | BD Biosciences | 612889 | 1:80 |
| 26 | | R 670/30 | CD66b | APC | QA17A51 | Mouse IgG1, κ | BioLegend | 396906 | 1:200 |
| 27 | 640 nm | R 710/50 | CD11c | AF700 | Bu15 | Mouse IgG1, κ | BioLegend | 337220 | 1:100 |
| 28 | | R 780/60 | CD32 (FcγRII) | APC-Fire750 | FUN-2 | Mouse IgG2b, κ | BioLegend | 303220 | 1:50 |

**Supplementary Table 1. High-dimensional, 30-parameter flow cytometry panel.** Flow cytometer configuration and cytometry reagent details used to interrogate myeloid-derived cells (MdCs) in PBMCs isolated from COVID-19 patients and healthy donors. Monoclonal antibody (mAb) master mixes were prepared in BD Horizon™ Brilliant Stain Buffer and samples stained as described in the methods section. Titrations of all reagents were determined empirically, in-house for each lot independently prior to use. AF: AlexaFluor, APC: Allophycocyanin, BB: Brilliant Blue, BUV: Brilliant Ultraviolet, BV: Brilliant Violet, FITC: Fluorescein isothiocyanate, PE: Phycoerythrin. Fluorophores marked with -P denote prototype reagents and are custom conjugations from BD Biosciences.
