## Supplementary material for "Lifting the curse from high dimensional data: Automated projection pursuit clustering for the variety of biological data modalities": Supplementary_Data_Pairwise_Projections_PBMCs.pdf

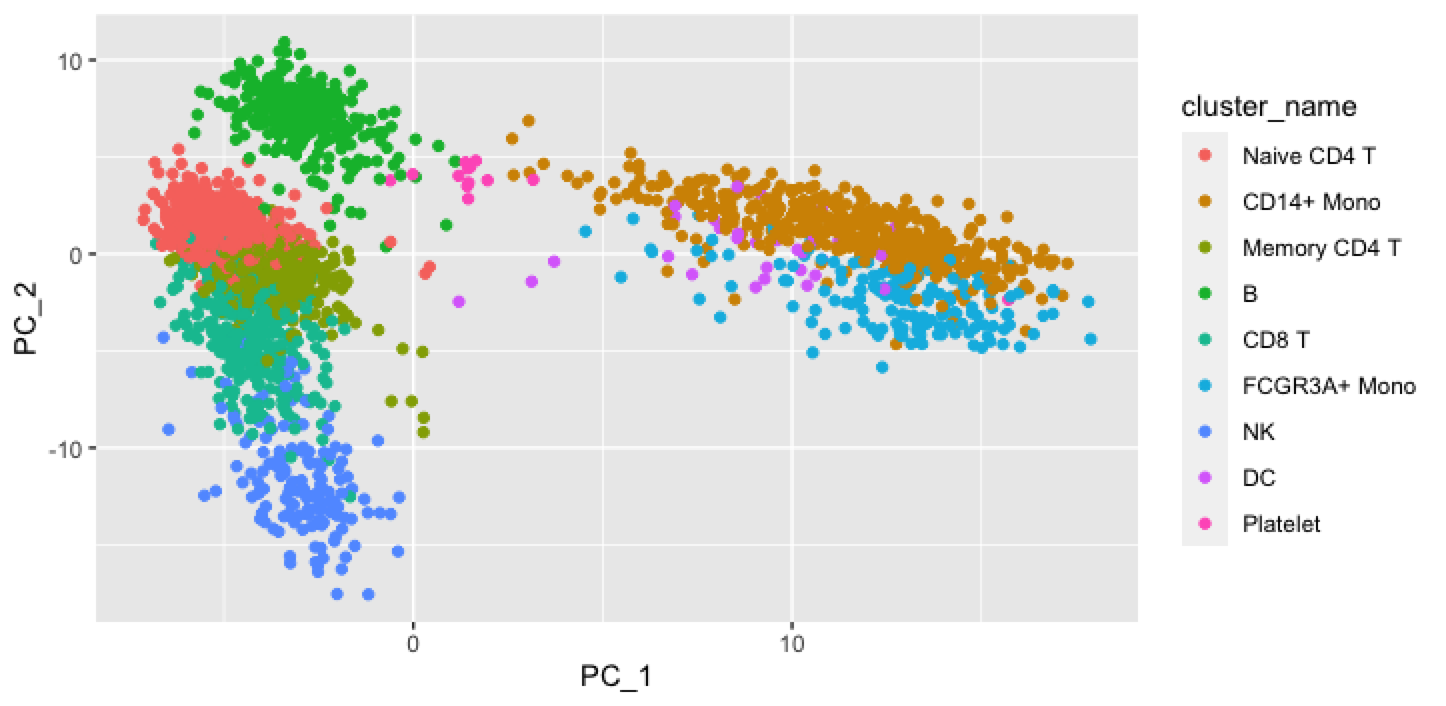

10
0
-10
PC_2
0
10
PC_1
cluster_name
Naive CD4 T
CD14+ Mono
Memory CD4 T
B
CD8 T
FCGR3A+ Mono
NK
DC
Platelet

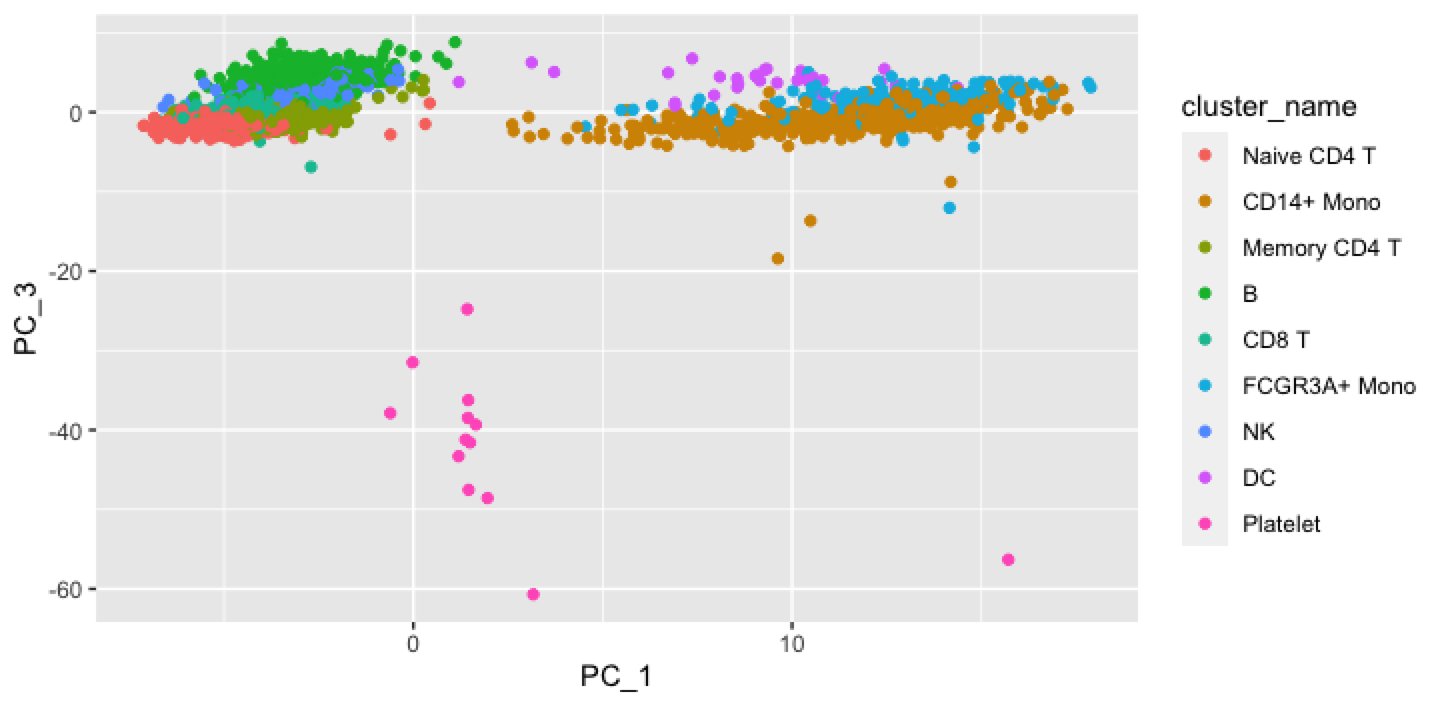
PC_3
PC_1
0
-20
-40
-60
0
10
cluster_name
Naive CD4 T
CD14+ Mono
Memory CD4 T
B
CD8 T
FCGR3A+ Mono
NK
DC
Platelet

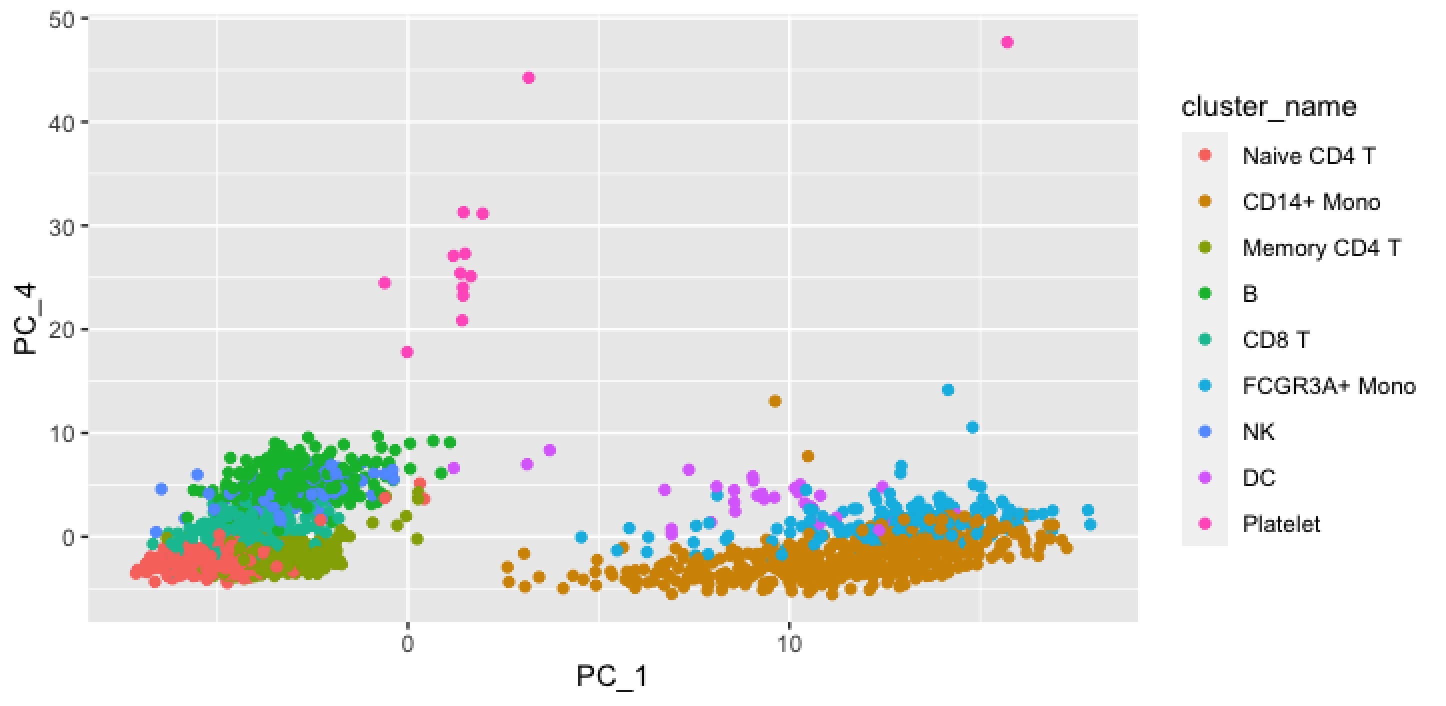

50
40
30
20
10
0
PC_4
0
10
PC_1
cluster_name
Naive CD4 T
CD14+ Mono
Memory CD4 T
B
CD8 T
FCGR3A+ Mono
NK
DC
Platelet

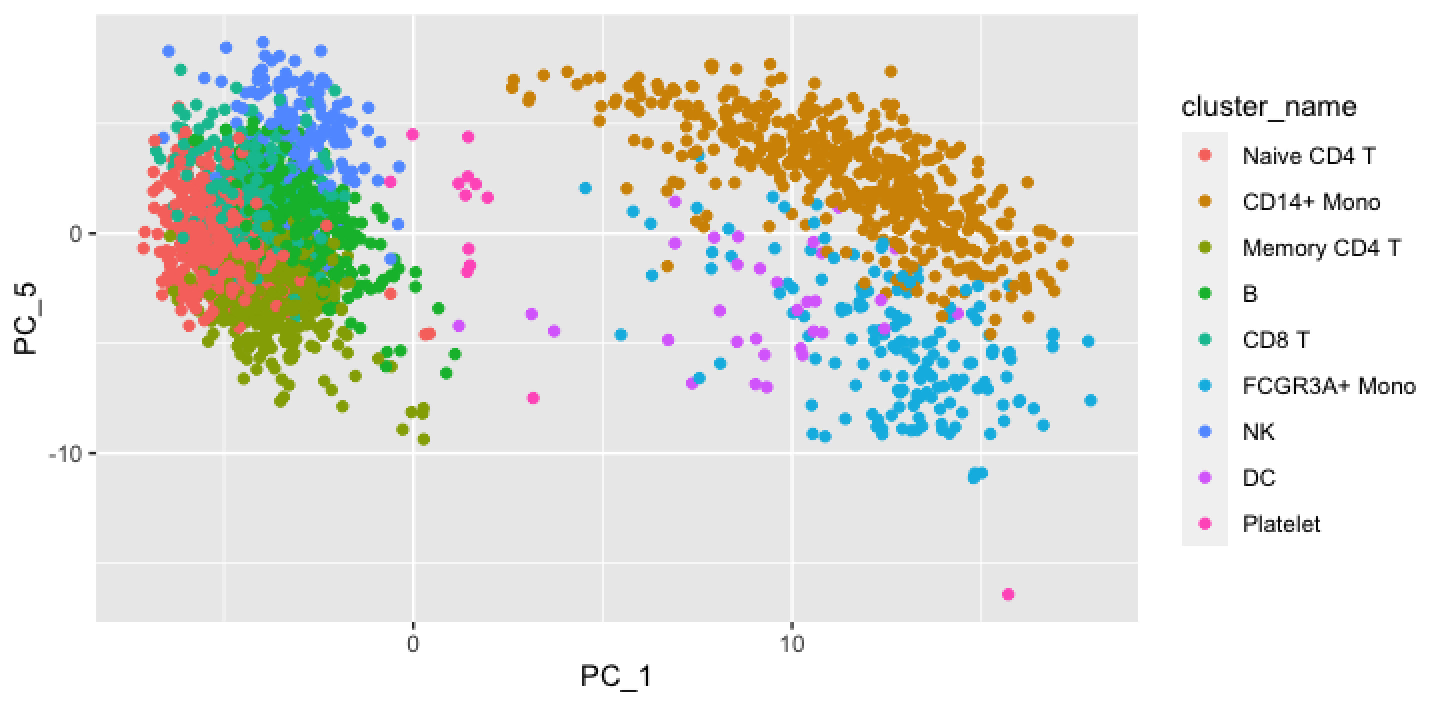
cluster_name
Naive CD4 T
CD14+ Mono
Memory CD4 T
B
CD8 T
FCGR3A+ Mono
NK
DC
Platelet
PC_5
PC_1
0
-10
0
10

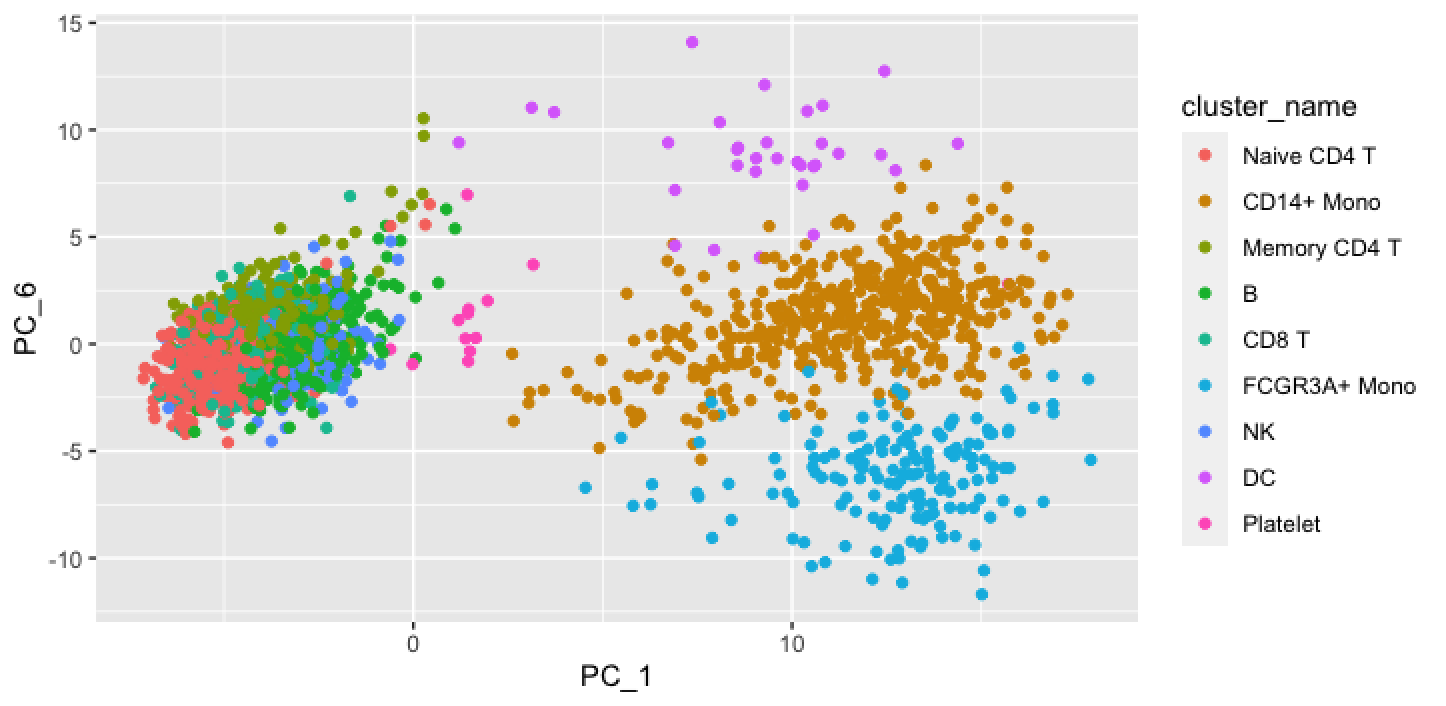
15
10
5
0
-5
-10
PC_6
0
10
PC_1
cluster_name
Naive CD4 T
CD14+ Mono
Memory CD4 T
B
CD8 T
FCGR3A+ Mono
NK
DC
Platelet

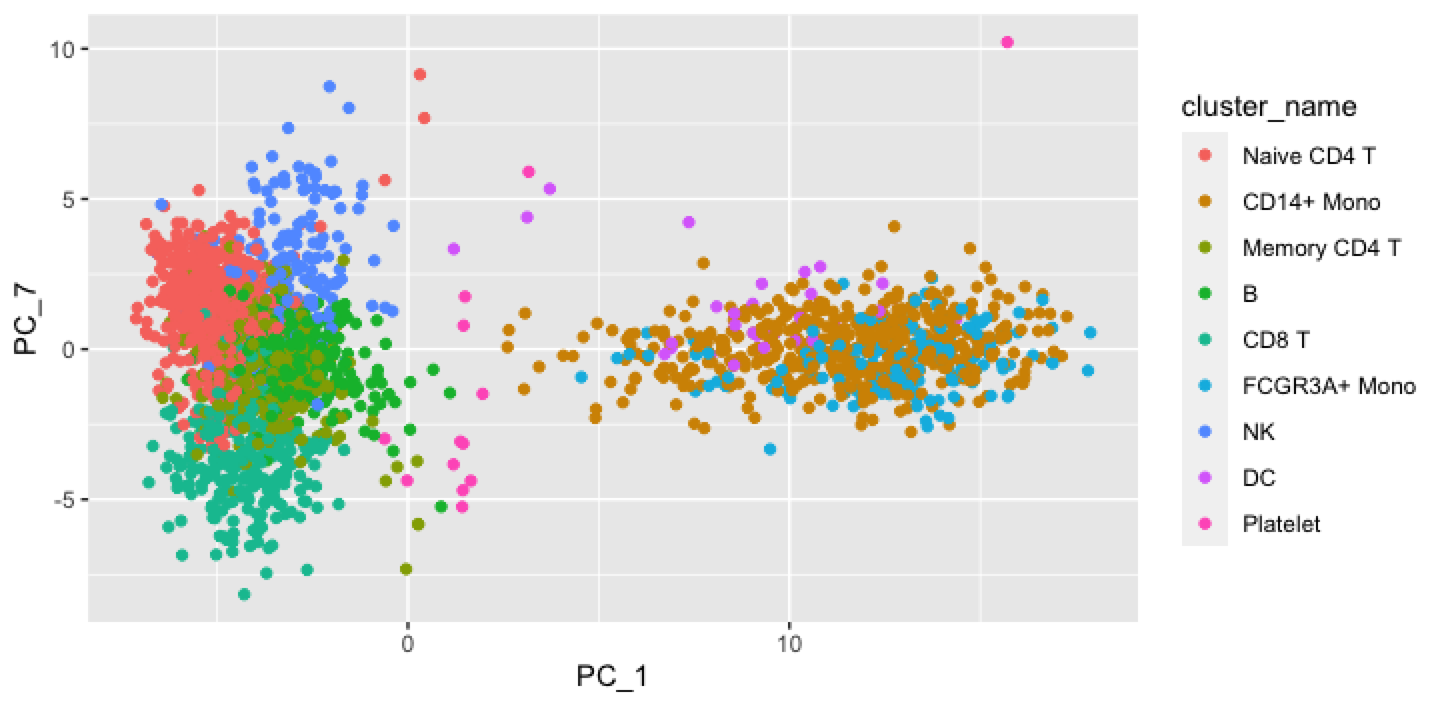
10
5
0
-5
PC_7
0
10
PC_1
cluster_name
Naive CD4 T
CD14+ Mono
Memory CD4 T
B
CD8 T
FCGR3A+ Mono
NK
DC
Platelet

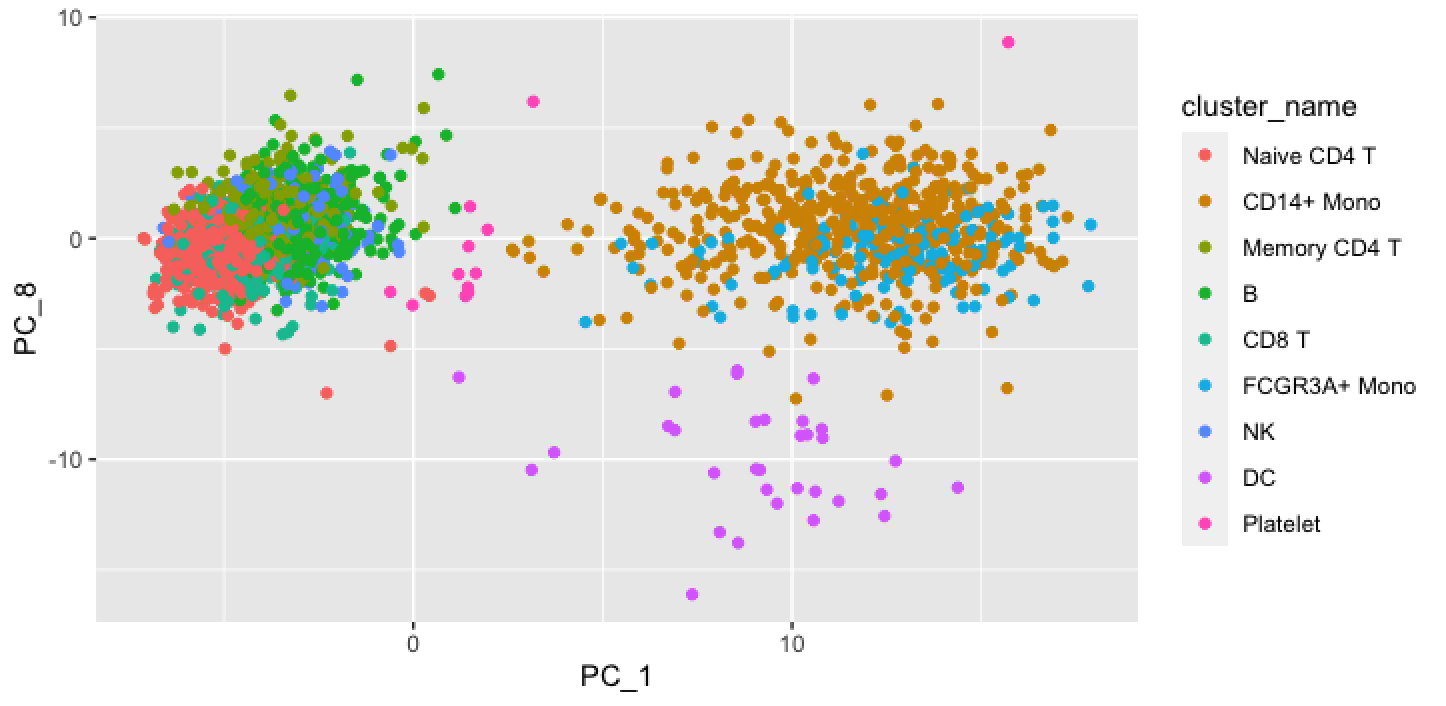

10
0
-10
PC_8
0
10
PC_1
cluster_name
Naive CD4 T
CD14+ Mono
Memory CD4 T
B
CD8 T
FCGR3A+ Mono
NK
DC
Platelet

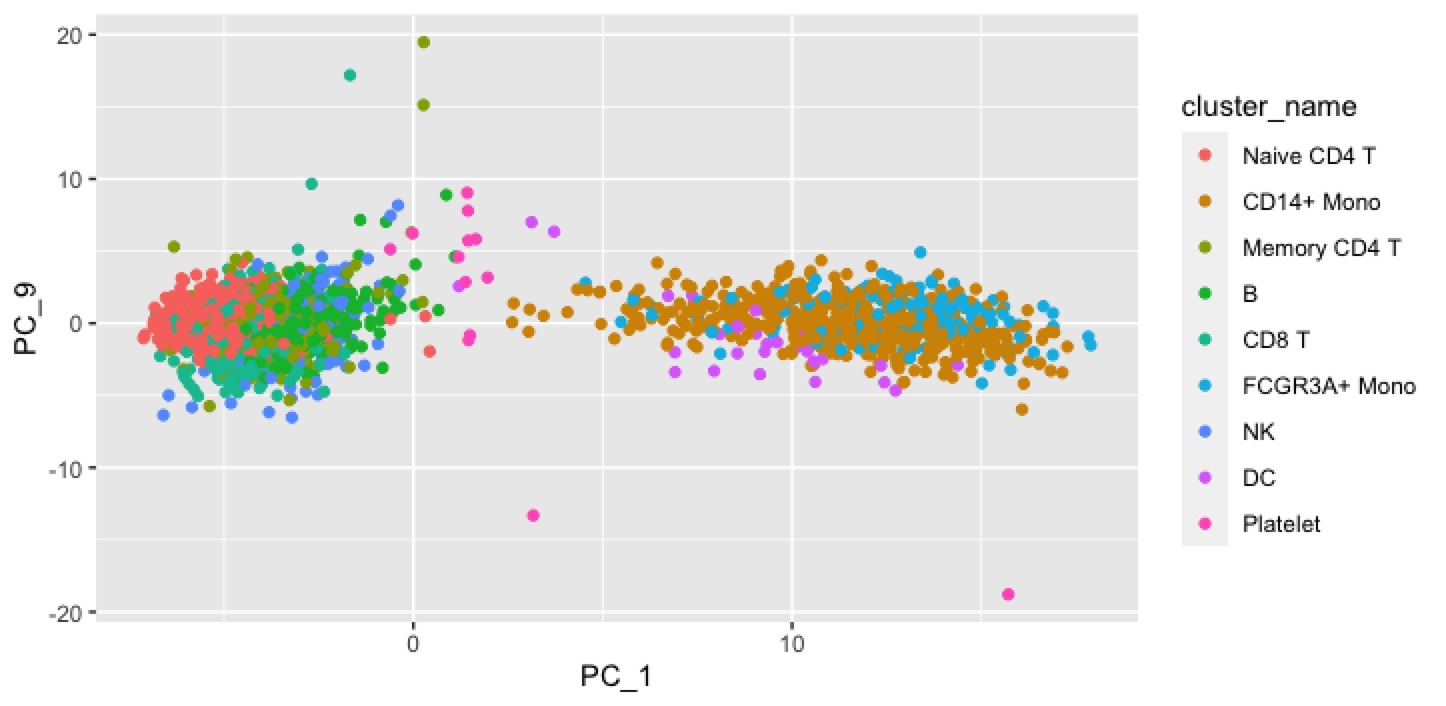

20
10
0
-10
-20
PC_9
0
10
PC_1
cluster_name
Naive CD4 T
CD14+ Mono
Memory CD4 T
B
CD8 T
FCGR3A+ Mono
NK
DC
Platelet

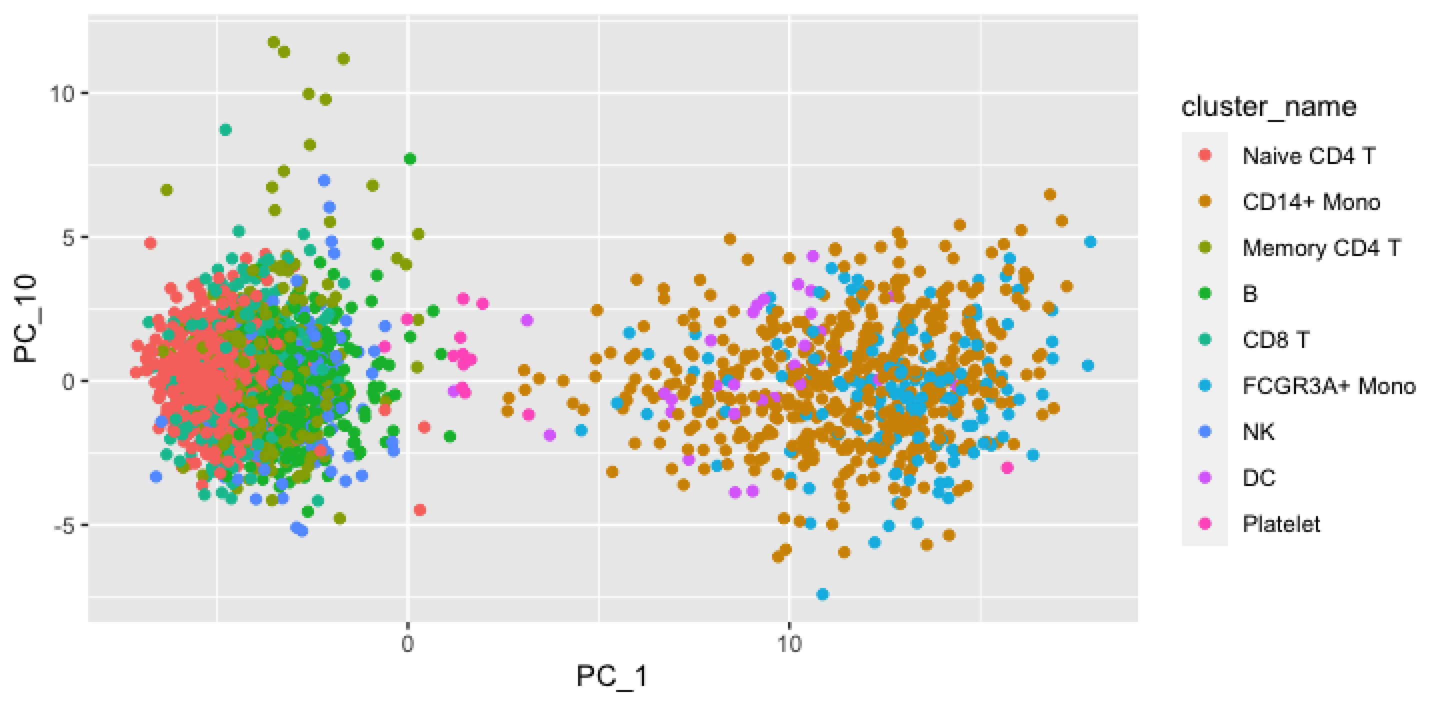

10
5
0
-5
PC_10
0
10
PC_1
cluster_name
Naive CD4 T
CD14+ Mono
Memory CD4 T
B
CD8 T
FCGR3A+ Mono
NK
DC
Platelet

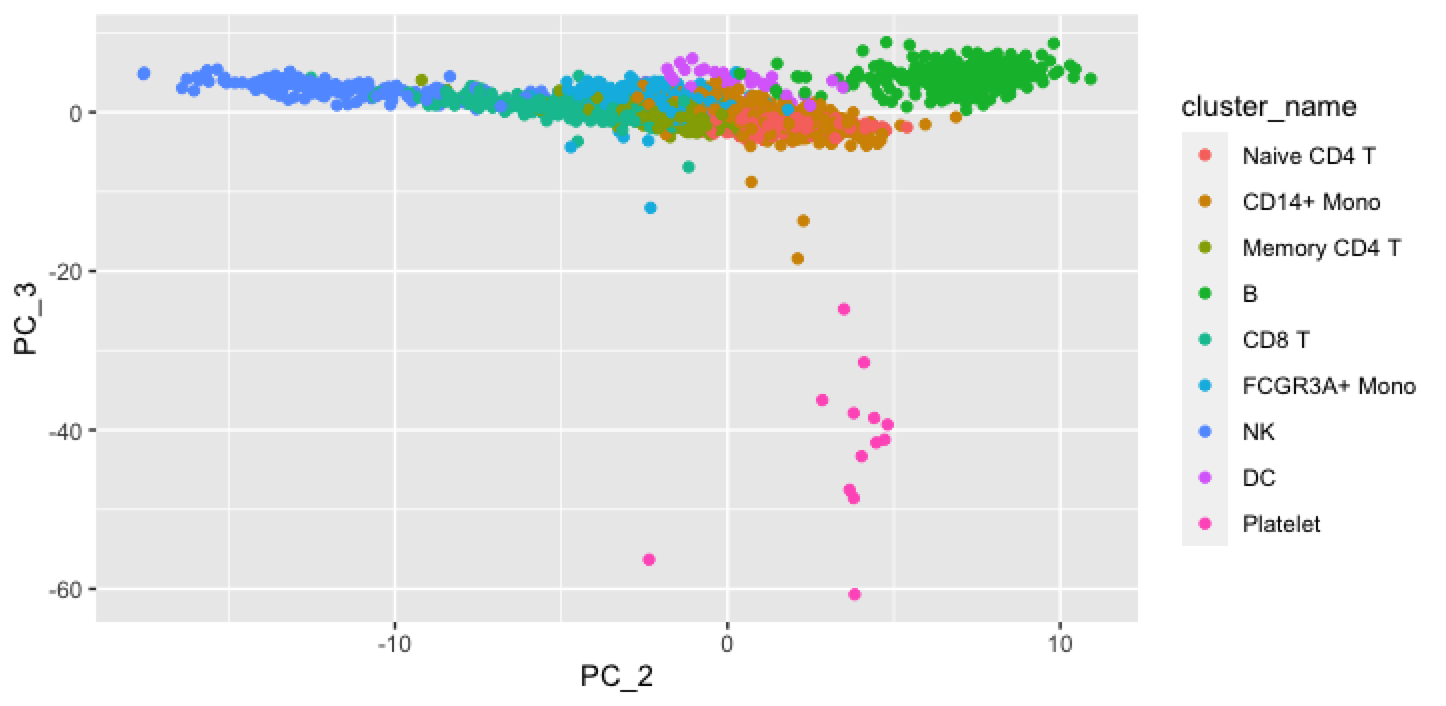

PC_3
0
-20
-40
-60
-10
0
10
PC_2
cluster_name
Naive CD4 T
CD14+ Mono
Memory CD4 T
B
CD8 T
FCGR3A+ Mono
NK
DC
Platelet

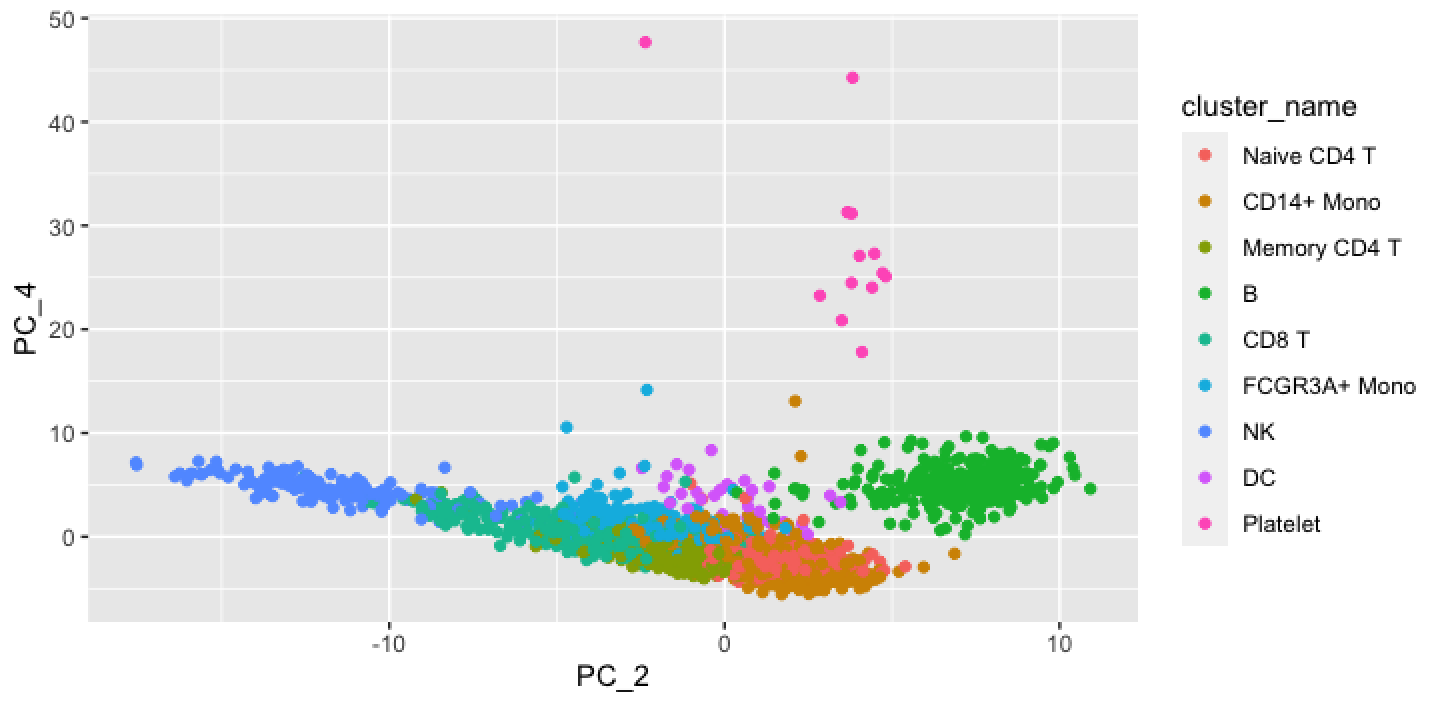

PC_4
PC_2
50
40
30
20
10
0
-10
0
10
cluster_name
Naive CD4 T
CD14+ Mono
Memory CD4 T
B
CD8 T
FCGR3A+ Mono
NK
DC
Platelet

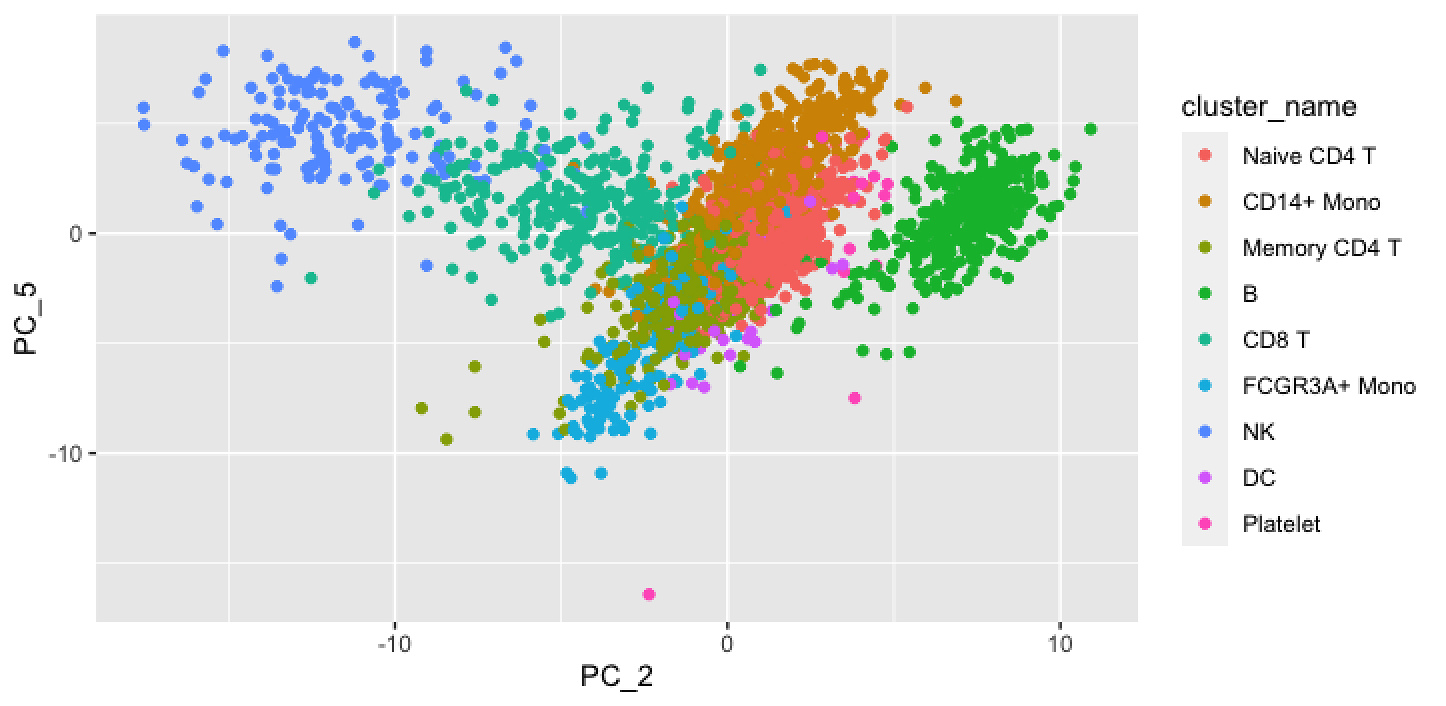

cluster_name
Naive CD4 T
CD14+ Mono
Memory CD4 T
B
CD8 T
FCGR3A+ Mono
NK
DC
Platelet
PC_5
PC_2
0
-10
10

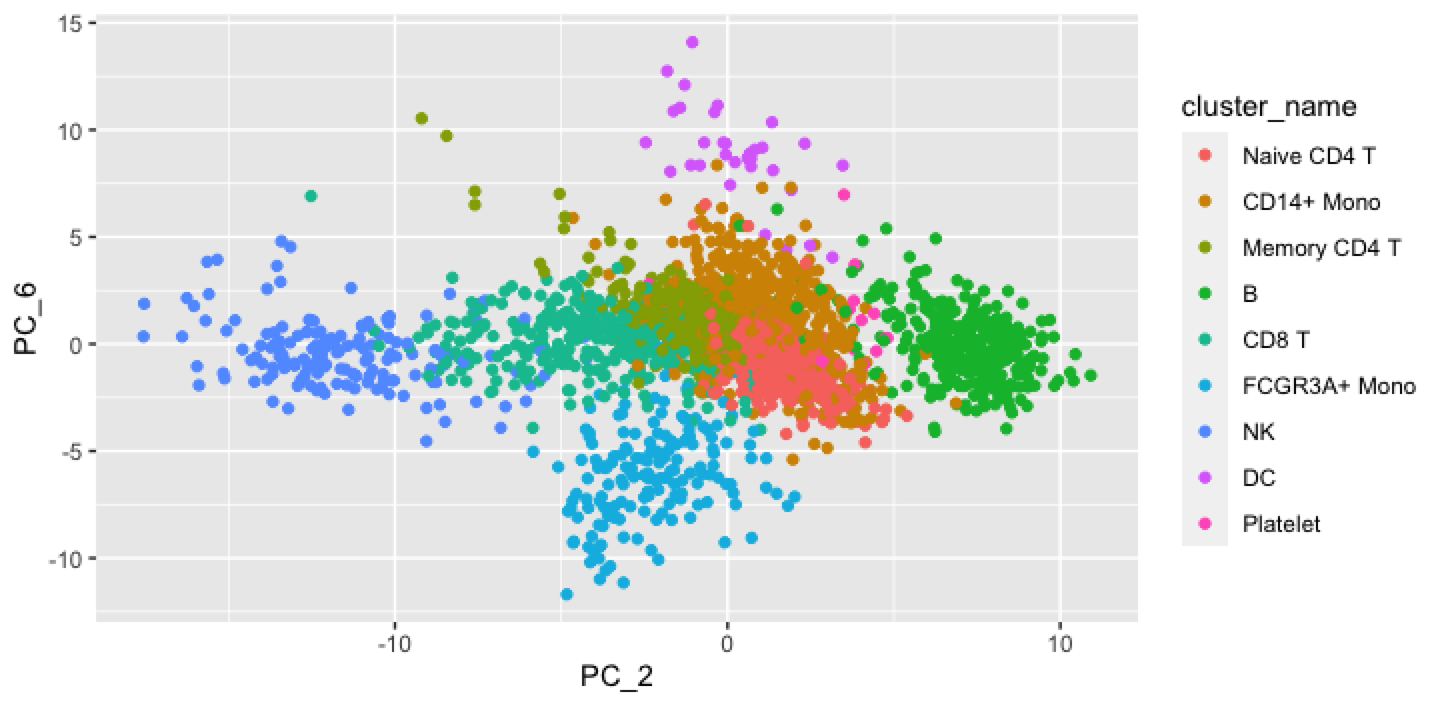

15
10
5
0
-5
-10
PC_6
-10
0
10
PC_2
cluster_name
Naive CD4 T
CD14+ Mono
Memory CD4 T
B
CD8 T
FCGR3A+ Mono
NK
DC
Platelet

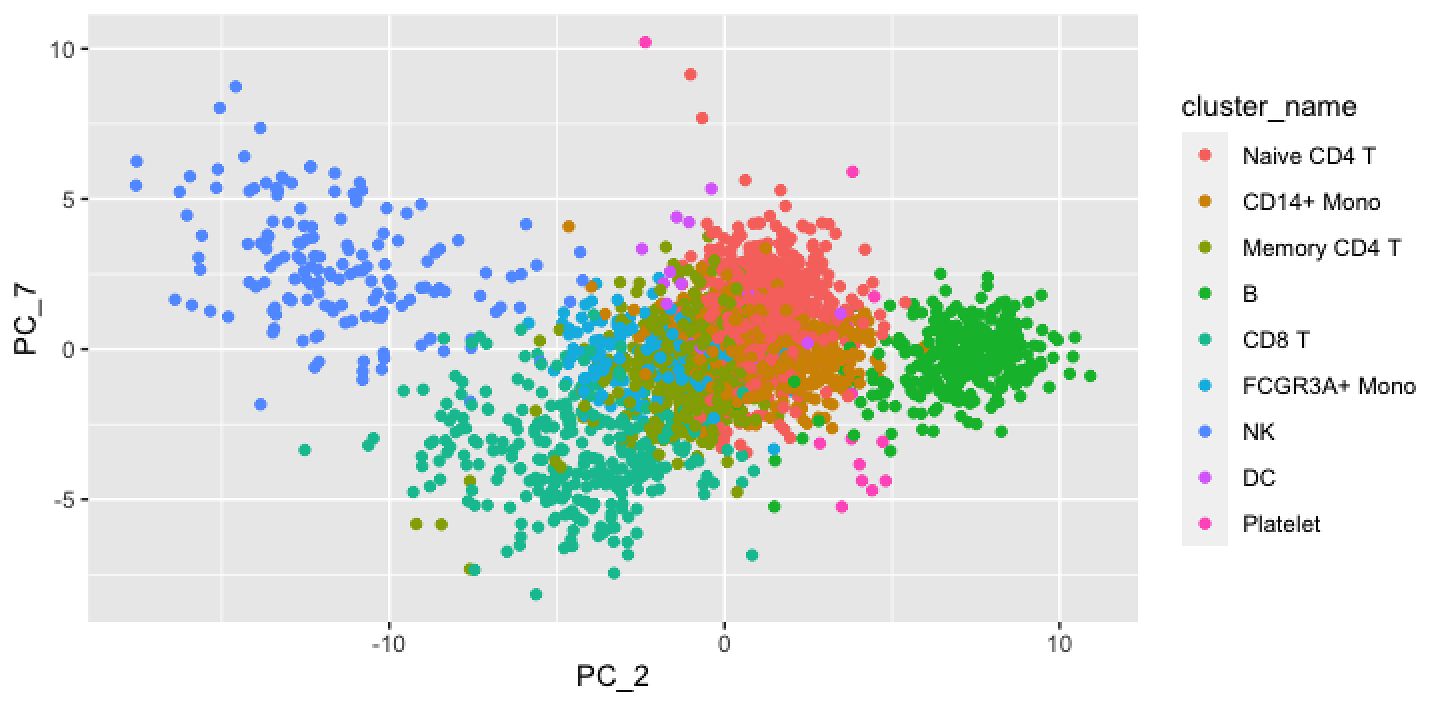

10
5
0
-5
-10
0
10
PC_7
PC_2
cluster_name
Naive CD4 T
CD14+ Mono
Memory CD4 T
B
CD8 T
FCGR3A+ Mono
NK
DC
Platelet

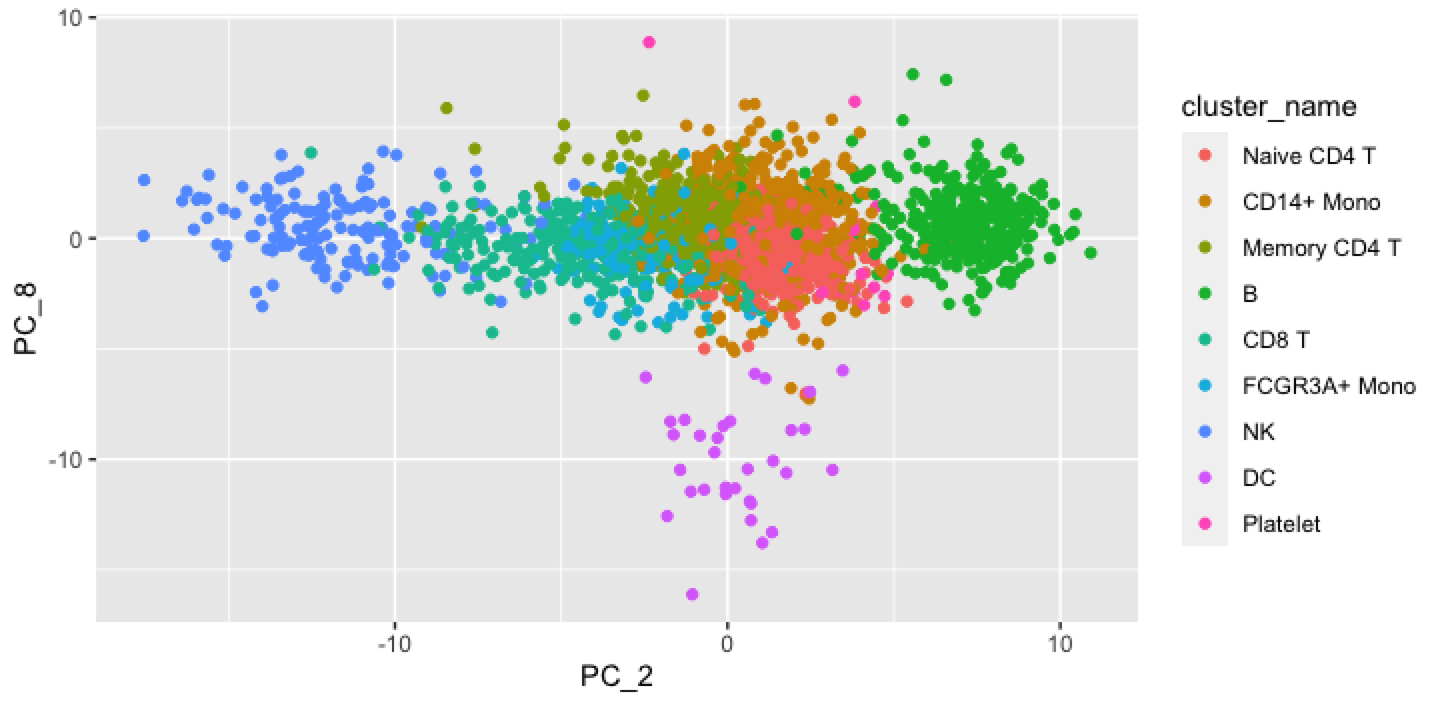

10
0
-10
-10
0
10
PC_8
PC_2
cluster_name
Naive CD4 T
CD14+ Mono
Memory CD4 T
B
CD8 T
FCGR3A+ Mono
NK
DC
Platelet

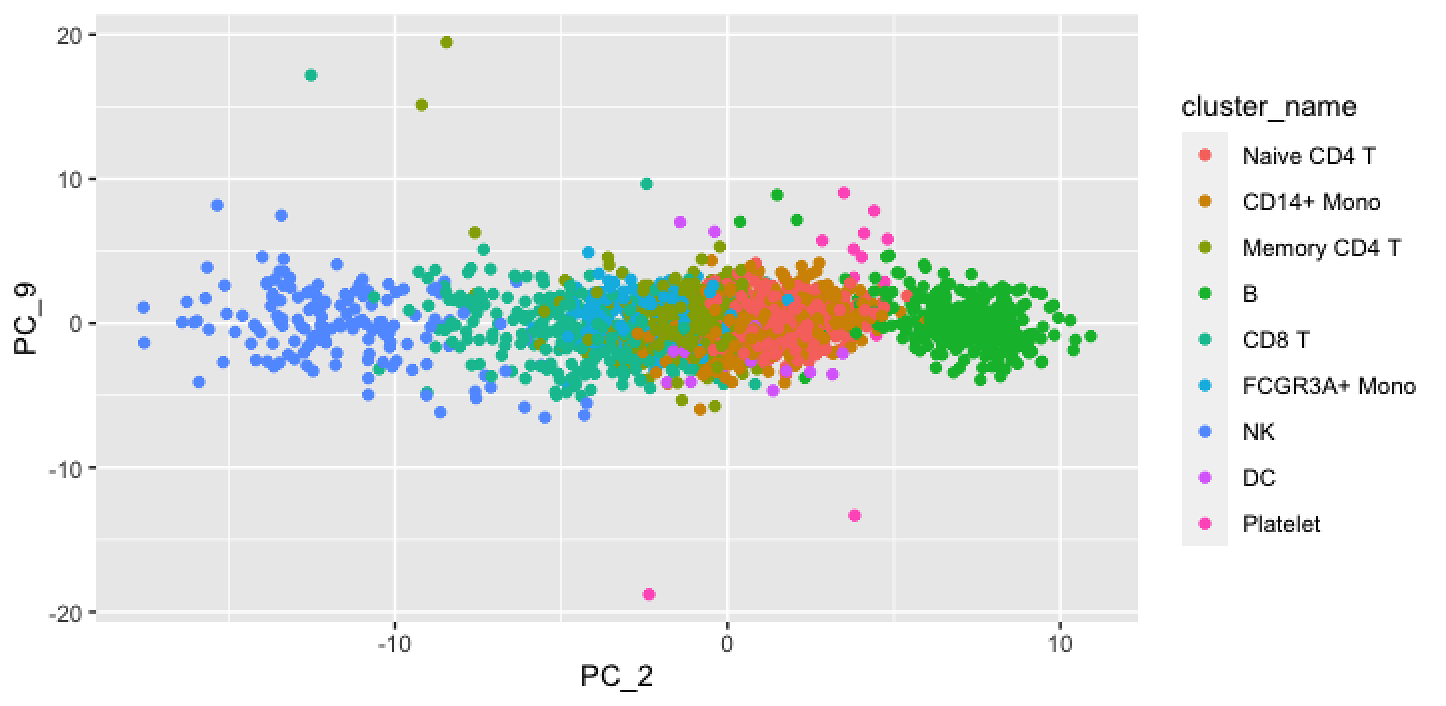

20
10
0
-10
-20
PC_9
-10
0
10
PC_2
cluster_name
Naive CD4 T
CD14+ Mono
Memory CD4 T
B
CD8 T
FCGR3A+ Mono
NK
DC
Platelet

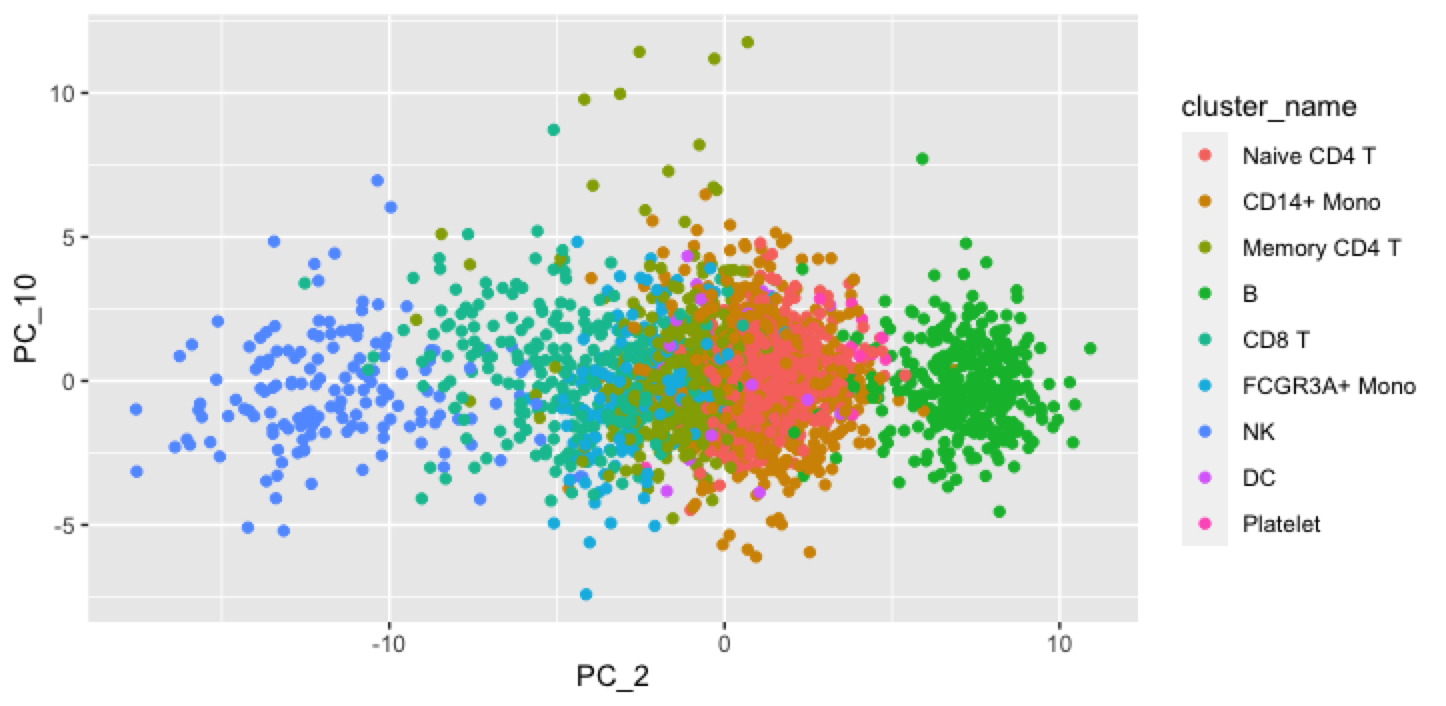

cluster_name
Naive CD4 T
CD14+ Mono
Memory CD4 T
B
CD8 T
FCGR3A+ Mono
NK
DC
Platelet
PC_10
PC_2
10
5
0
-5
-10
0
10

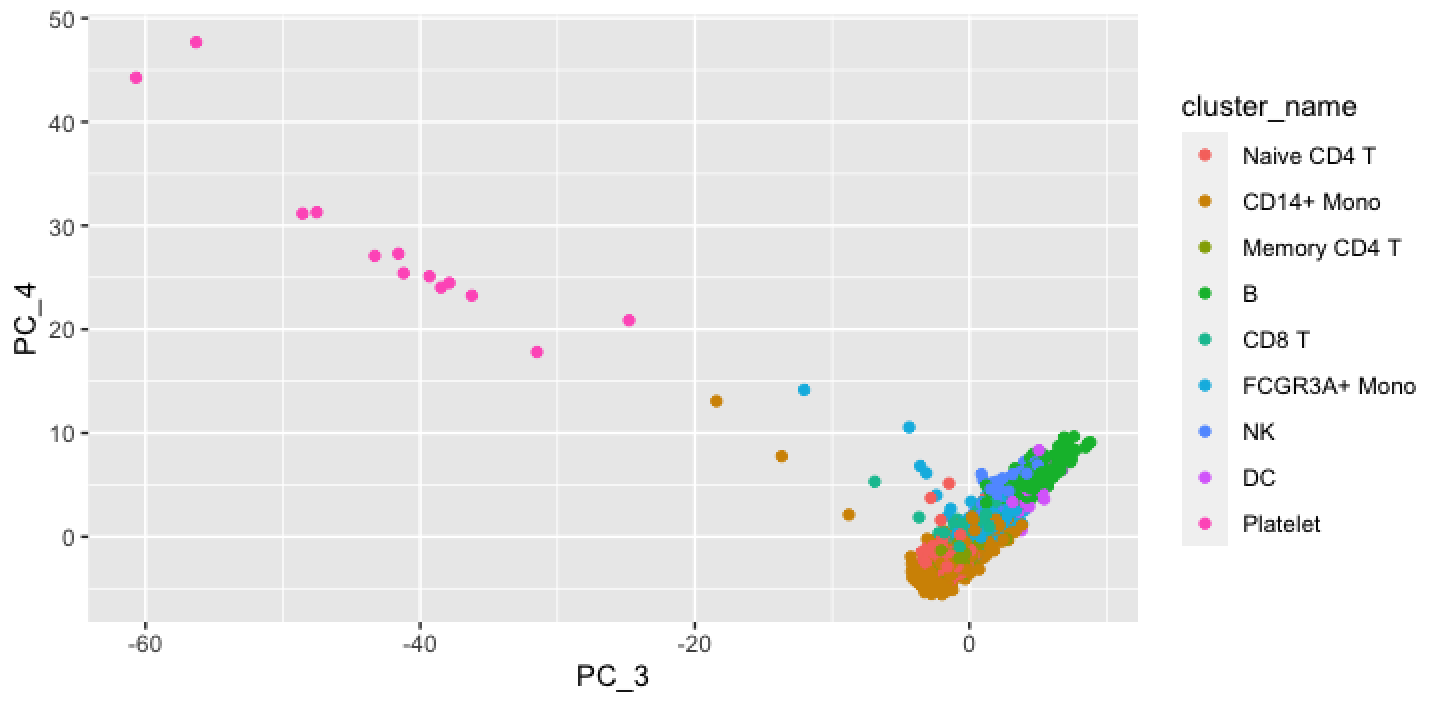
50
40
30
20
10
0
PC_4
-60
-40
-20
0
PC_3
cluster_name
Naive CD4 T
CD14+ Mono
Memory CD4 T
B
CD8 T
FCGR3A+ Mono
NK
DC
Platelet

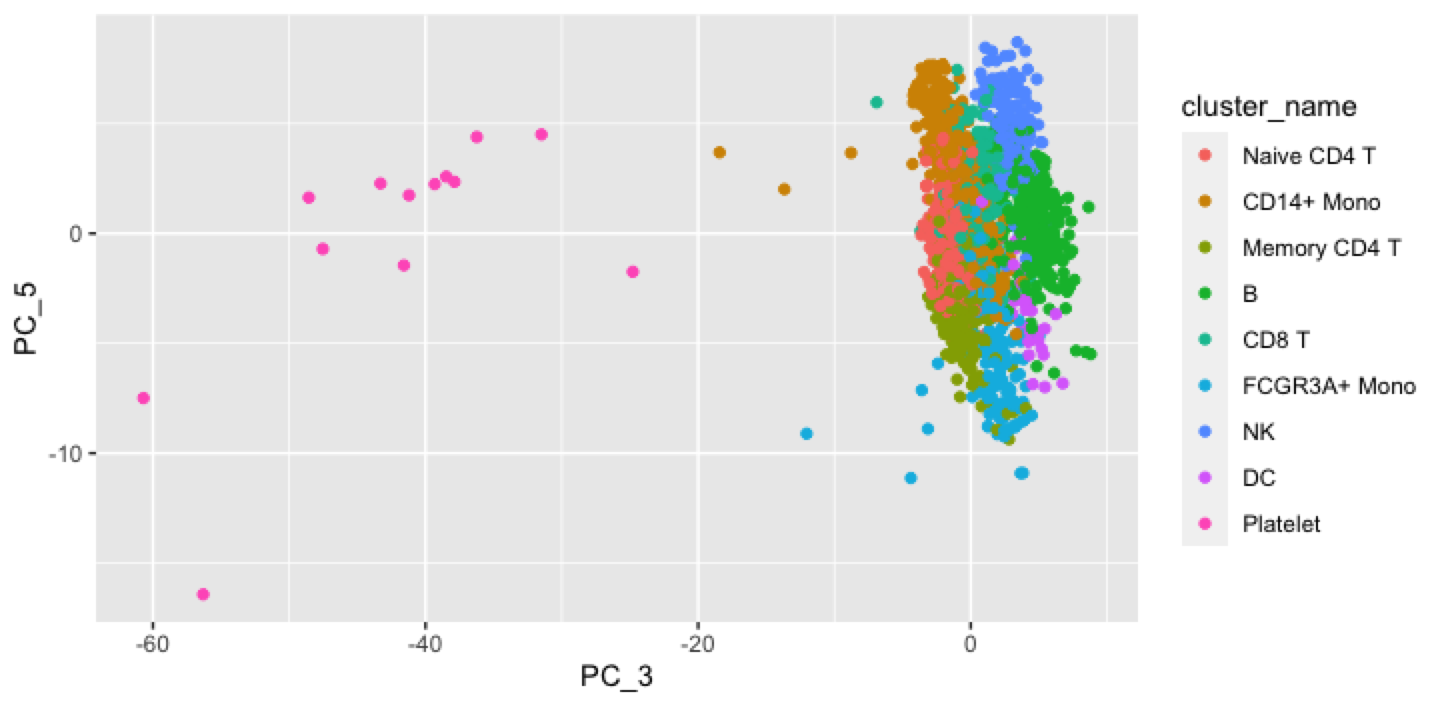

PC_5
0
-10
-60
-40
-20
0
PC_3
cluster_name
Naive CD4 T
CD14+ Mono
Memory CD4 T
B
CD8 T
FCGR3A+ Mono
NK
DC
Platelet

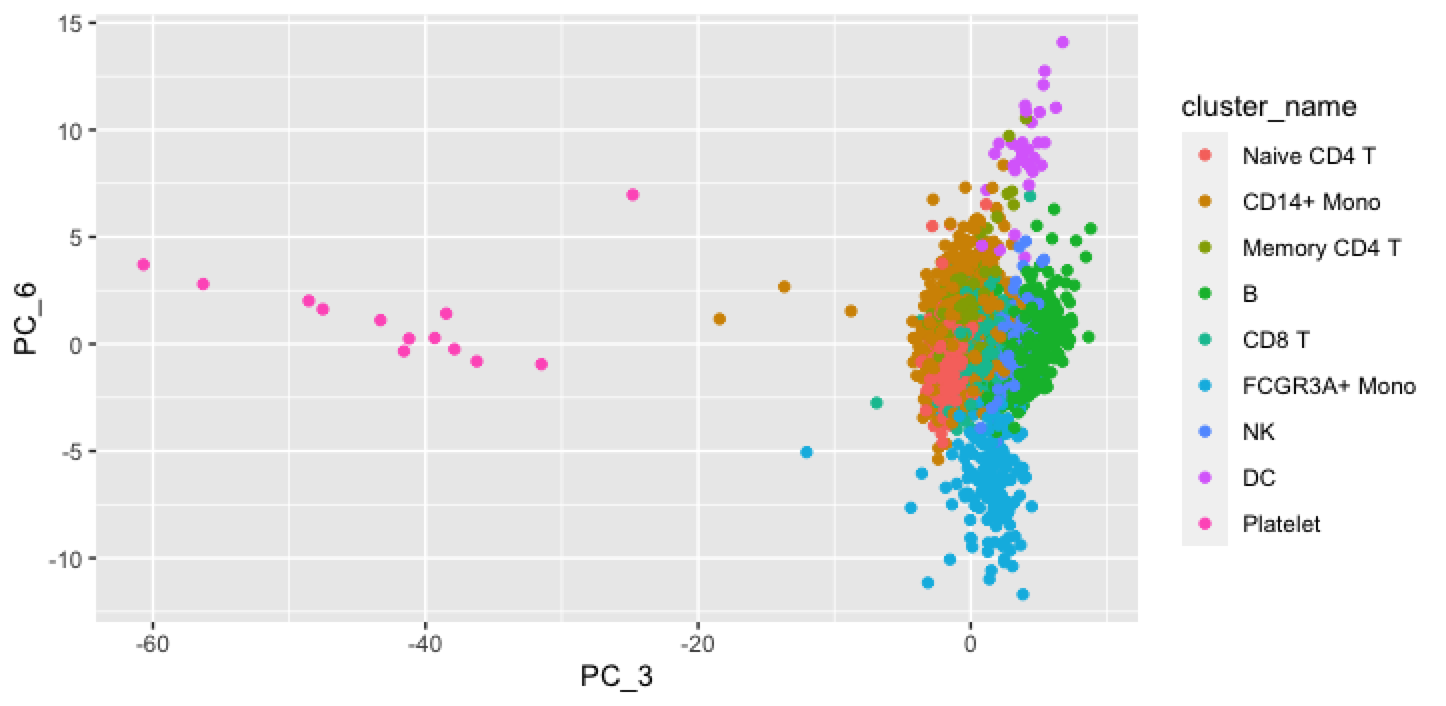

15
10
5
0
-5
-10
PC_6
-60
-40
-20
0
PC_3
cluster_name
Naive CD4 T
CD14+ Mono
Memory CD4 T
B
CD8 T
FCGR3A+ Mono
NK
DC
Platelet

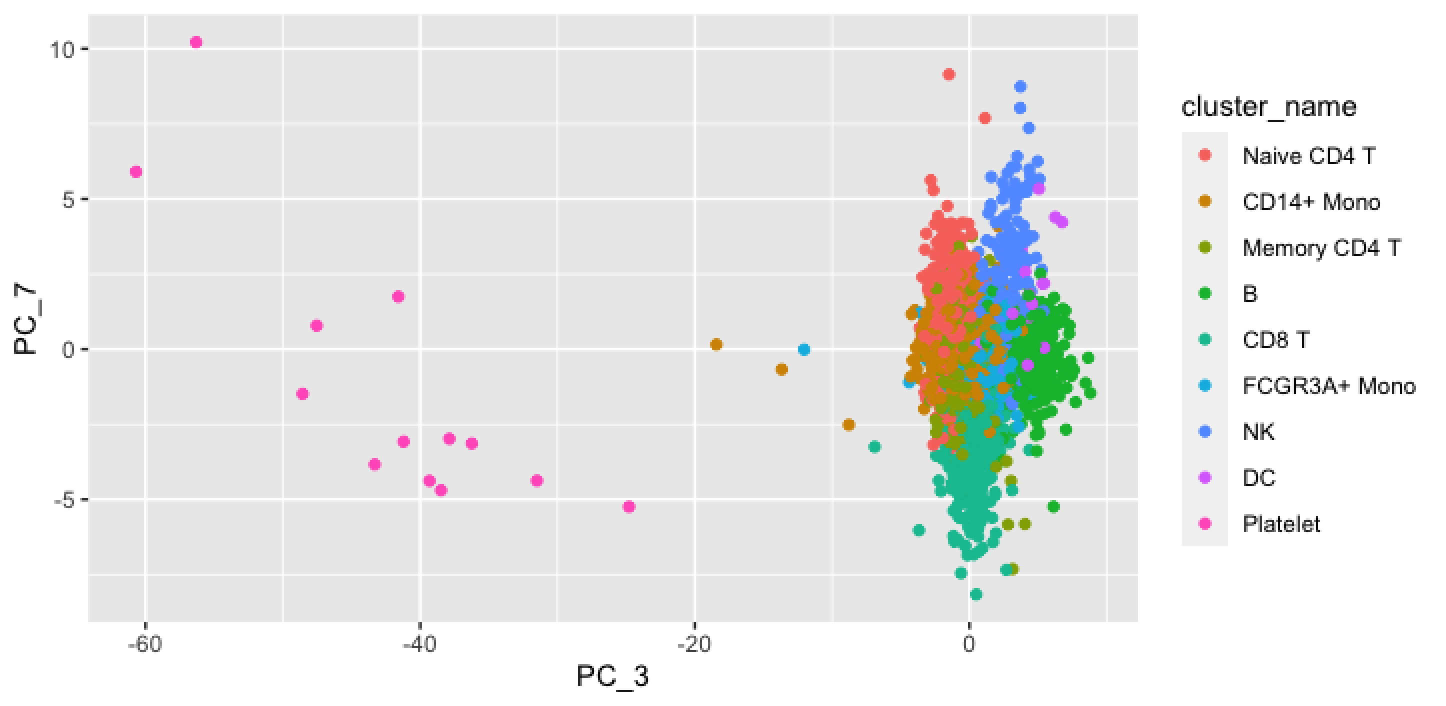
10
5
0
-5
PC_7
-60
-40
-20
0
PC_3
cluster_name
Naive CD4 T
CD14+ Mono
Memory CD4 T
B
CD8 T
FCGR3A+ Mono
NK
DC
Platelet

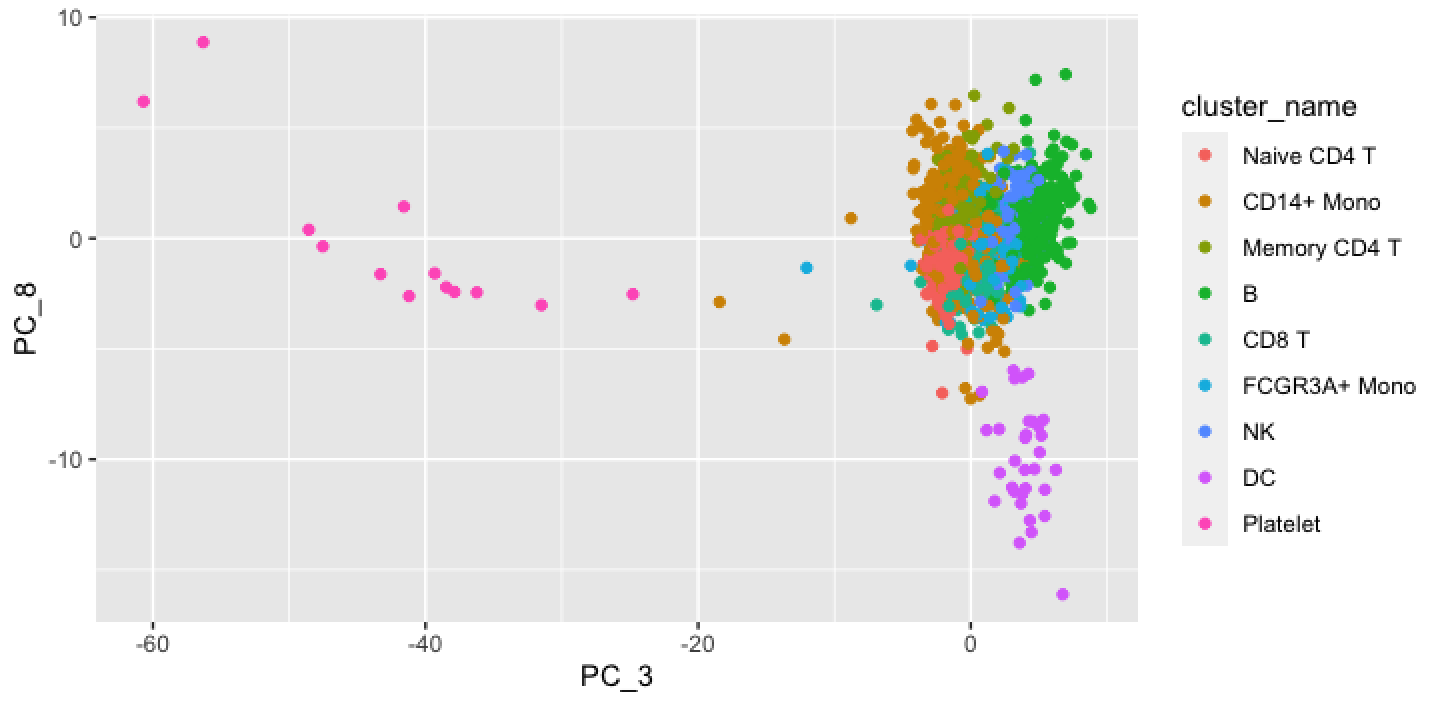
10
0
-10
PC_8
-60
-40
-20
0
PC_3
cluster_name
Naive CD4 T
CD14+ Mono
Memory CD4 T
B
CD8 T
FCGR3A+ Mono
NK
DC
Platelet

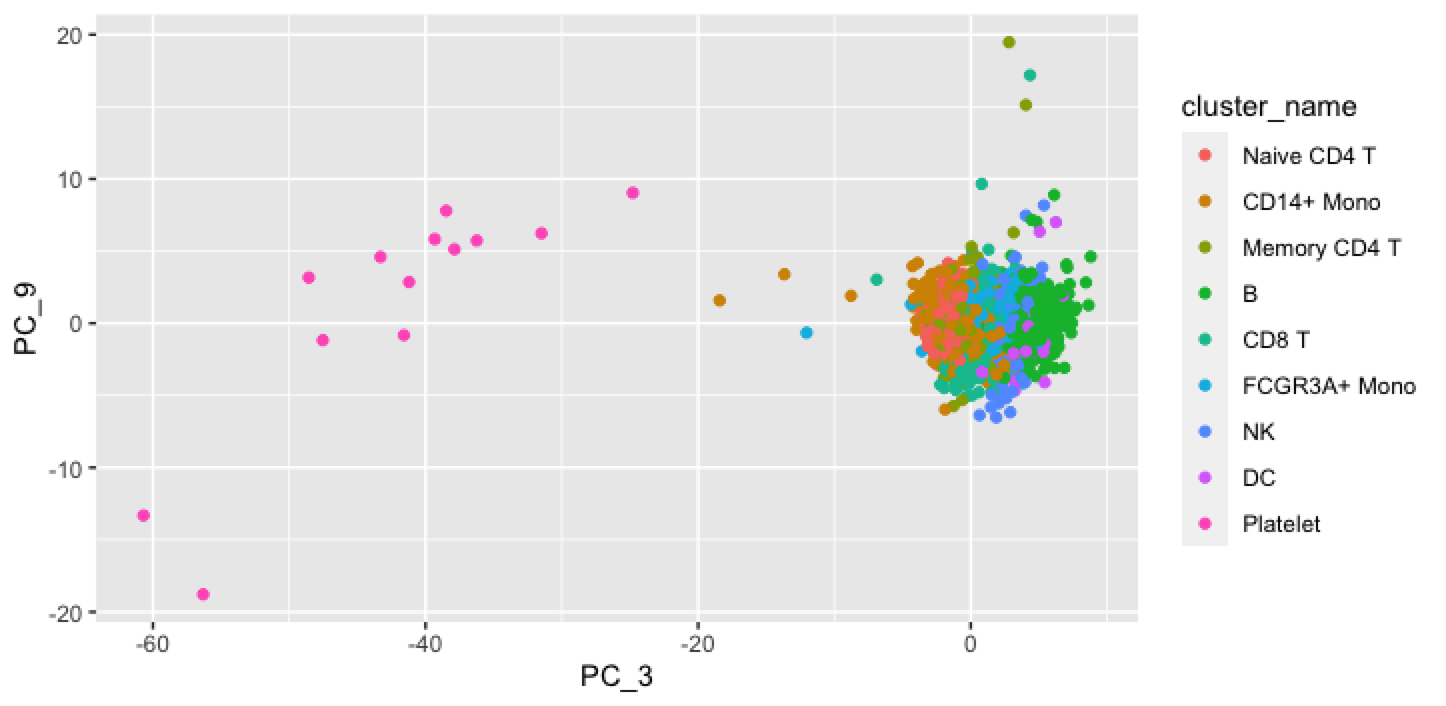

20
10
0
-10
-20
PC_9
-60
-40
-20
0
PC_3
cluster_name
Naive CD4 T
CD14+ Mono
Memory CD4 T
B
CD8 T
FCGR3A+ Mono
NK
DC
Platelet

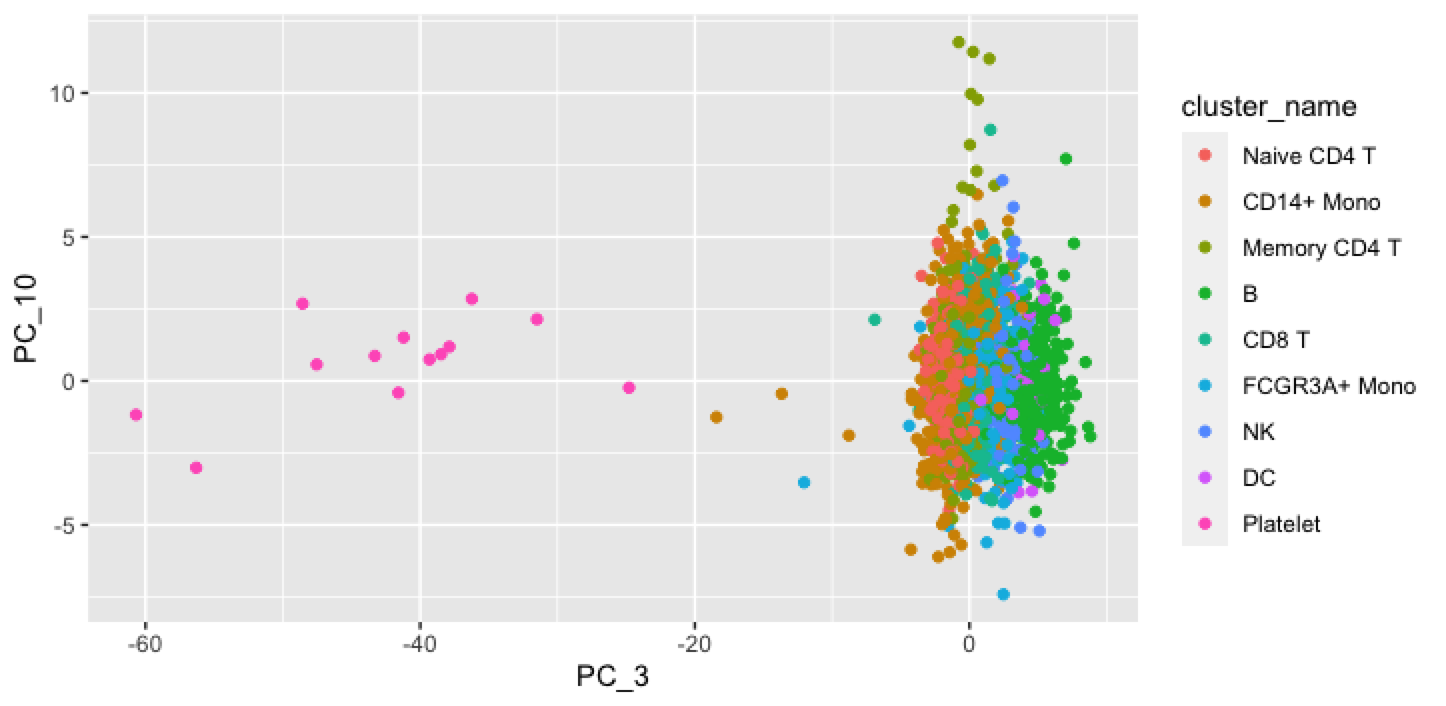

10
5
0
-5
PC_10
-60
-40
-20
0
PC_3
cluster_name
Naive CD4 T
CD14+ Mono
Memory CD4 T
B
CD8 T
FCGR3A+ Mono
NK
DC
Platelet

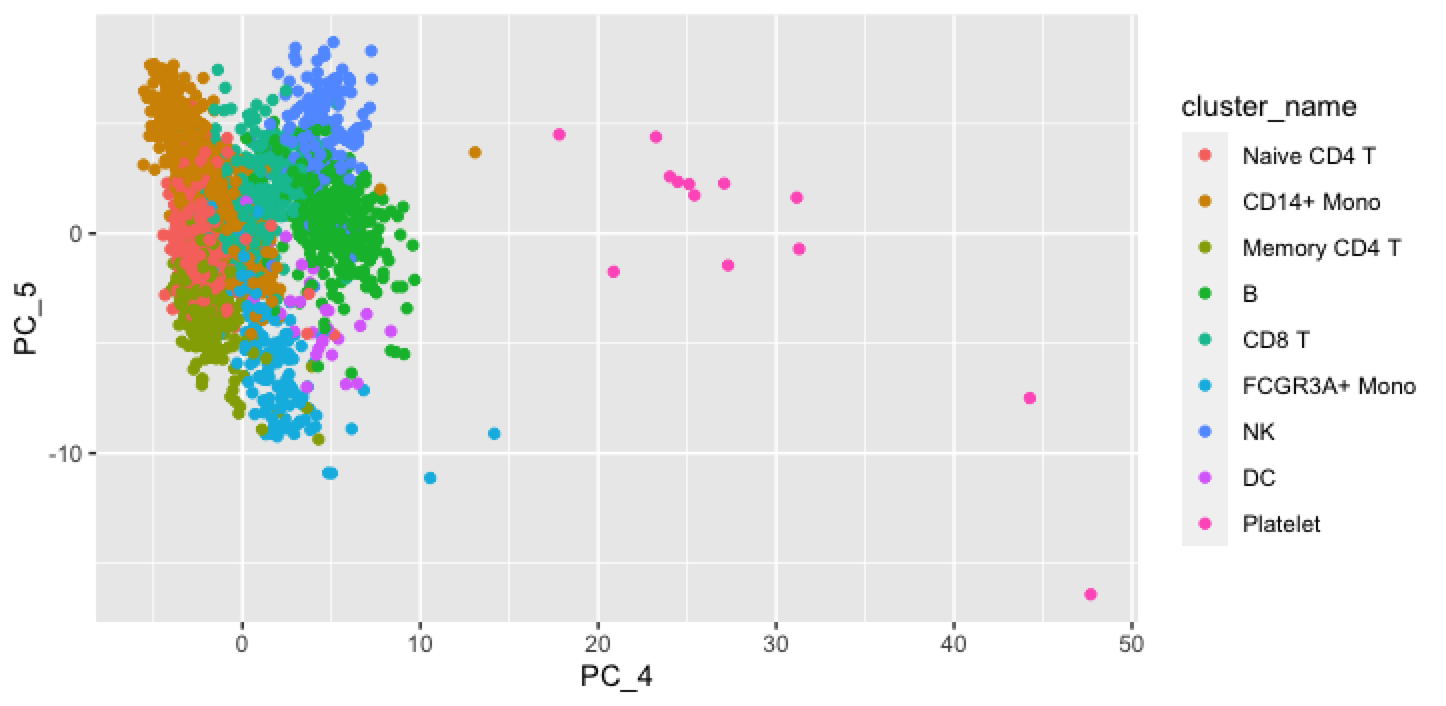

cluster_name
Naive CD4 T
CD14+ Mono
Memory CD4 T
B
CD8 T
FCGR3A+ Mono
NK
DC
Platelet
PC_5
PC_4
0
-10
0
10
20
30
40
50

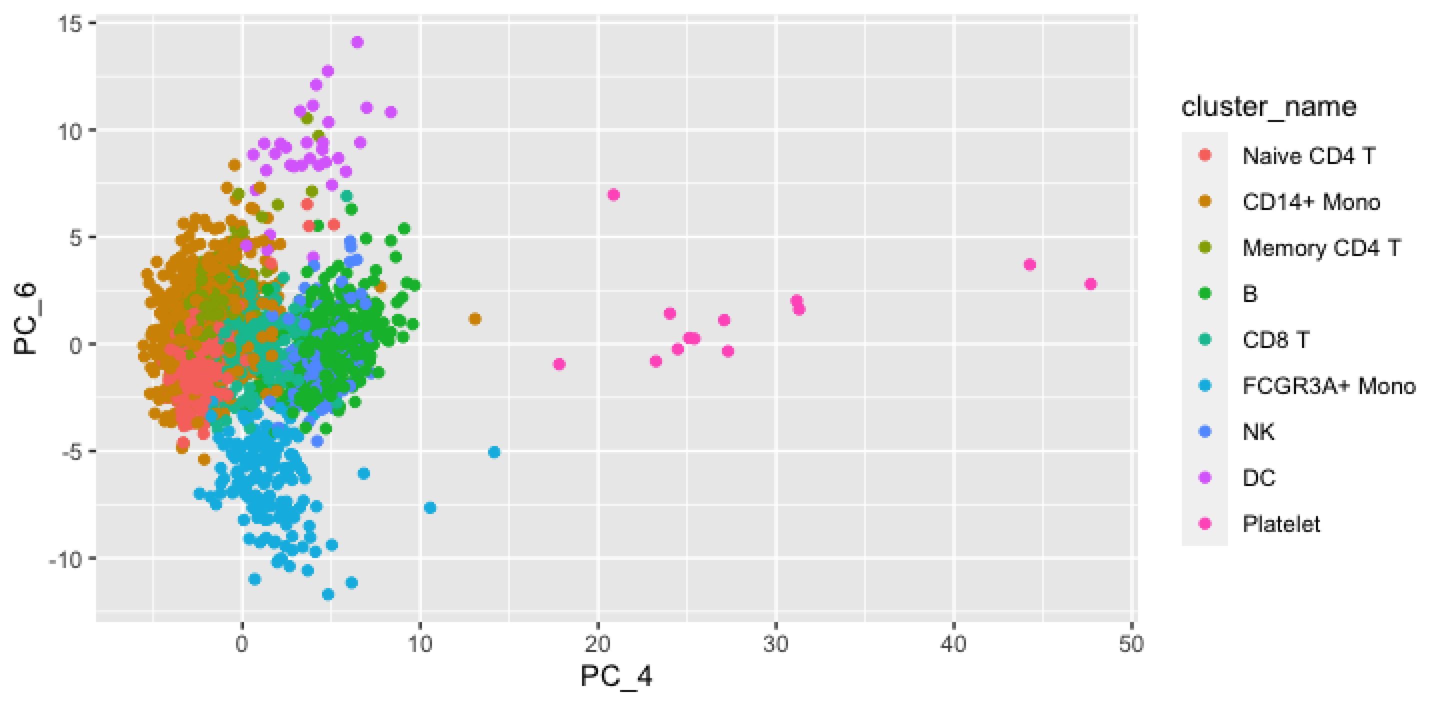

15
10
5
0
-5
-10
PC_6
0
10
20
30
40
50
PC_4
cluster_name
Naive CD4 T
CD14+ Mono
Memory CD4 T
B
CD8 T
FCGR3A+ Mono
NK
DC
Platelet

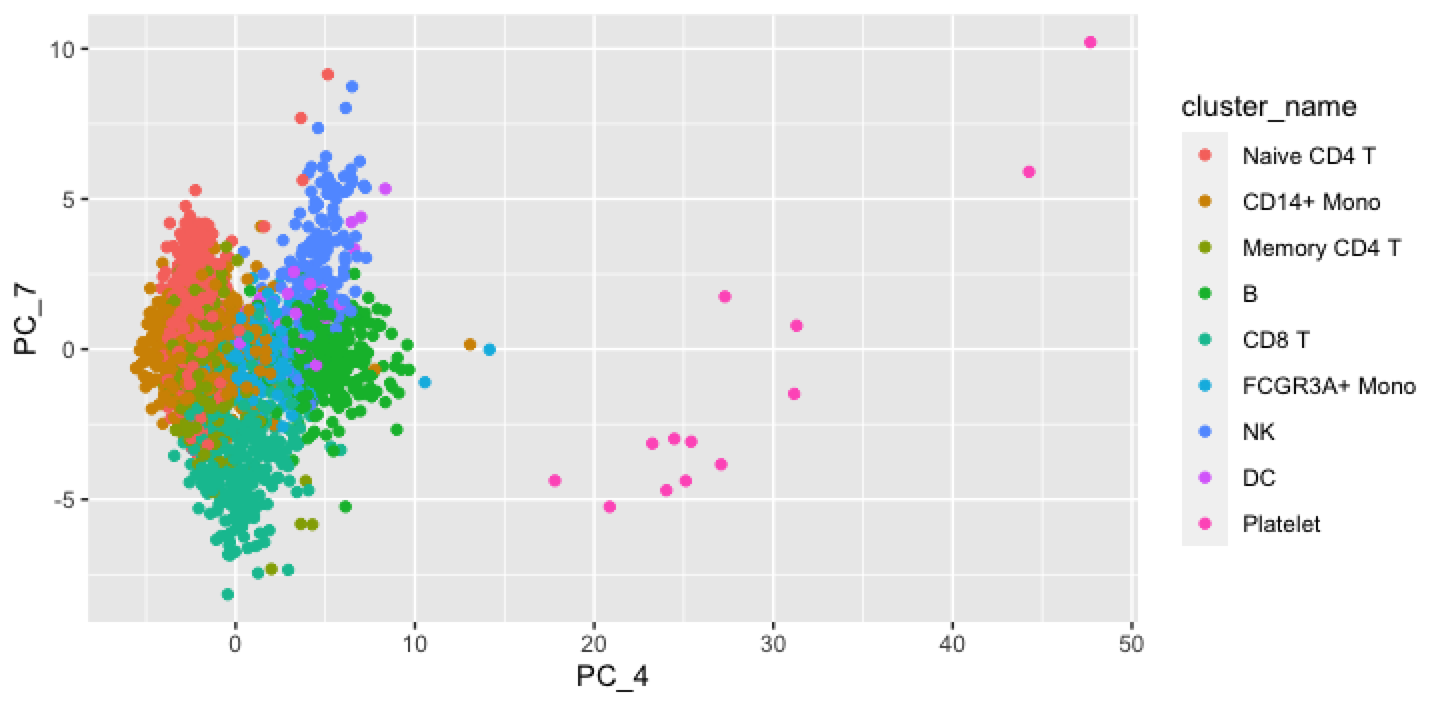

10
5
0
-5
PC_7
0
10
20
30
40
50
PC_4
cluster_name
Naive CD4 T
CD14+ Mono
Memory CD4 T
B
CD8 T
FCGR3A+ Mono
NK
DC
Platelet

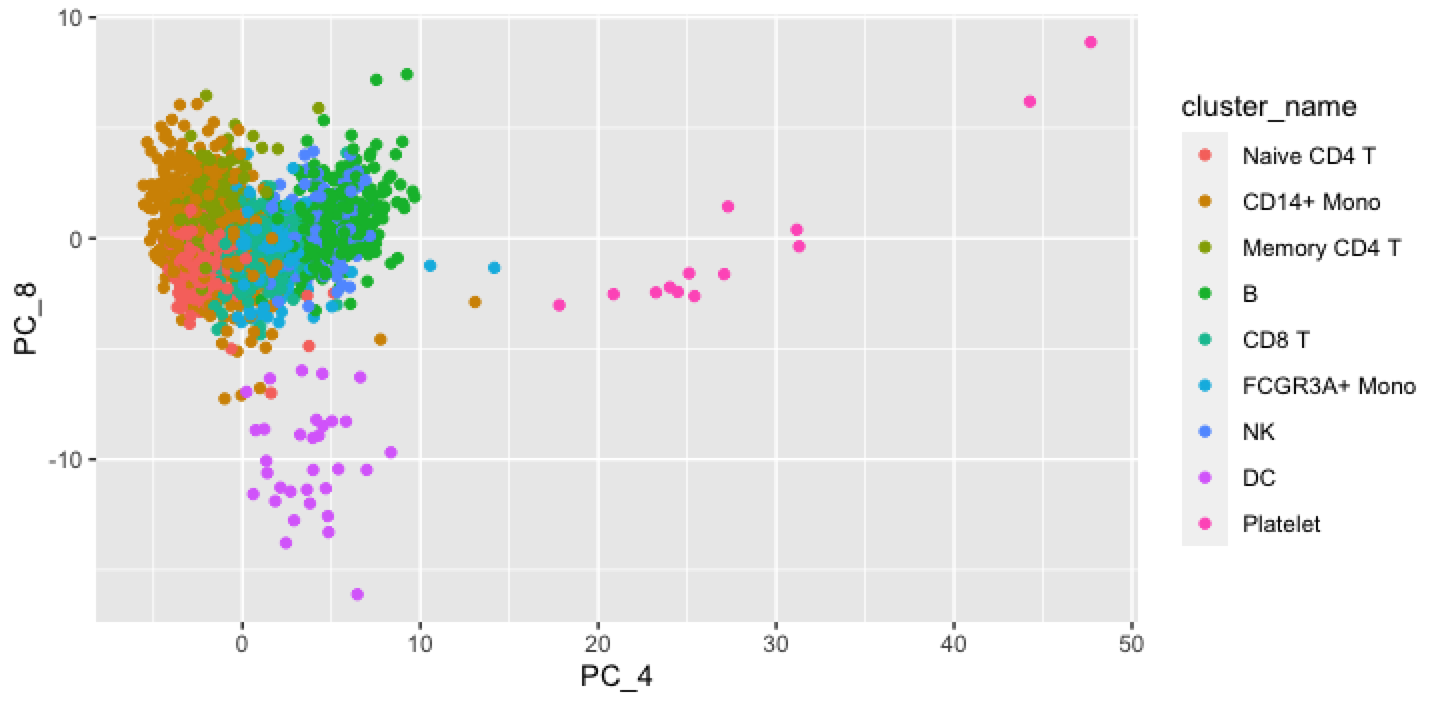
10
0
-10
PC_8
0
10
20
30
40
50
PC_4
cluster_name
Naive CD4 T
CD14+ Mono
Memory CD4 T
B
CD8 T
FCGR3A+ Mono
NK
DC
Platelet

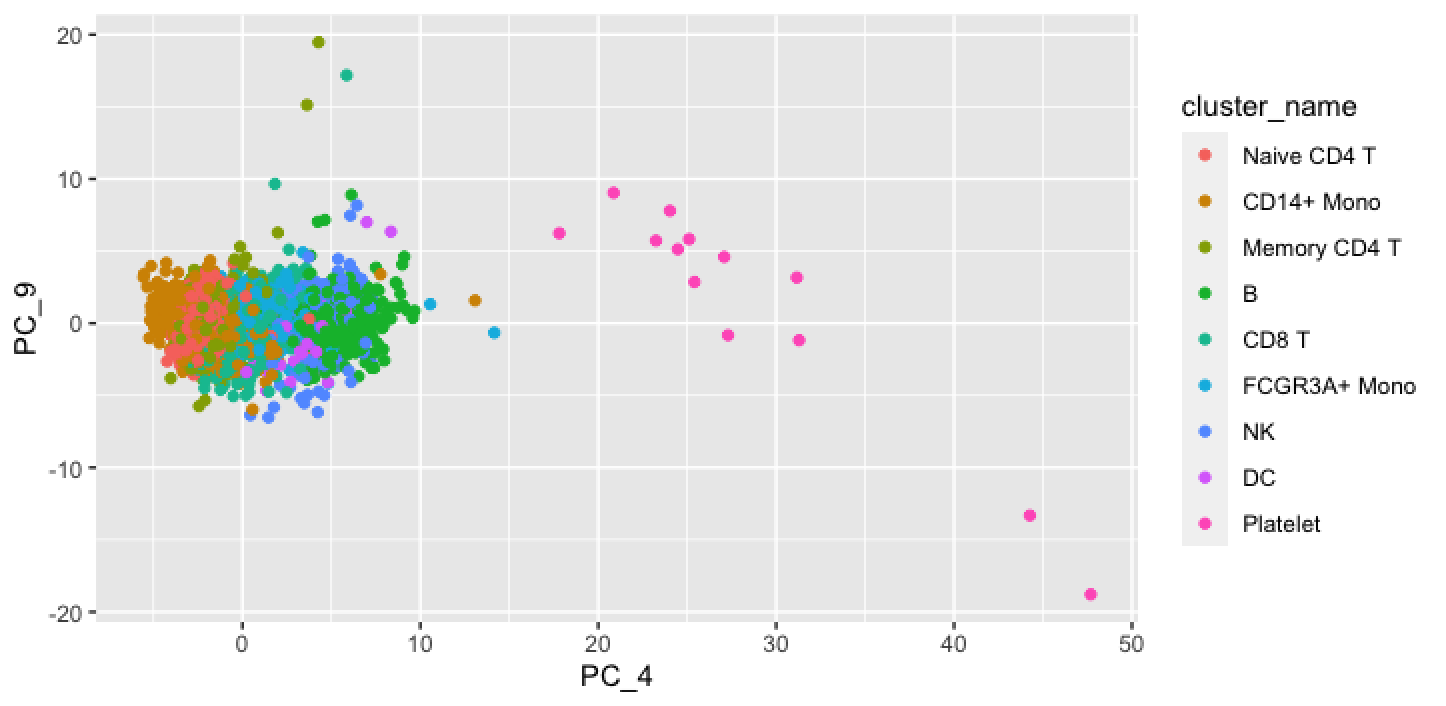
20
10
0
-10
-20
PC_9
0
10
20
30
40
50
PC_4
cluster_name
Naive CD4 T
CD14+ Mono
Memory CD4 T
B
CD8 T
FCGR3A+ Mono
NK
DC
Platelet

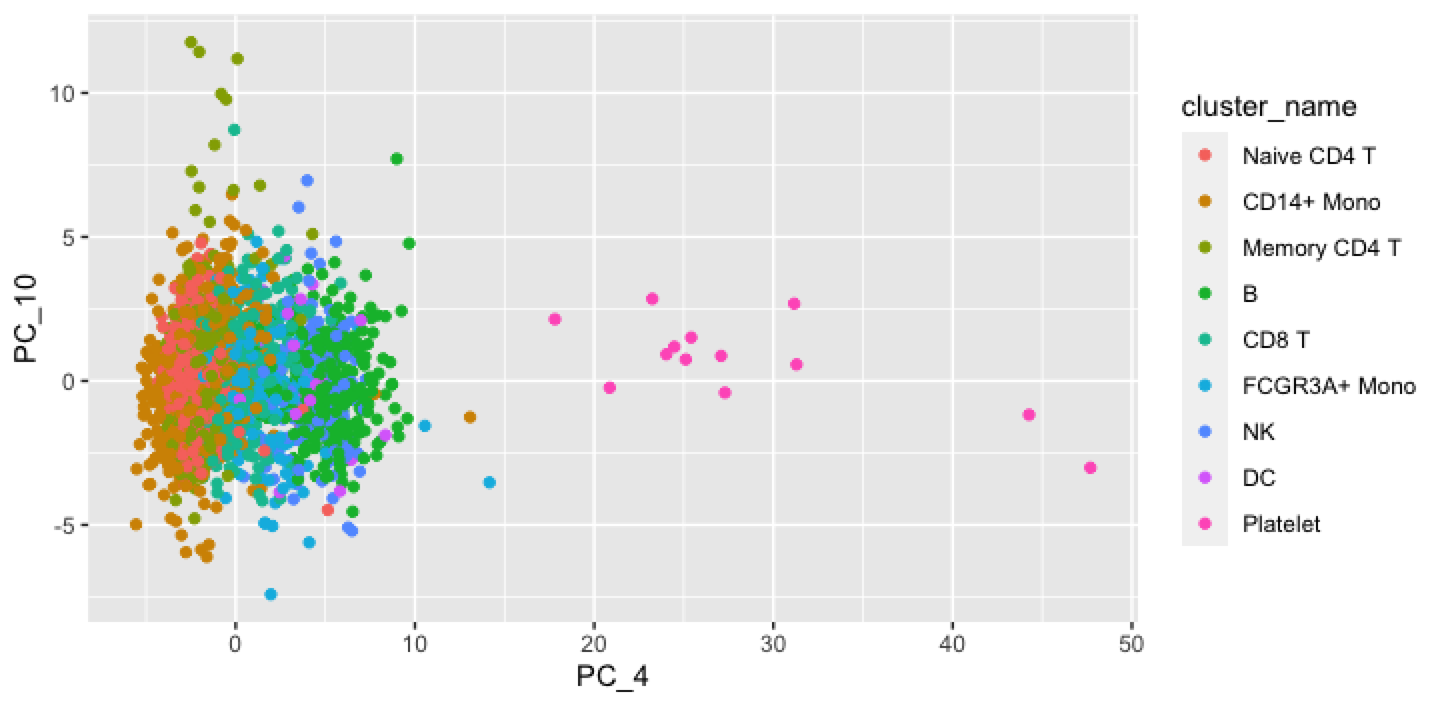

10
5
0
-5
PC_10
0
10
20
30
40
50
PC_4
cluster_name
Naive CD4 T
CD14+ Mono
Memory CD4 T
B
CD8 T
FCGR3A+ Mono
NK
DC
Platelet

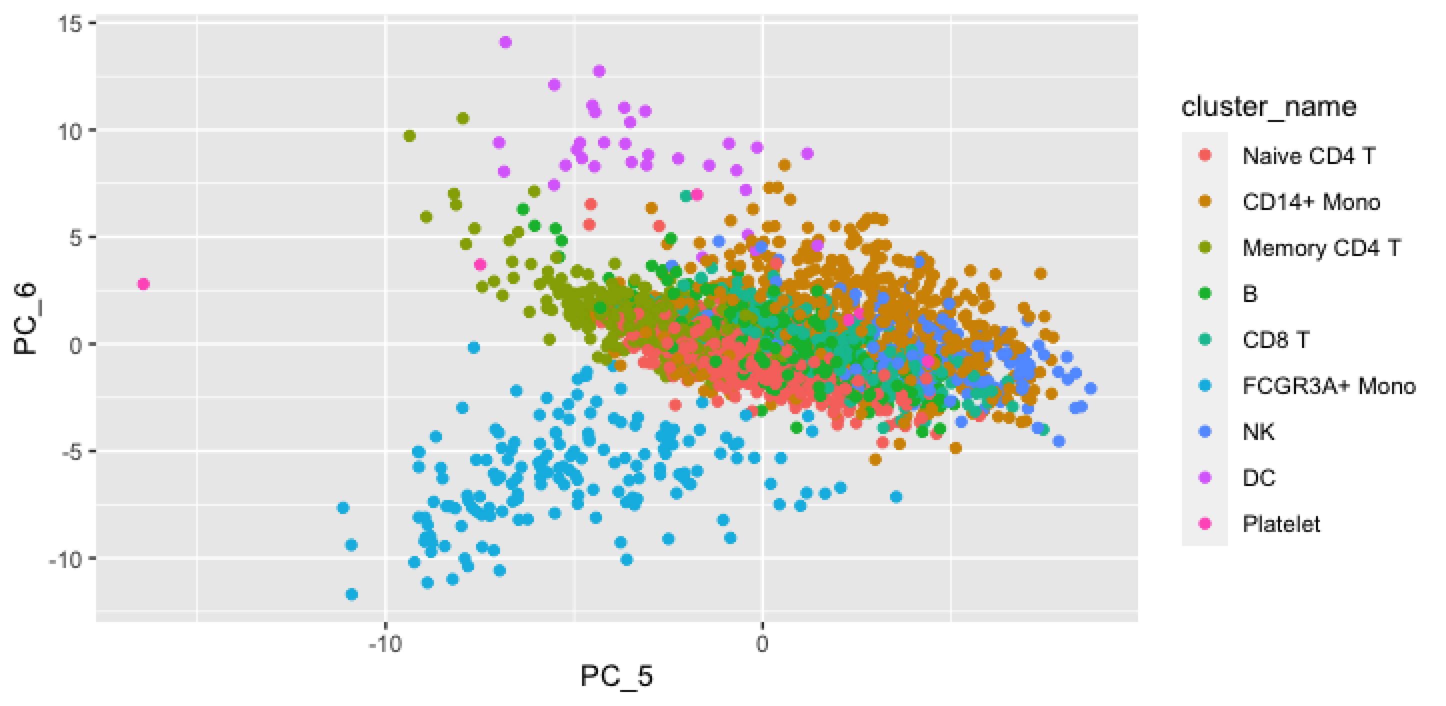

PC_6
PC_5
15
10
5
0
-5
-10
-10
0
cluster_name
Naive CD4 T
CD14+ Mono
Memory CD4 T
B
CD8 T
FCGR3A+ Mono
NK
DC
Platelet

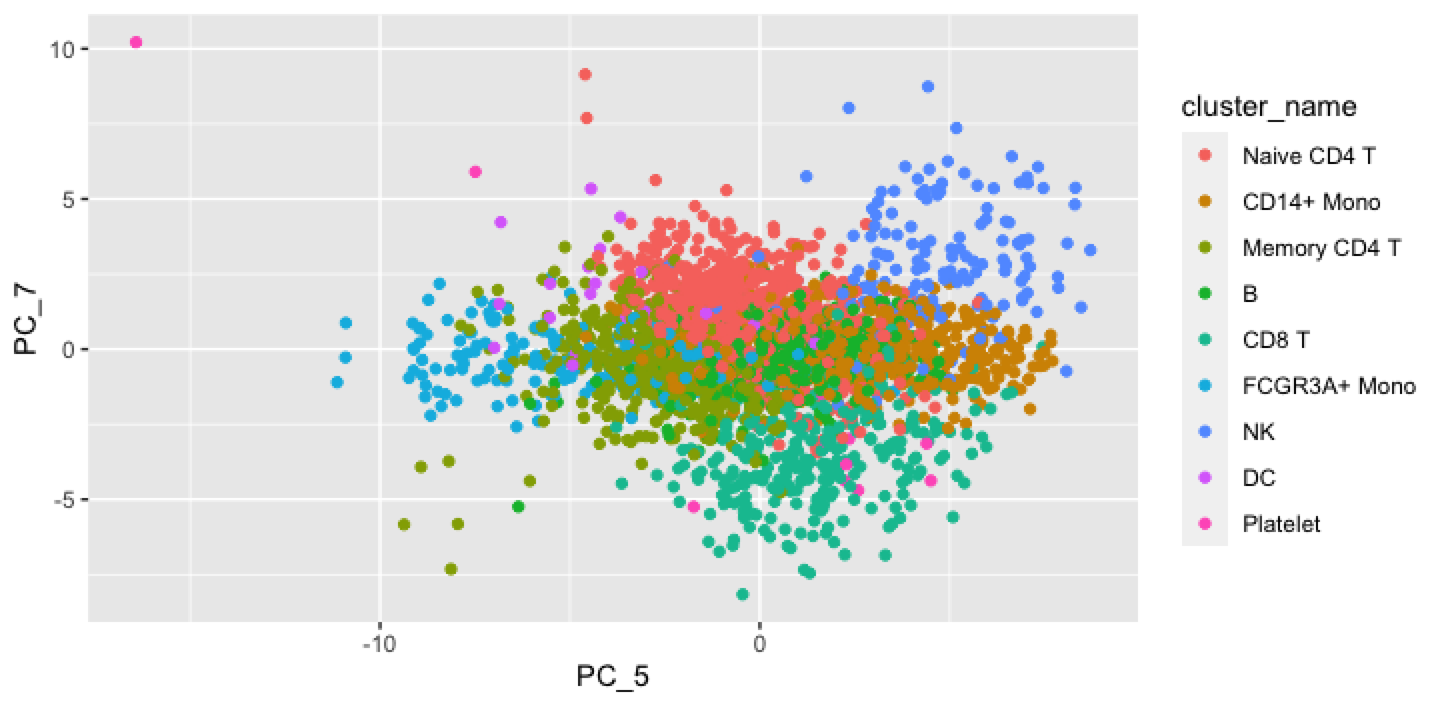

10
5
0
-5
PC_7
-10
0
PC_5
cluster_name
Naive CD4 T
CD14+ Mono
Memory CD4 T
B
CD8 T
FCGR3A+ Mono
NK
DC
Platelet

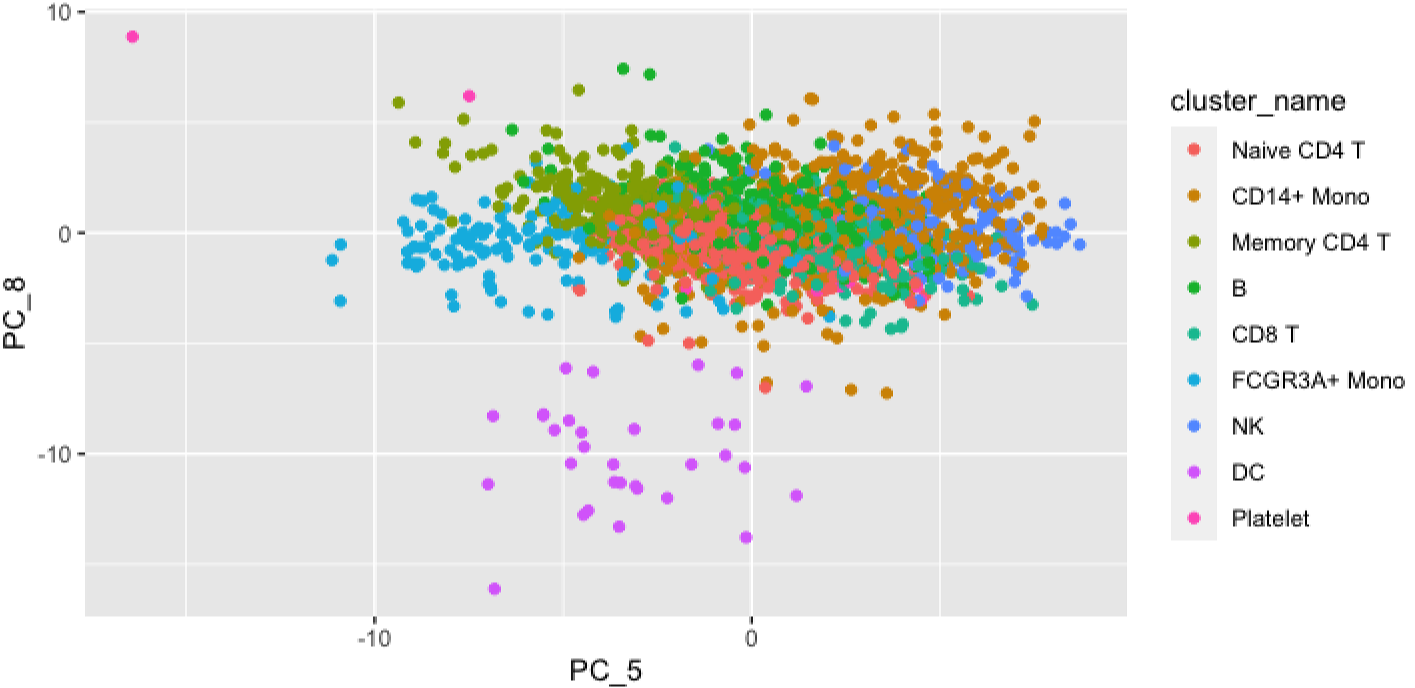
10
0
-10
-10
0
PC_5
PC_8
cluster_name
Naive CD4 T
CD14+ Mono
Memory CD4 T
B
CD8 T
FCGR3A+ Mono
NK
DC
Platelet

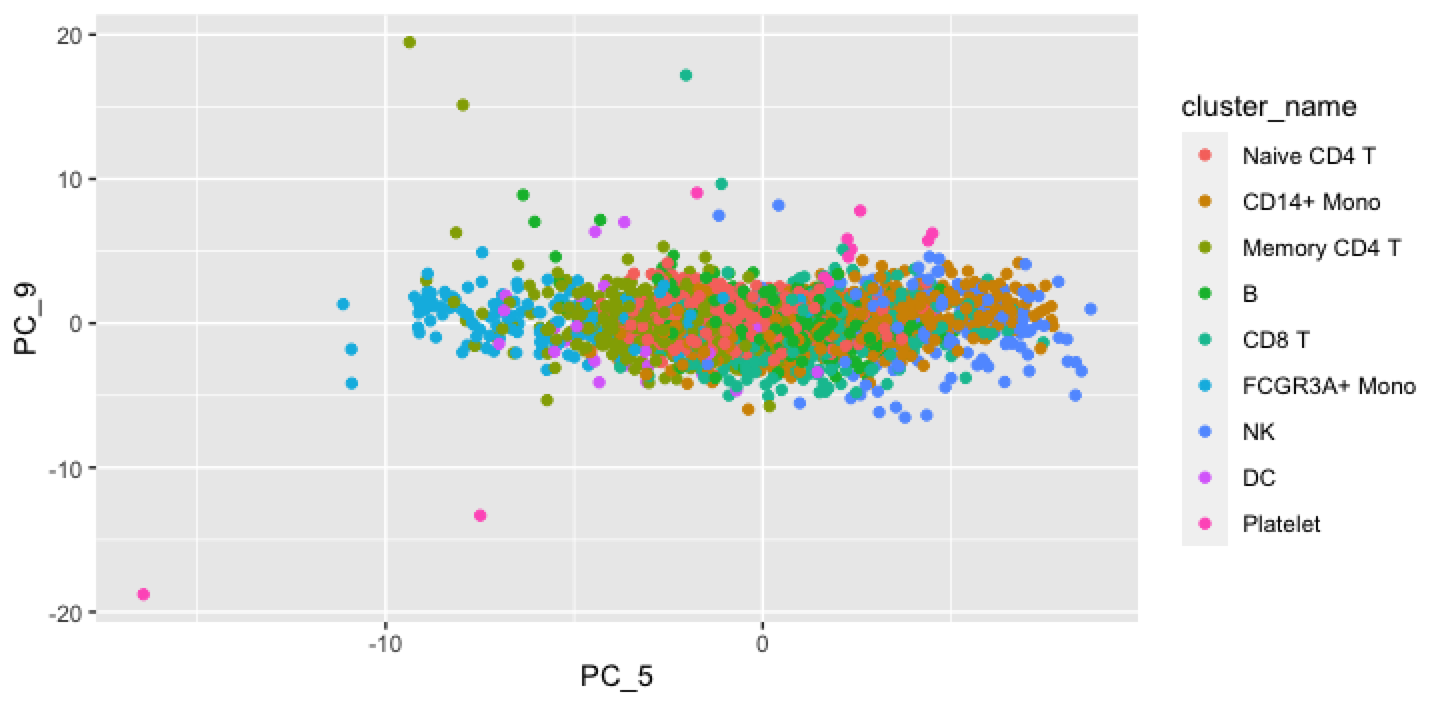
20
10
0
-10
-20
-10
0
PC_9
PC_5
cluster_name
Naive CD4 T
CD14+ Mono
Memory CD4 T
B
CD8 T
FCGR3A+ Mono
NK
DC
Platelet

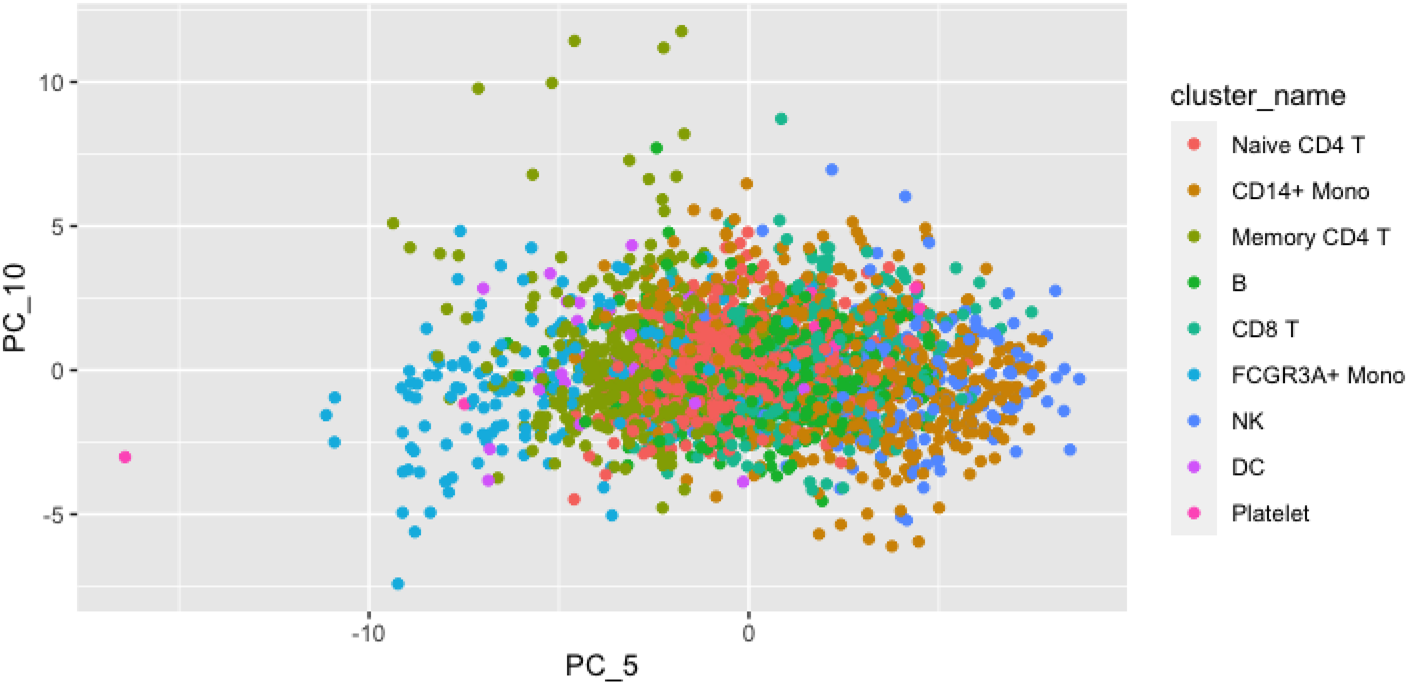

10
5
0
-5
PC_10
-10
0
PC_5
cluster_name
Naive CD4 T
CD14+ Mono
Memory CD4 T
B
CD8 T
FCGR3A+ Mono
NK
DC
Platelet

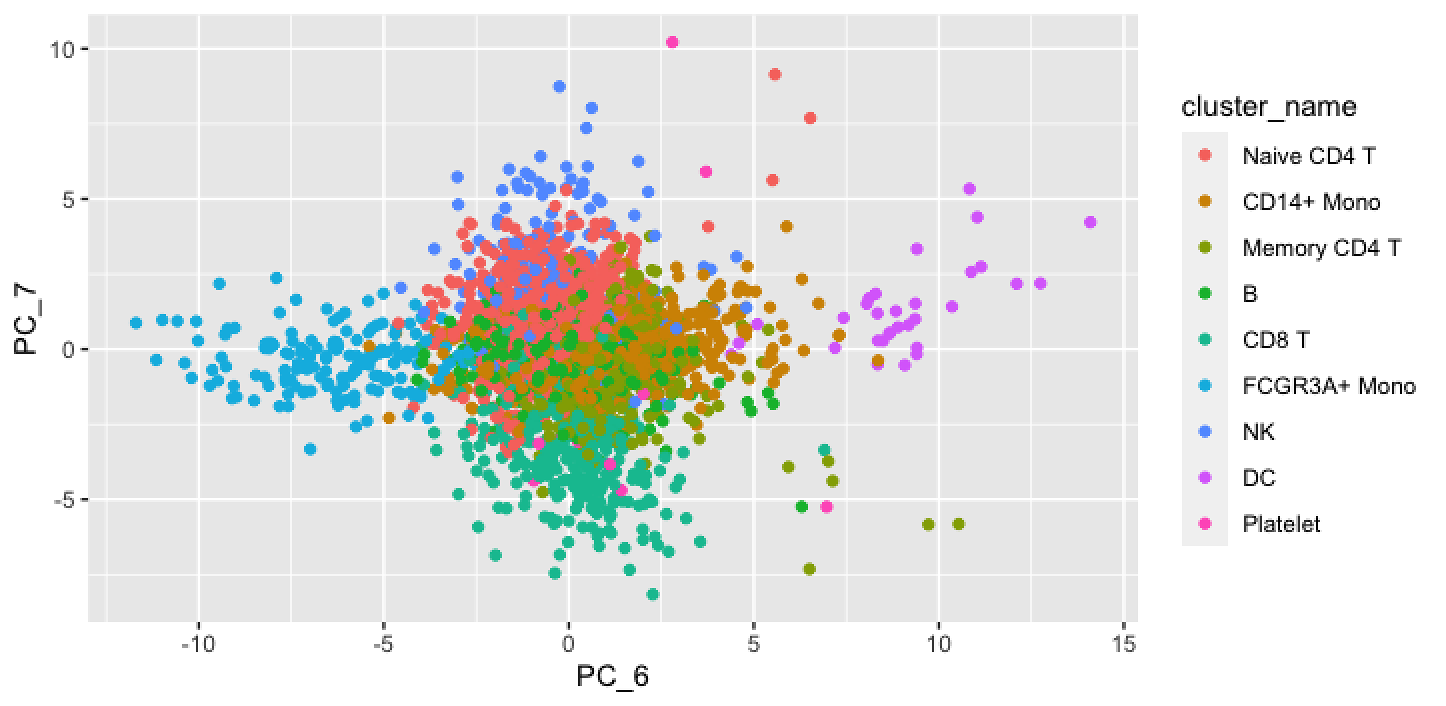

cluster_name
Naive CD4 T
CD14+ Mono
Memory CD4 T
B
CD8 T
FCGR3A+ Mono
NK
DC
Platelet
PC_7
PC_6
10
5
0
-5
-10
15

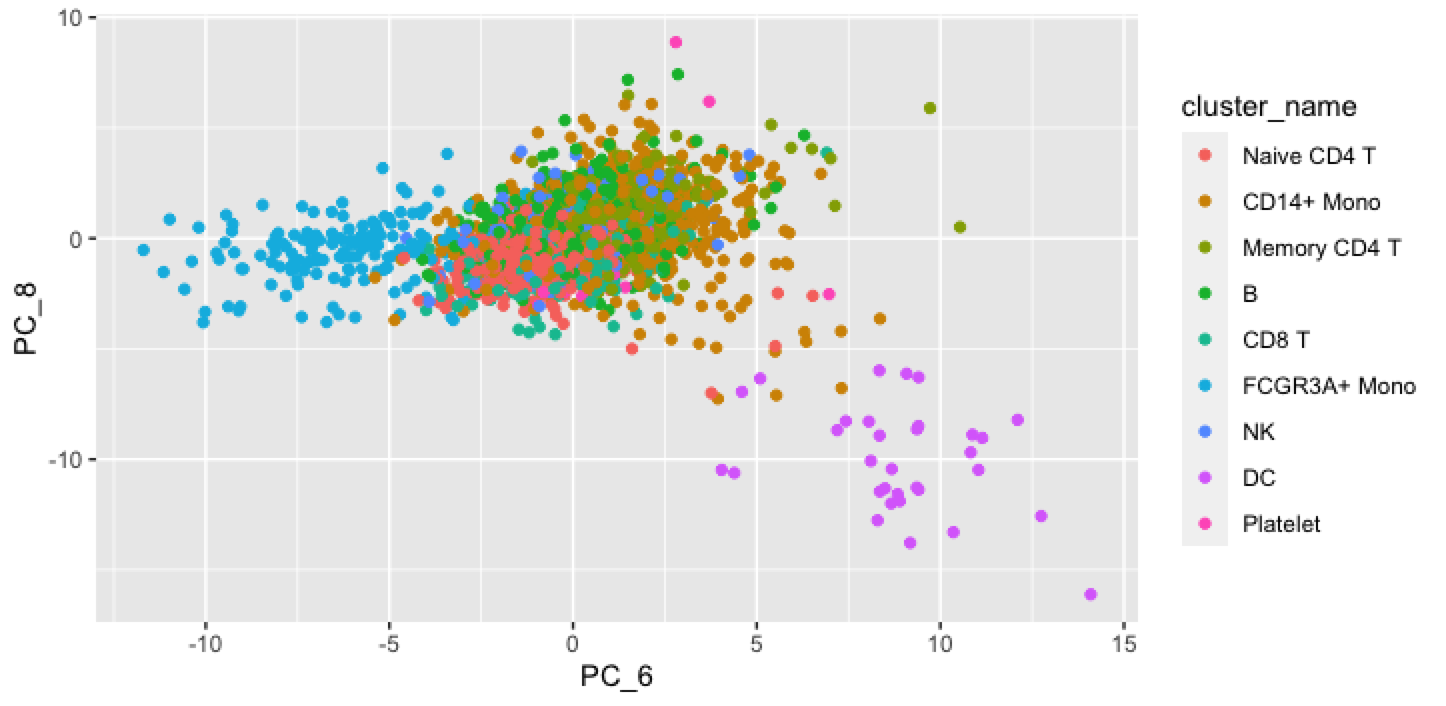
PC_8
10
0
-10
PC_6
-10
-5
0
5
10
15
cluster_name
Naive CD4 T
CD14+ Mono
Memory CD4 T
B
CD8 T
FCGR3A+ Mono
NK
DC
Platelet

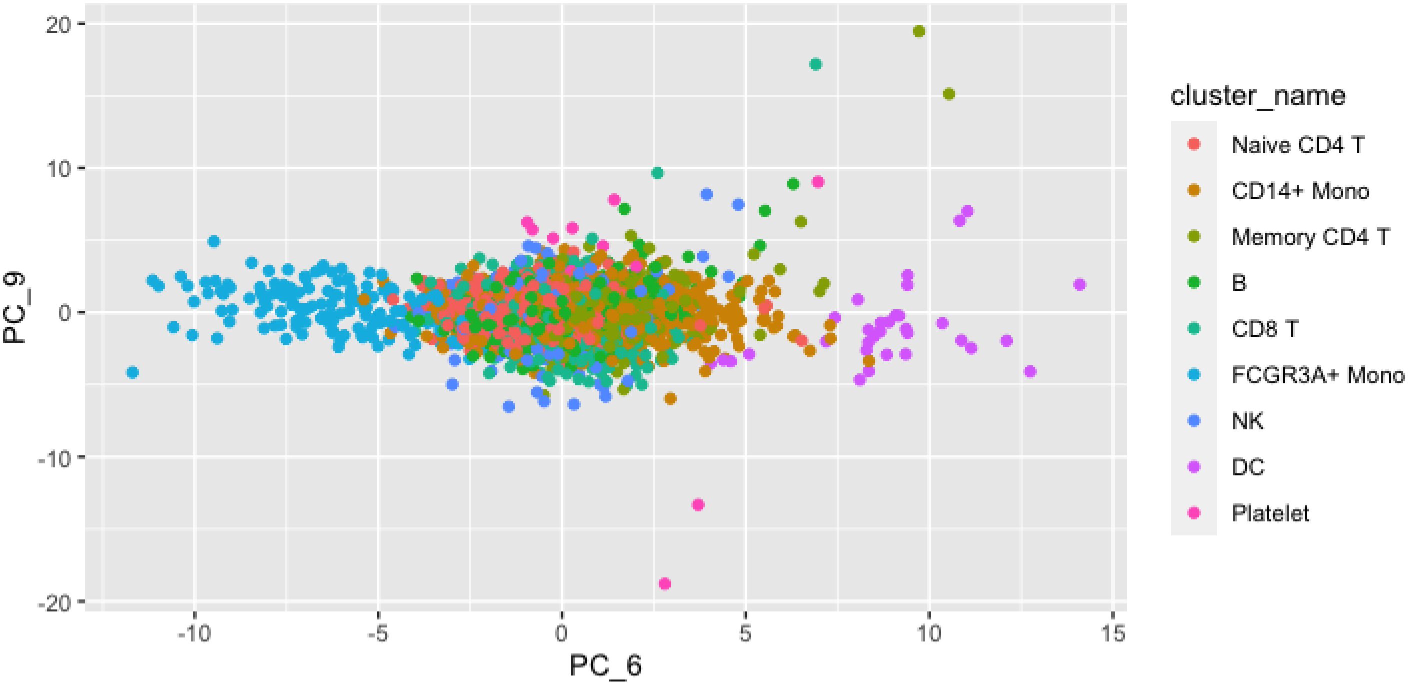

20
10
0
-10
-20
PC_9
-10
-5
0
5
10
15
PC_6
cluster_name
Naive CD4 T
CD14+ Mono
Memory CD4 T
B
CD8 T
FCGR3A+ Mono
NK
DC
Platelet

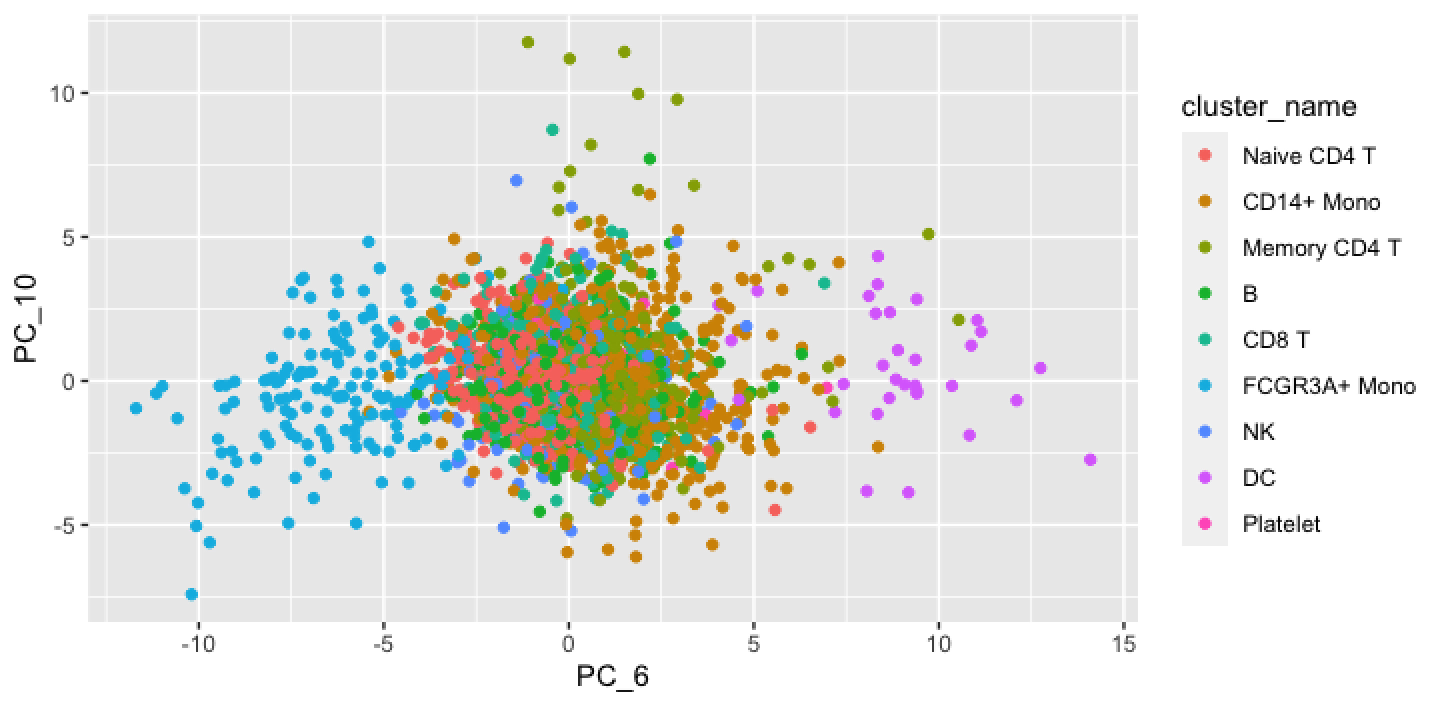
10
5
0
-5
PC_10
-10
-5
0
5
10
15
PC_6
cluster_name
Naive CD4 T
CD14+ Mono
Memory CD4 T
B
CD8 T
FCGR3A+ Mono
NK
DC
Platelet

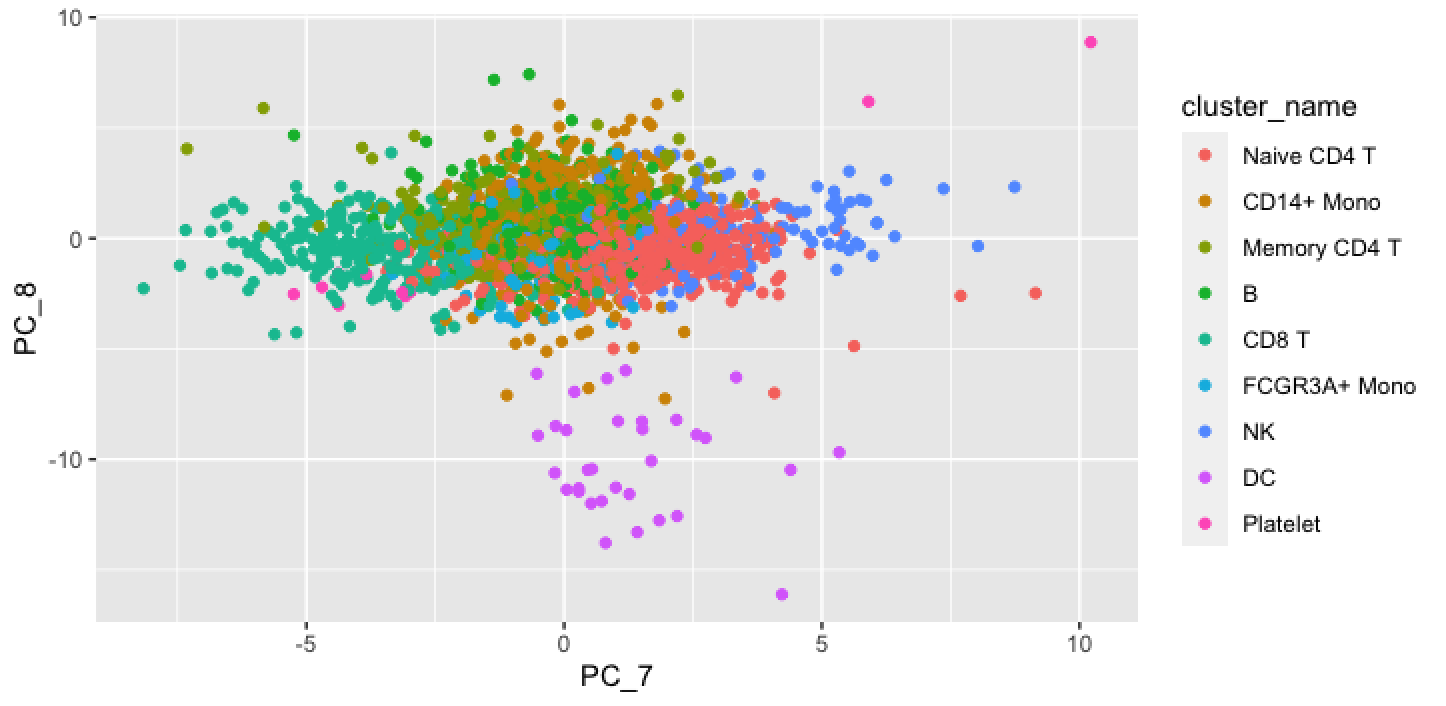

10
0
-10
PC_8
-5
0
5
10
PC_7
cluster_name
Naive CD4 T
CD14+ Mono
Memory CD4 T
B
CD8 T
FCGR3A+ Mono
NK
DC
Platelet

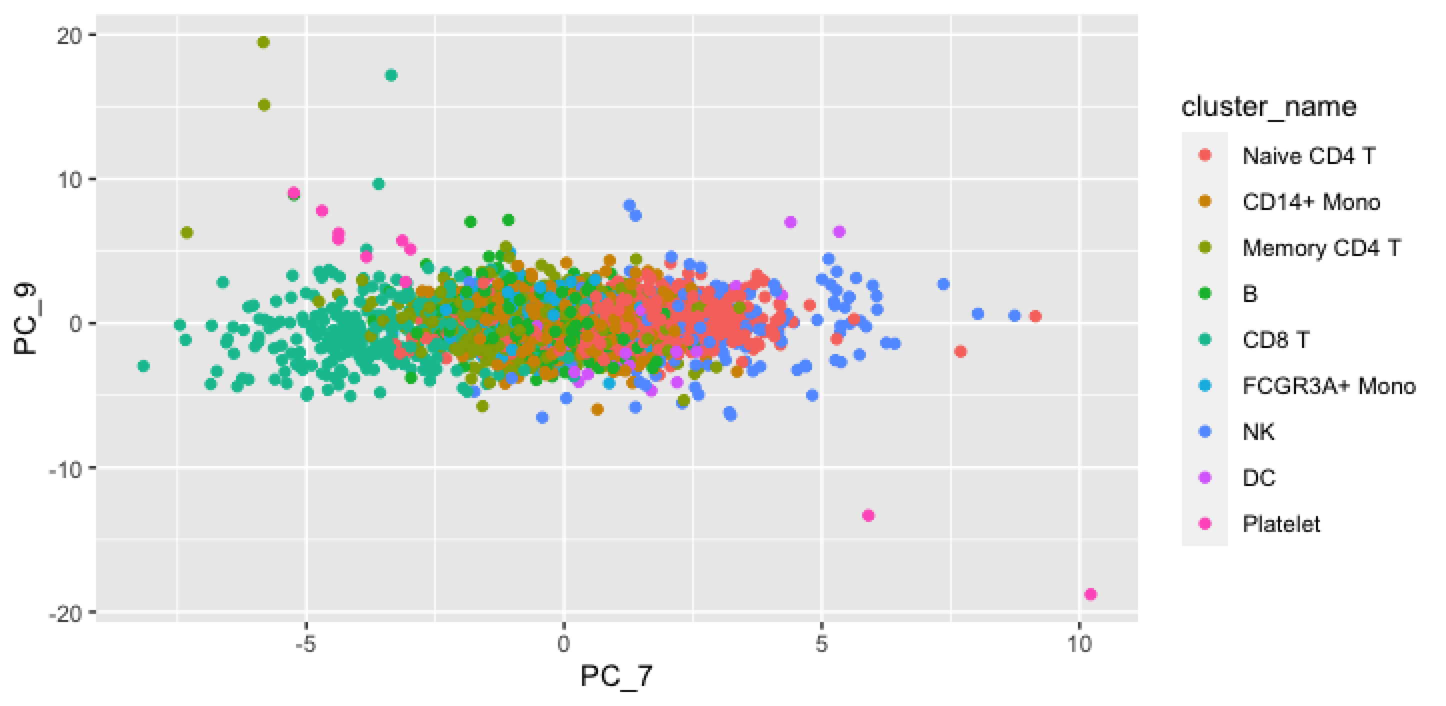
20
10
0
-10
-20
PC_9
-5
0
5
10
PC_7
cluster_name
Naive CD4 T
CD14+ Mono
Memory CD4 T
B
CD8 T
FCGR3A+ Mono
NK
DC
Platelet

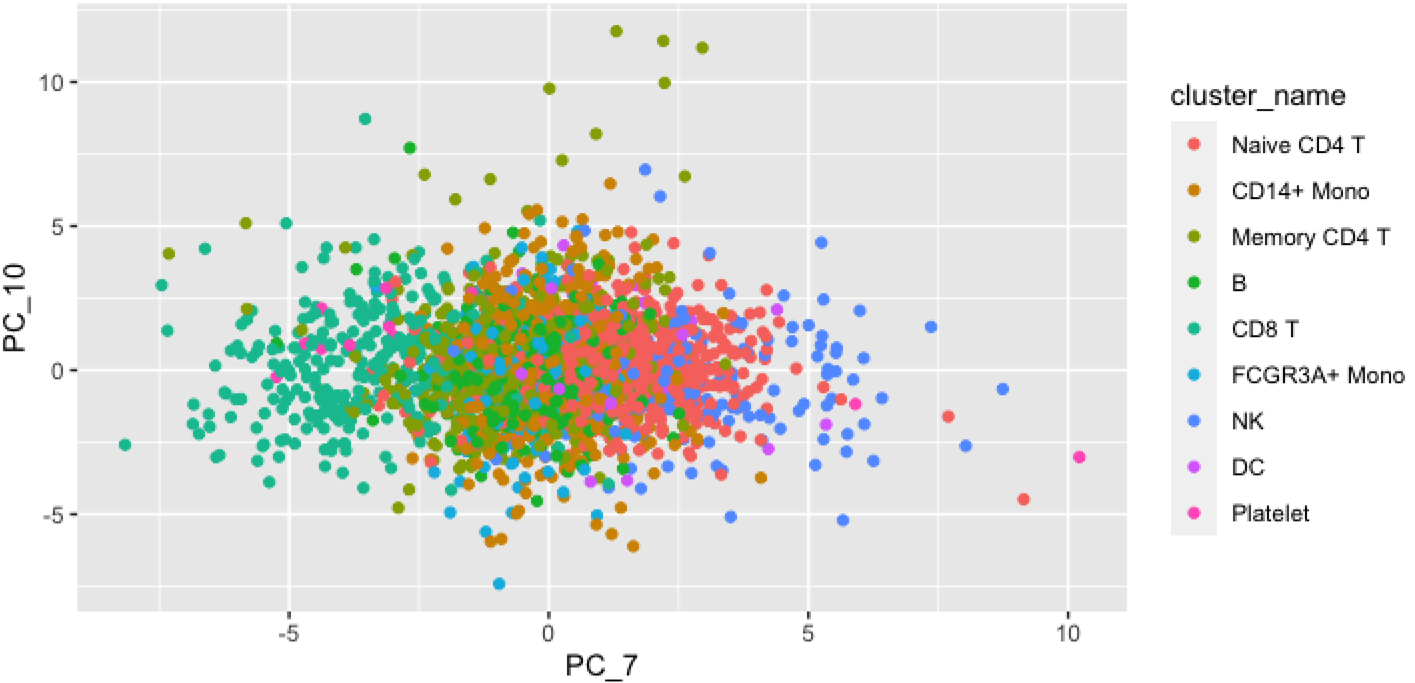

10
5
0
-5
PC_10
-5
0
5
10
PC_7
cluster_name
Naive CD4 T
CD14+ Mono
Memory CD4 T
B
CD8 T
FCGR3A+ Mono
NK
DC
Platelet

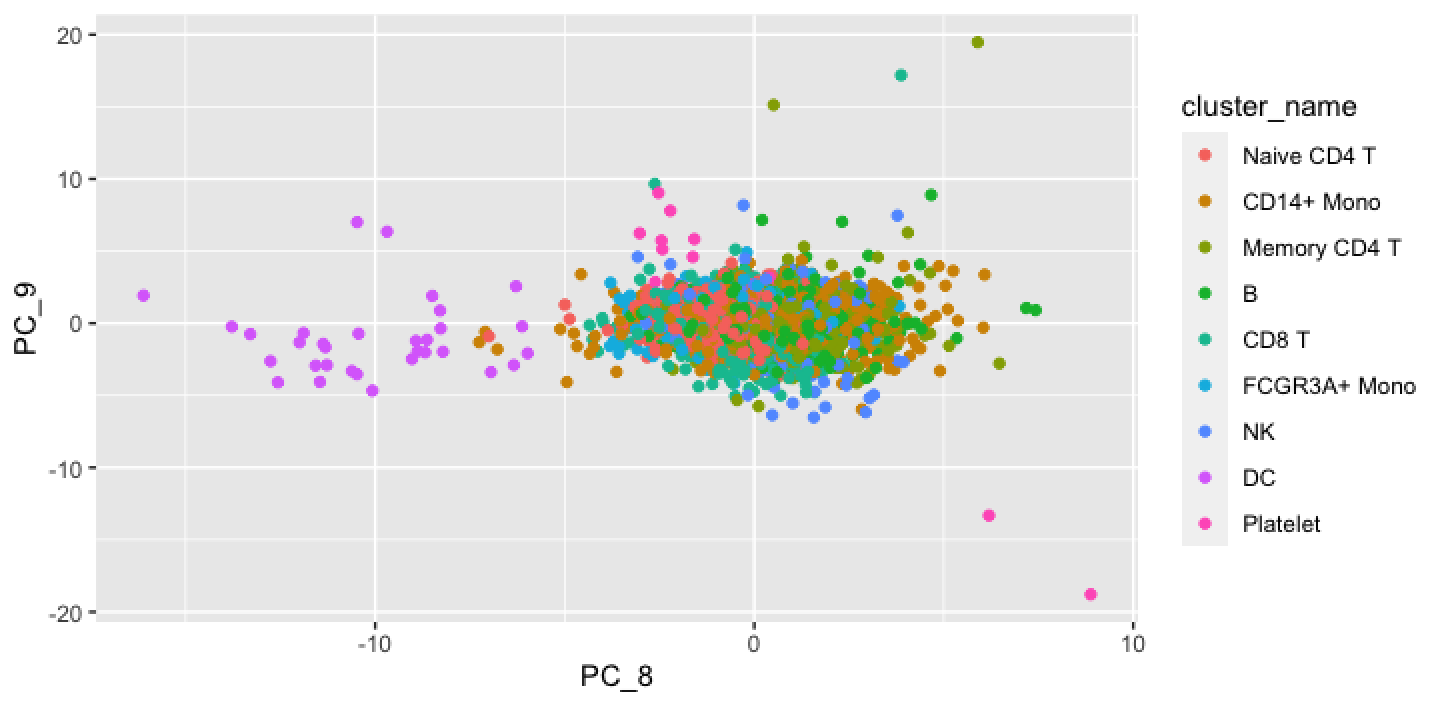

20
10
0
-10
-20
PC_9
-10
0
10
PC_8
cluster_name
Naive CD4 T
CD14+ Mono
Memory CD4 T
B
CD8 T
FCGR3A+ Mono
NK
DC
Platelet

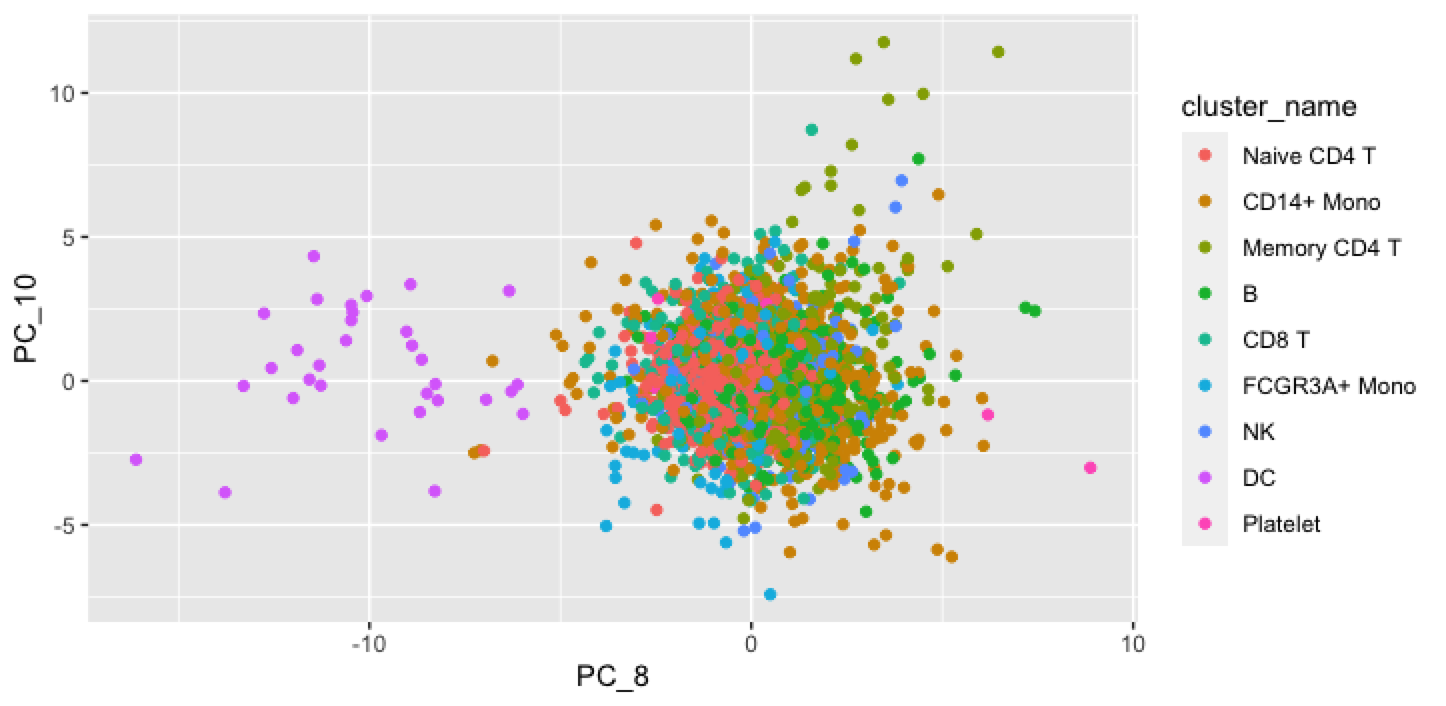

10
5
0
-5
PC_10
-10
0
10
PC_8
cluster_name
Naive CD4 T
CD14+ Mono
Memory CD4 T
B
CD8 T
FCGR3A+ Mono
NK
DC
Platelet

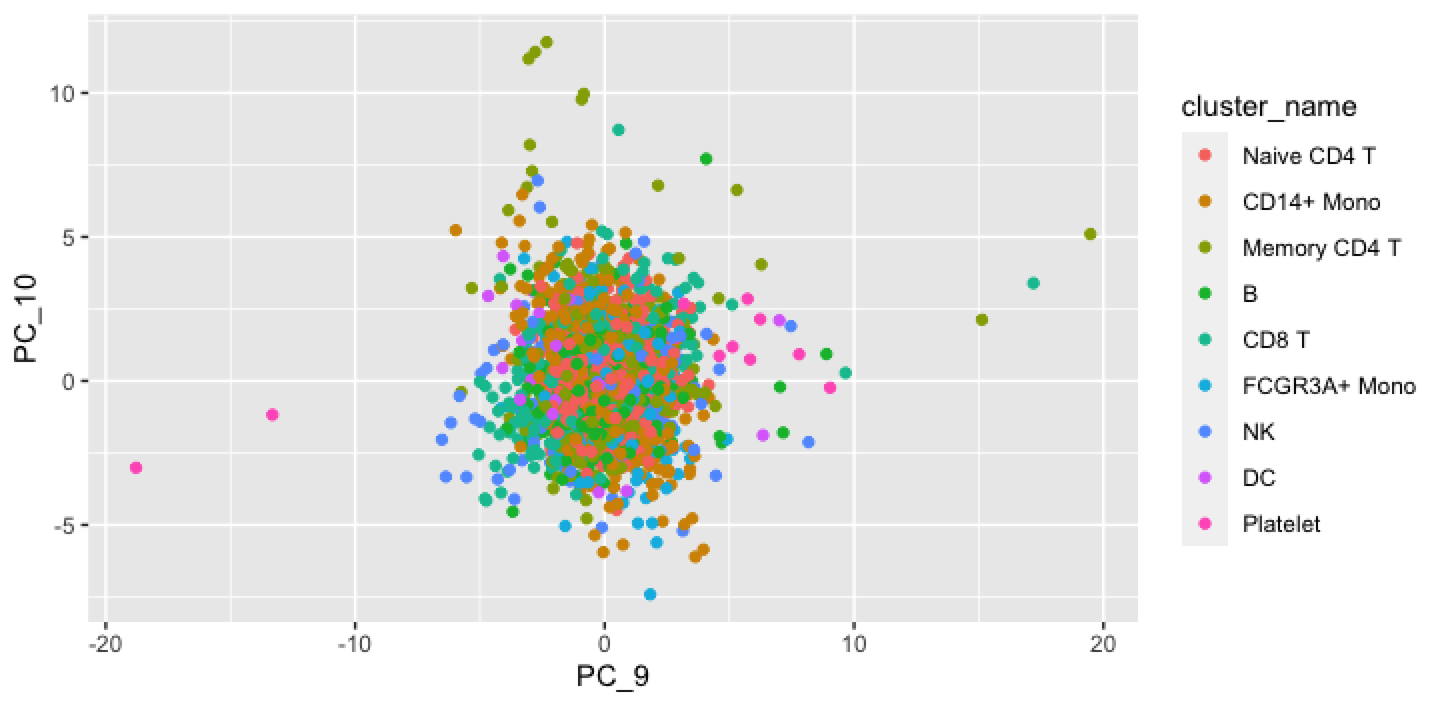
10
5
0
-5
PC_10
-20
-10
0
10
20
PC_9
cluster_name
Naive CD4 T
CD14+ Mono
Memory CD4 T
B
CD8 T
FCGR3A+ Mono
NK
DC
Platelet
