## Supplementary figures and images for "Lifting the curse from high dimensional data: Automated projection pursuit clustering for the variety of biological data modalities"

### 1_2.png

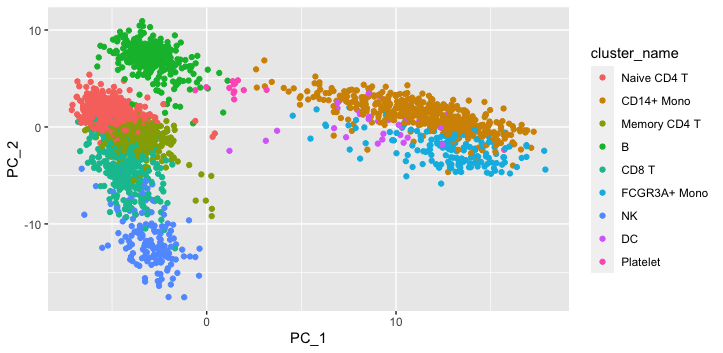

### 1_3.png

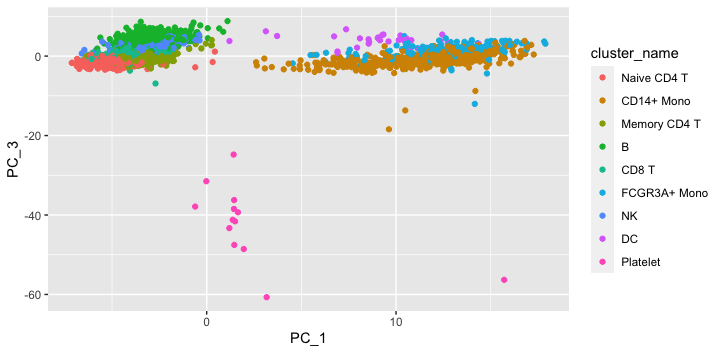

### 1_4.png

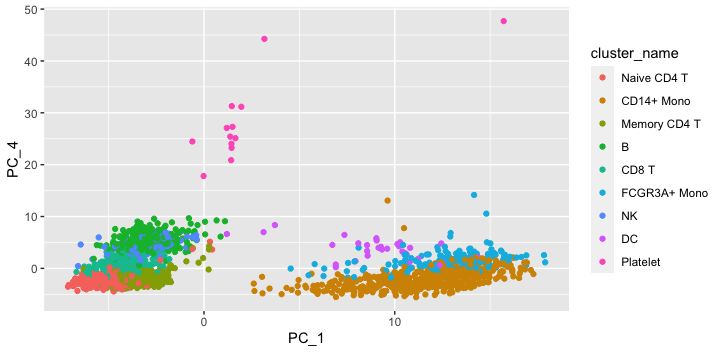

### 1_5.png

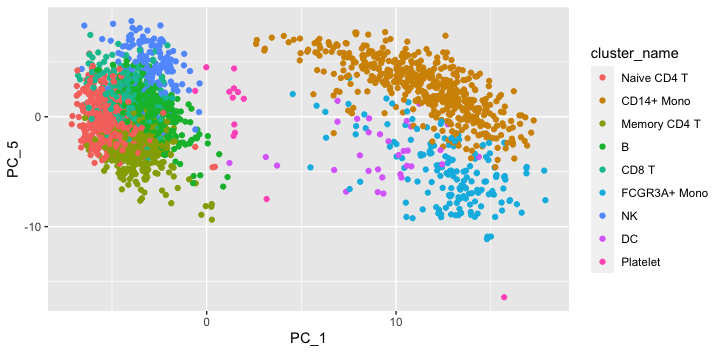

### 1_6.png

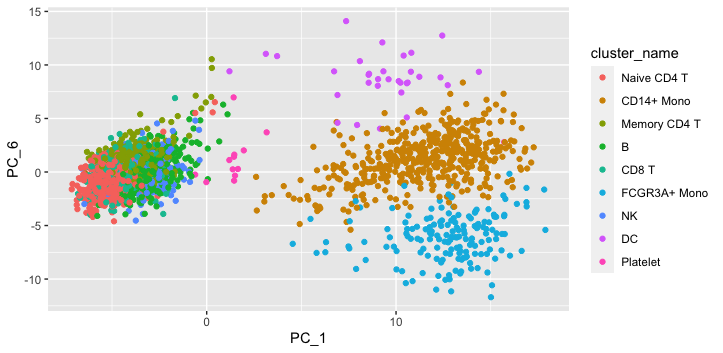

### 1_7.png

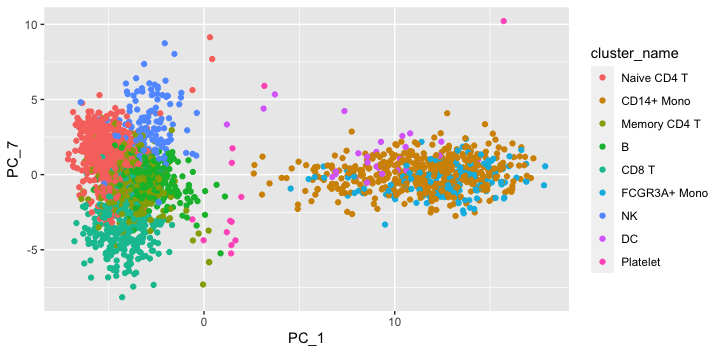

### 1_8.png

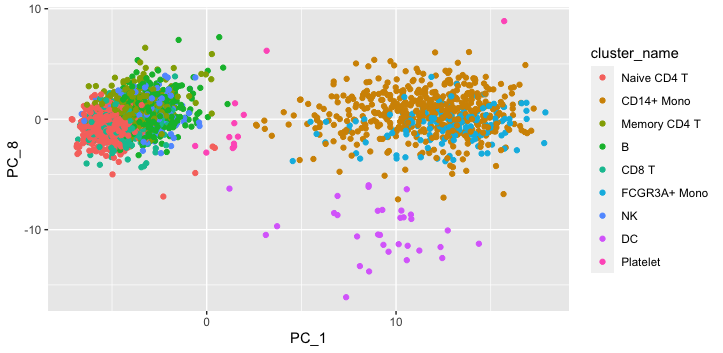

### 1_9.png

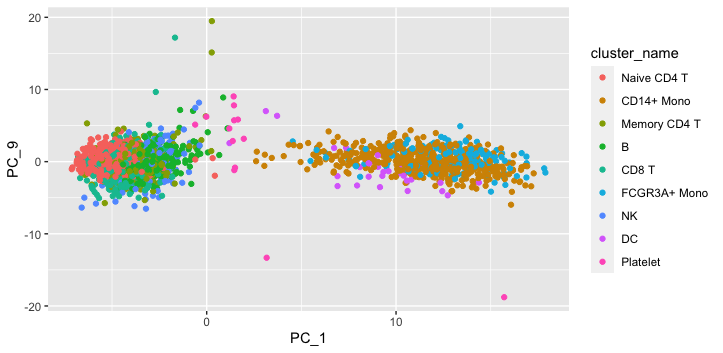

### 1_10.png

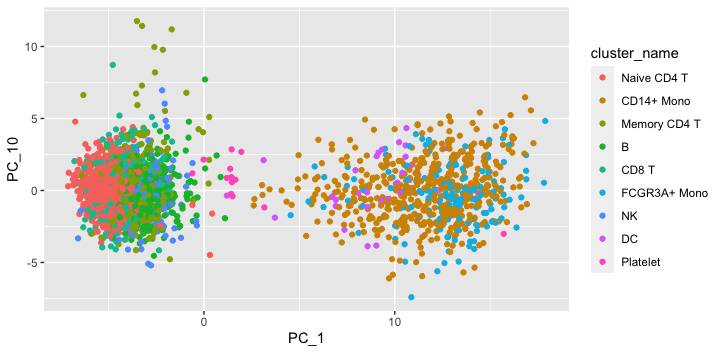

### 2_3.png

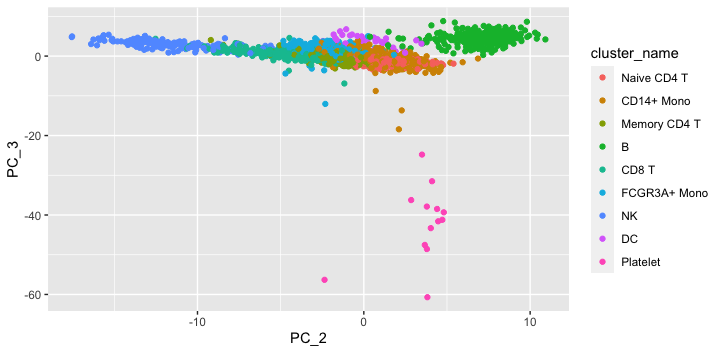

### 2_4.png

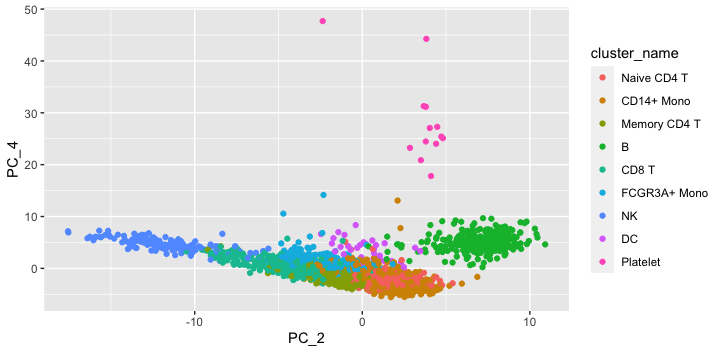

### 2_5.png

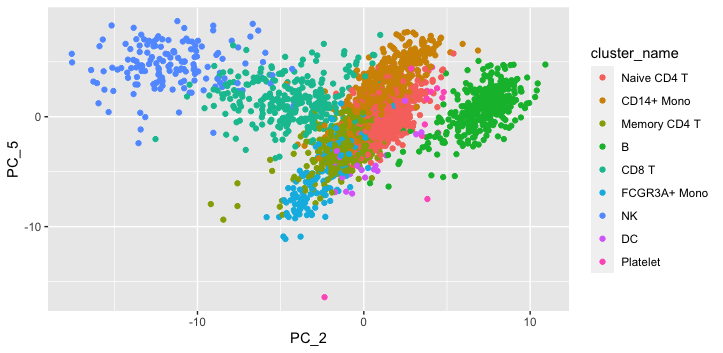

### 2_6.png

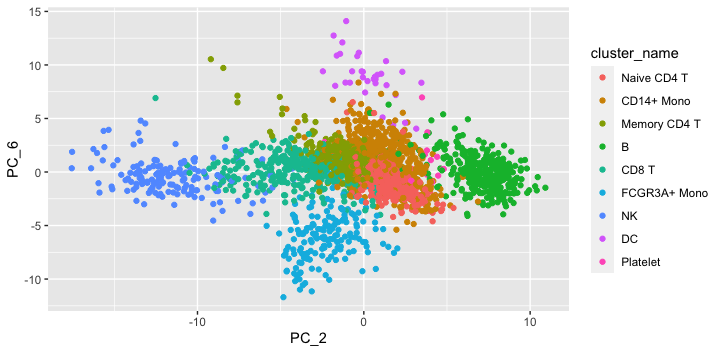

### 2_7.png

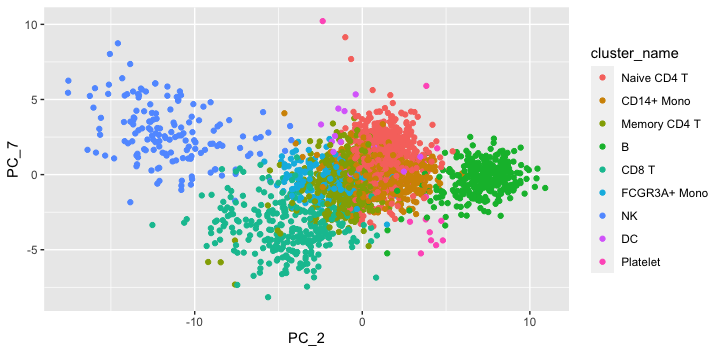

### 2_8.png

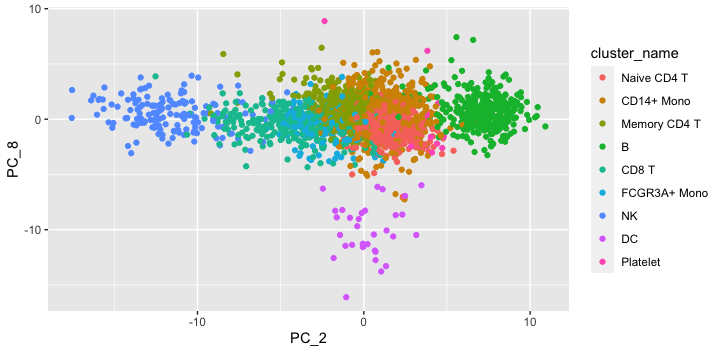

### 2_9.png

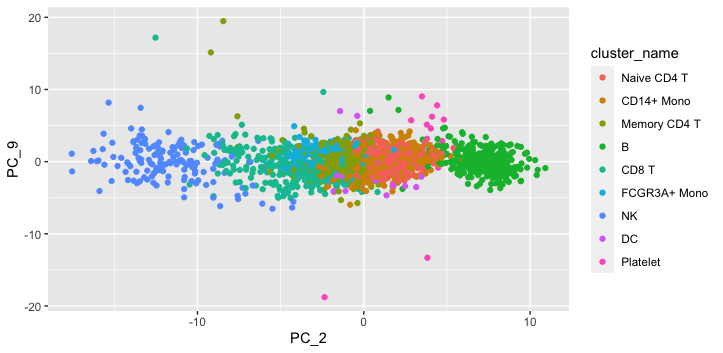

### 2_10.png

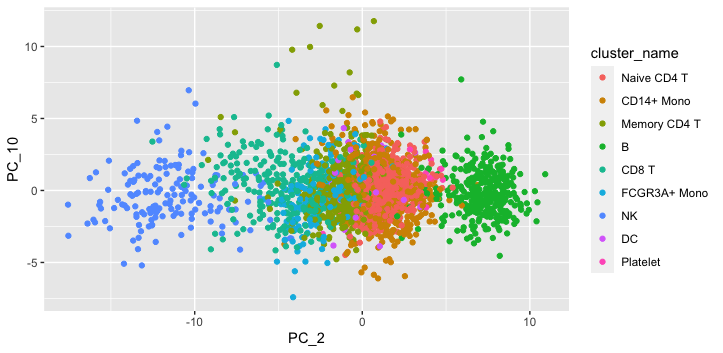

### 3_4.png

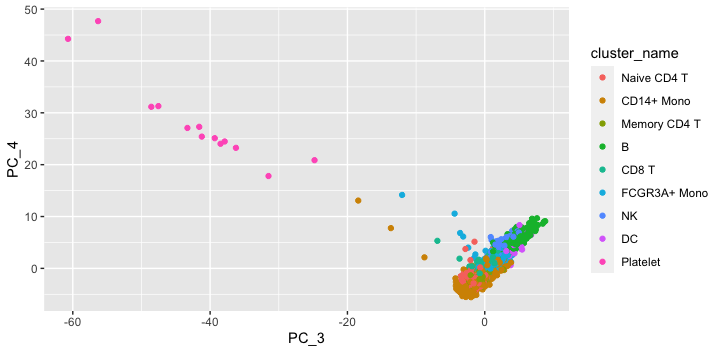

### 3_5.png

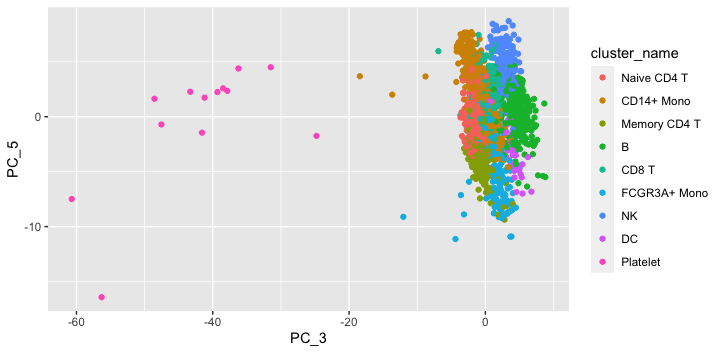

### 3_6.png

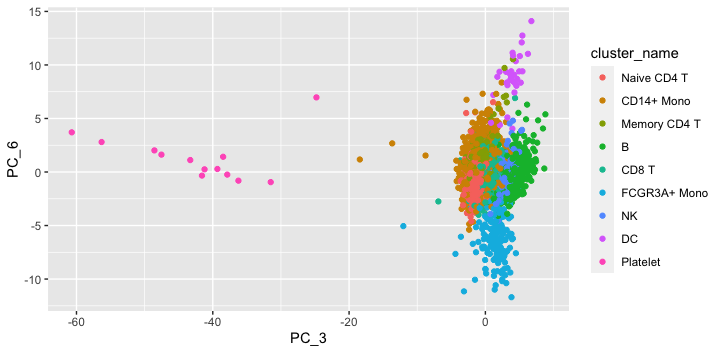

### 3_7.png

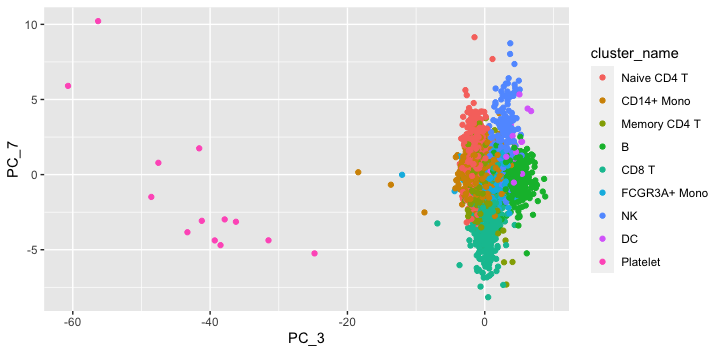

### 3_8.png

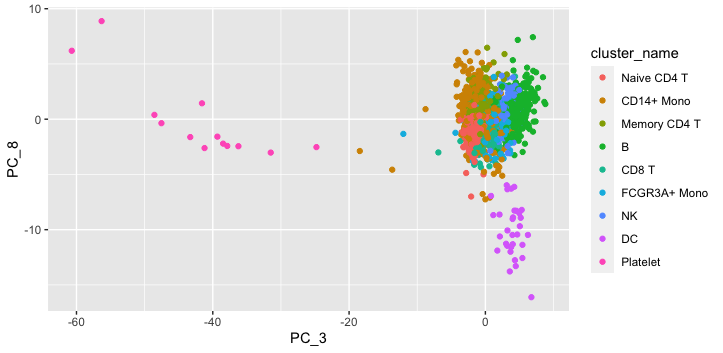

### 3_9.png

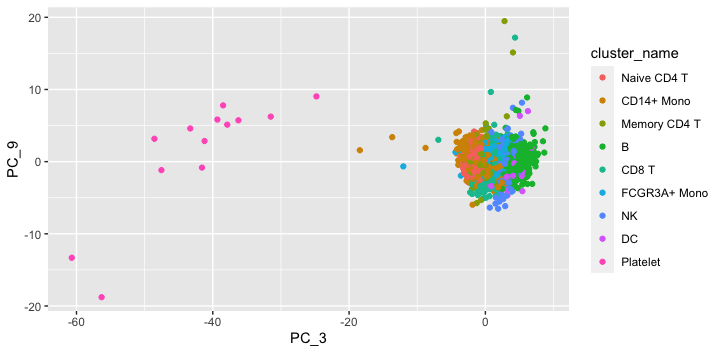

### 3_10.png

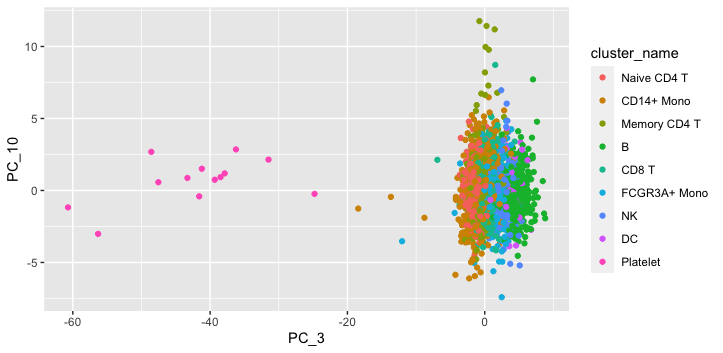

### 4_5.png

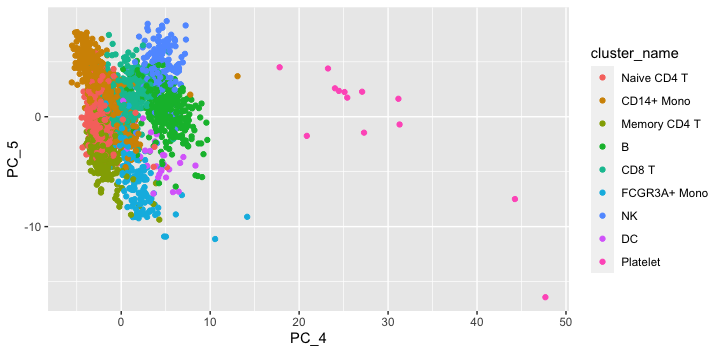

### 4_6.png

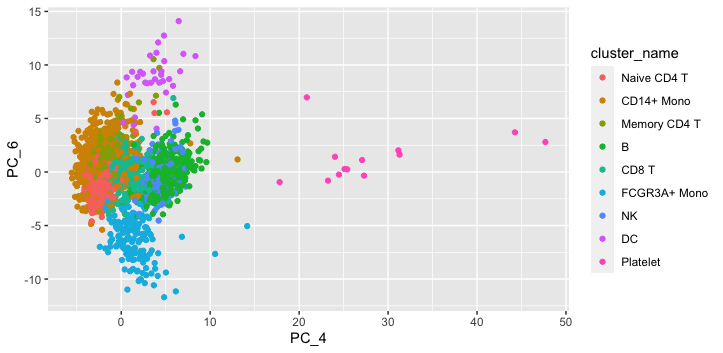

### 4_7.png

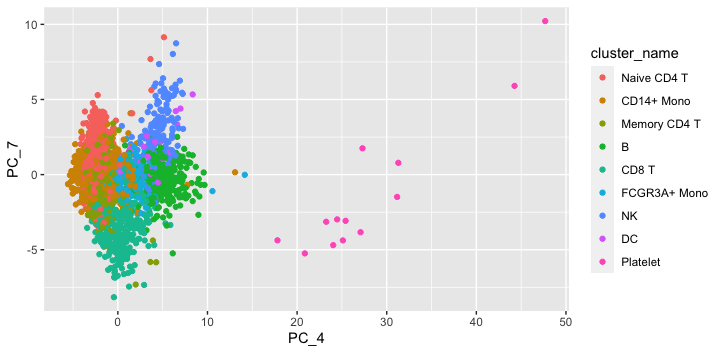

### 4_8.png

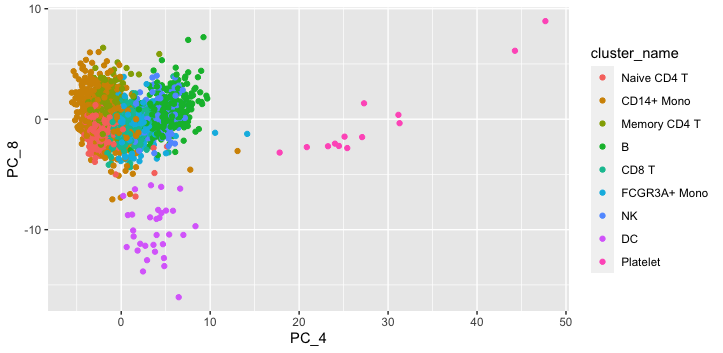

### 4_9.png

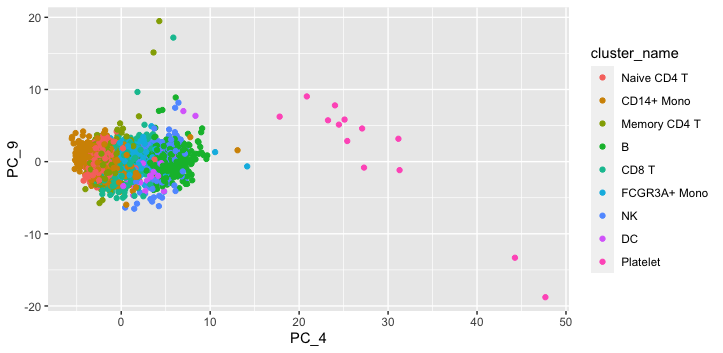

### 4_10.png

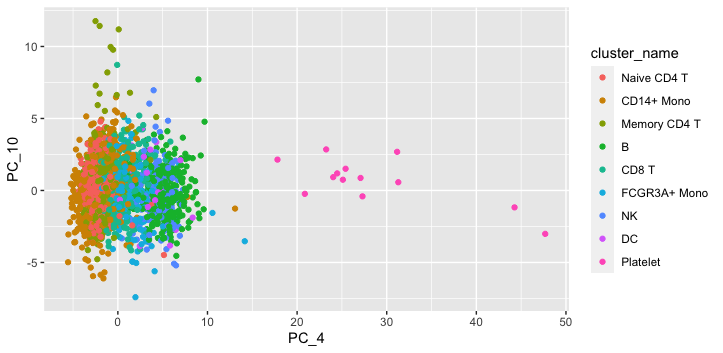

### 5_6.png

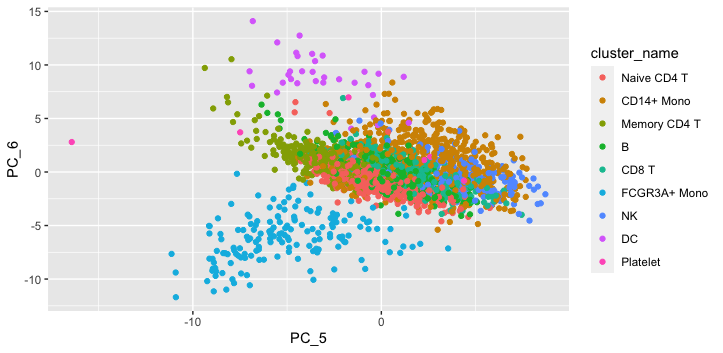

### 5_7.png

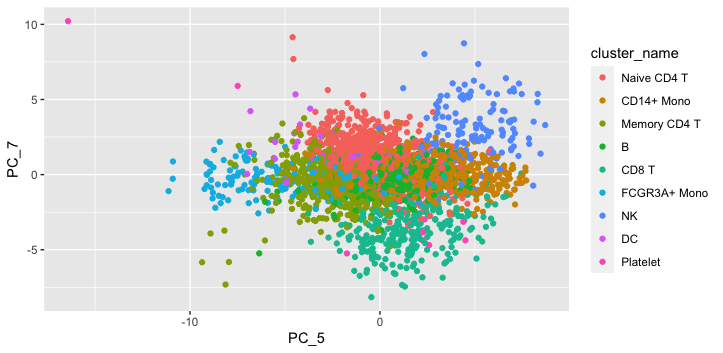

### 5_8.png

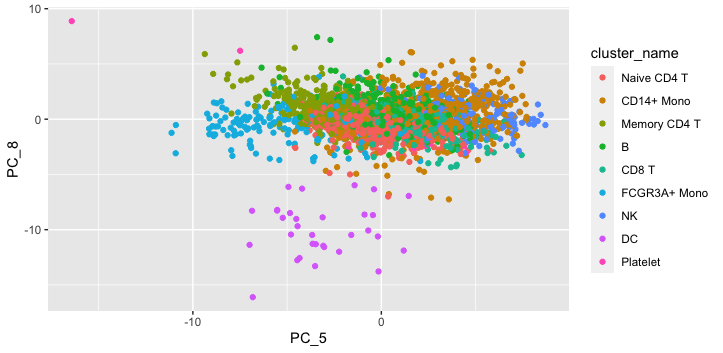

### 5_9.png

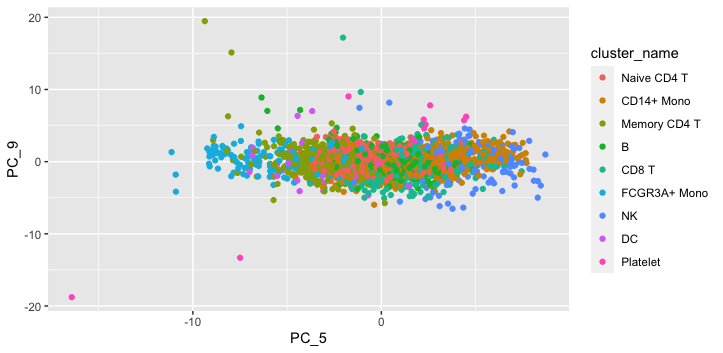

### 5_10.png

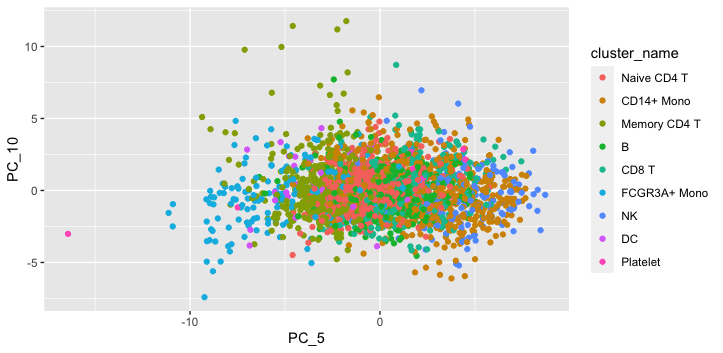

### 6_7.png

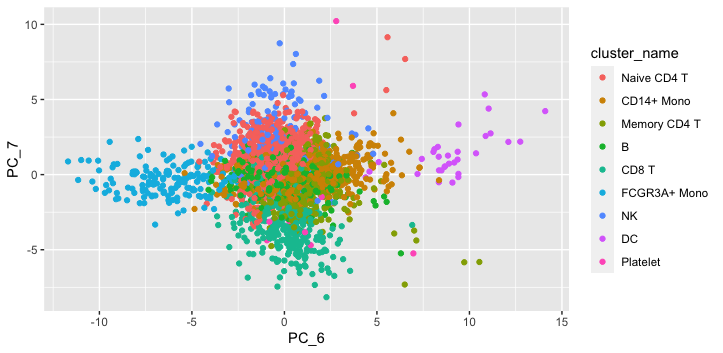

### 6_8.png

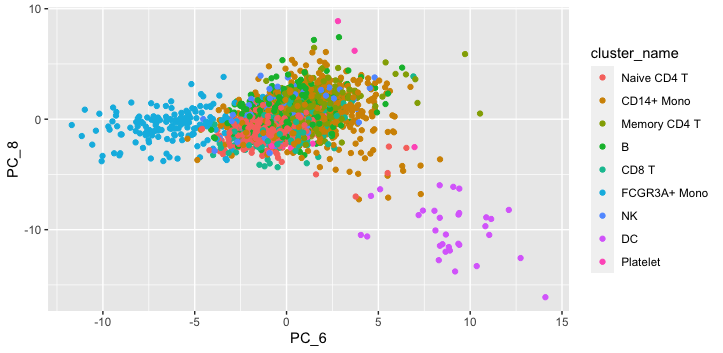

### 6_9.png

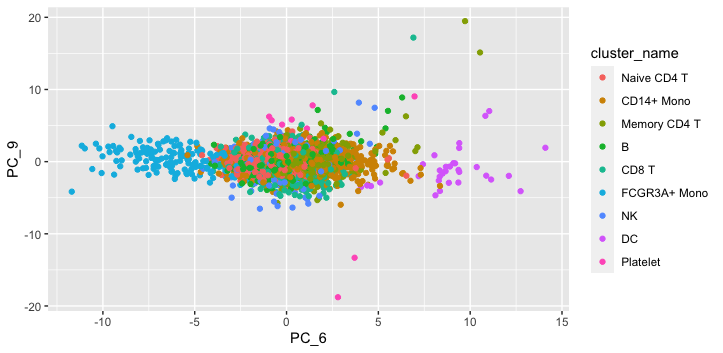

### 6_10.png

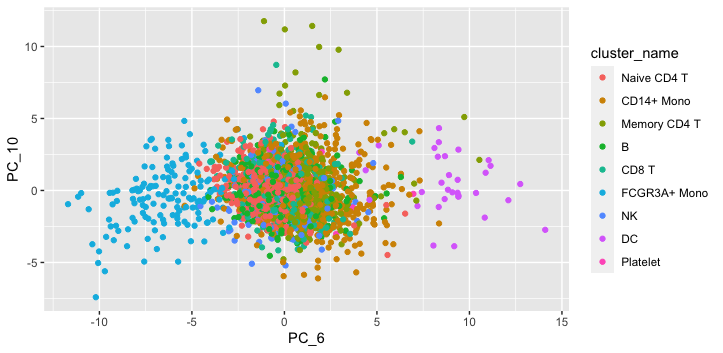

### 7_8.png

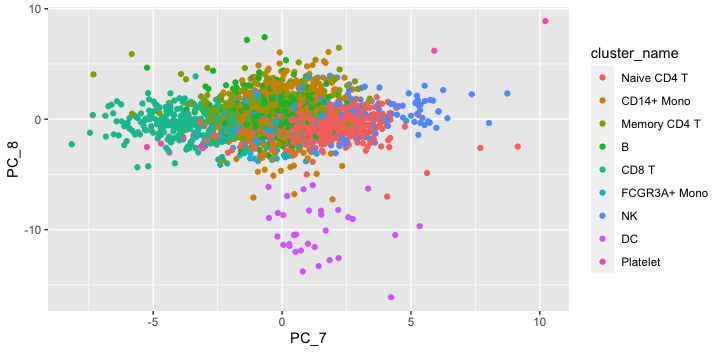

### 7_9.png

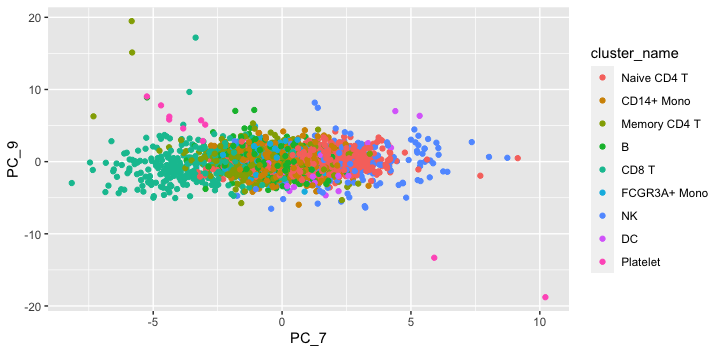

### 7_10.png

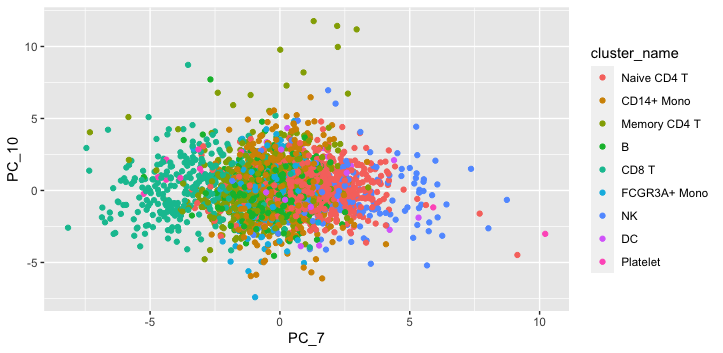

### 8_9.png

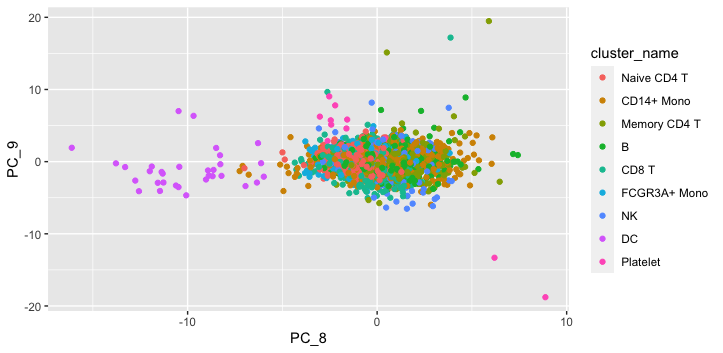

### 8_10.png

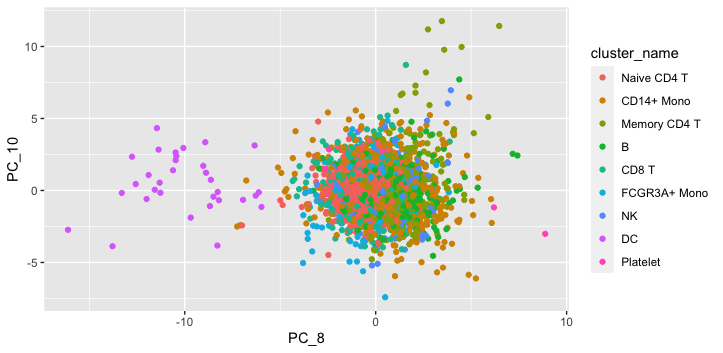

### 9_10.png

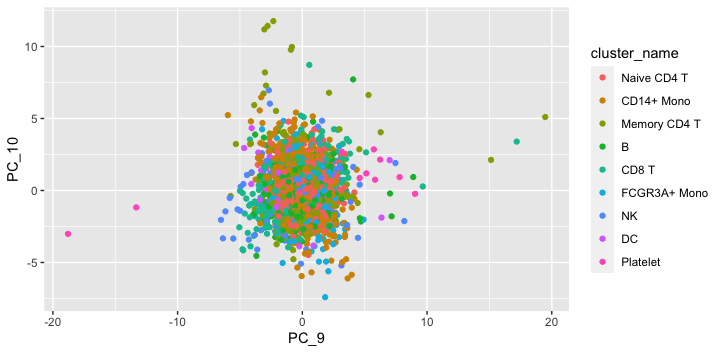
